# Chromatin priming and Hunchback recruitment integrate spatial and temporal cues in *Drosophila* neuroblasts

**DOI:** 10.1101/2025.11.25.690435

**Authors:** Ayanthi Bhattacharya, Hemalatha Rao, Sonia Q Sen

## Abstract

Neural stem cells generate diverse cell types by integrating spatial and temporal cues to activate neuron-specific terminal selector (TS) genes. In *Drosophila* neuroblasts (NBs), spatial patterning sets lineage identity, while a temporal transcription factor (TTF) cascade sets birth order. Two proposed mechanisms could integrate these inputs. In *direct regulation*, spatial transcription factors (STFs) and TTFs co-occupy and regulate TS enhancers within NBs. In *epigenetic regulation*, STFs first prime NB-specific chromatin, creating selected enhancers that can later recruit TTFs.

We tested these models in the NB5-6 and NB7-4 lineages using their candidate STFs, Gooseberry (Gsb) and Engrailed (En), together with the first TTF, Hunchback (Hb). We find that En preferentially occupies pre-accessible chromatin in the NB7-4 lineage, including highly accessible En–Hb co-bound sites. This is consistent with En acting within an already established chromatin landscape whose formation likely depends on additional NB7-4 factors. In contrast, Gsb binds both accessible and less-accessible chromatin in the NB5-6 lineage and can remodel accessibility bidirectionally when ectopically expressed in NB7-4 lineage, with corresponding changes in Hb occupancy. However, Gsb binding alone does not determine which sites become accessible or recruit Hb, indicating that productive Gsb–Hb regulatory states require additional NB5-6-specific inputs.

Thus, direct and epigenetic regulation are not alternative mechanisms, but distinct steps within the integration process. We therefore propose a third possibility: *distributed STF code model* in which these steps — chromatin priming and direct STF–Hb engagement — are distributed across the members of each NB-specific STF code. The STF code therefore shapes the enhancer landscape available to Hb and enables productive Hb engagement at lineage-appropriate enhancers.

## Introduction

The diversity of cell types in the nervous system arises from a small pool of neural stem cells (NSCs). Decades of work in *Drosophila* show that NSCs use a hierarchical regulatory logic: positional cues endow NSCs with distinct molecular identities and lineage potentials, while a temporal gene cascade determines the production of different neuron types over time. NSCs activate neuron-specific terminal selector (TS) genes by integrating these spatial and temporal inputs along with Notch signalling. TSs then drive effector gene programs that implement neuronal morphology, connectivity, and neurotransmitter identity (1–3).

This has been best elucidated in the embryonic ventral nerve cord (VNC) and the optic lobe. In the VNC, 30 neuroblasts (NBs) per hemisegment each generate a lineage with a unique identity that is determined at the time of delamination (4–8). This identity reflects the NB’s neuroectodermal origin, where intersecting anterior–posterior (A–P) and dorsoventral (D–V) patterning systems create distinct domains of TF expression (2, 3, 9) (Figure 1A). NBs inherit these spatial TFs, which specify lineage identity. For example, the A–P gene *gsb* is expressed in row 5 of the neuroectoderm and required for row-5 identity: its loss transforms NBs to a row 3/4 fate, while misexpression induces the converse transformation (10). Other STFs act non-autonomously: *wingless* (*wg*) shapes the identity of adjacent rows without directly specifying row 5 (11). More posteriorly, *en* defines rows 6 and 7: *en* mutants lack NB6-4 and NB7-3, while *en* misexpression duplicates NB7-3 identity (12–14). D–V patterning factors, including *vnd, ind*, and *msh*, similarly define medial, intermediate, and lateral NB columns. Gain- and loss-of-function studies demonstrate that these D–V genes also respecify NB columnar identity (15–18). Together, these intersecting spatial cues generate 30 distinct NB identities per hemisegment, each with a unique developmental potential.

**Fig. 1.**
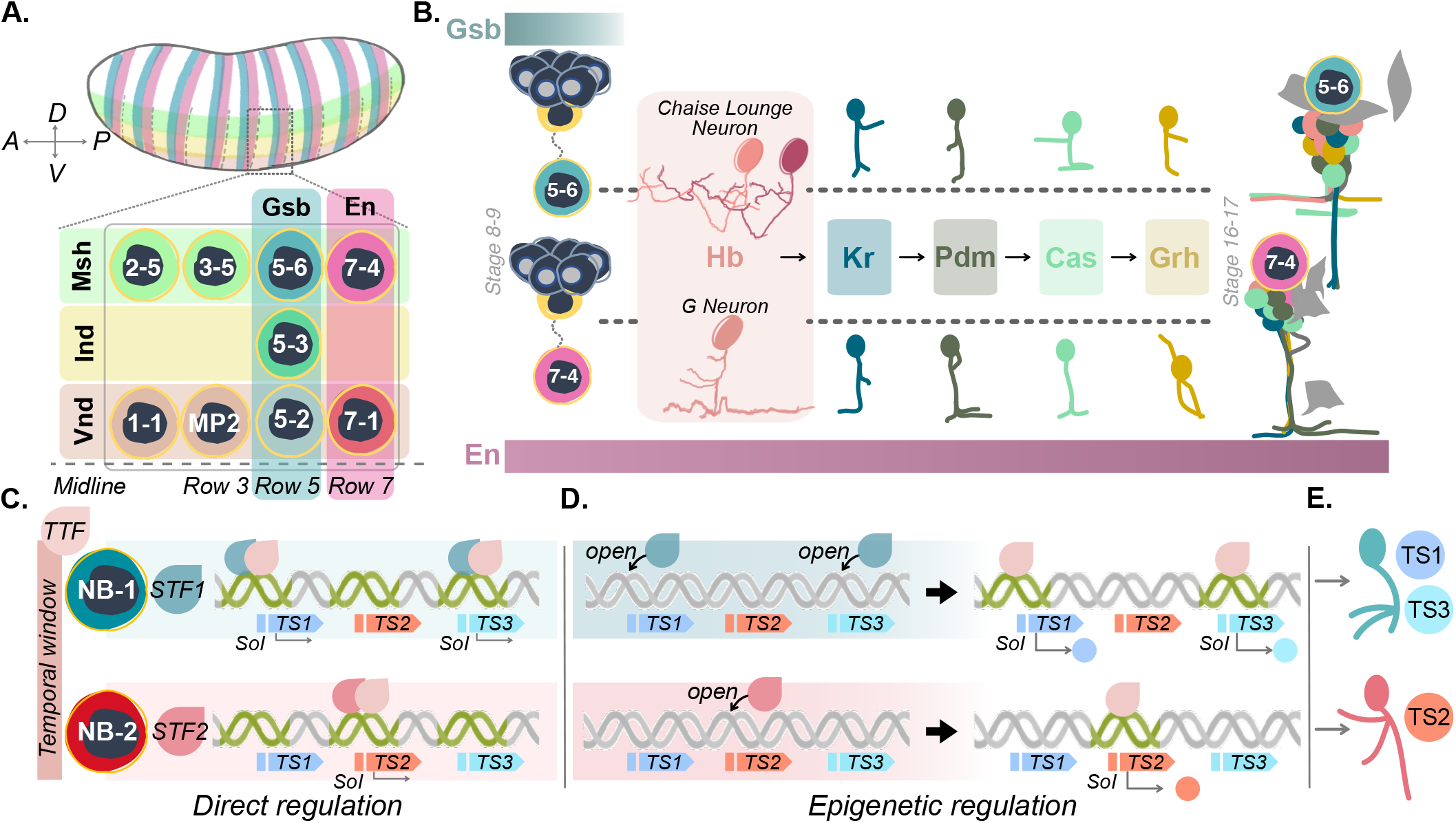
Spatial and temporal patterning of neuroblasts and models for their integration. **A. Spatial patterning:** Early embryo patterning genes that establish the A–P and D–V body axes, also pattern the neuroectoderm, located ventro-laterally in the embryo. The dashed box outlines one segmental repeat of this patterned neuroectoderm, with representative A–P gene expression shown in blue and pink, and D–V columns in light green, yellow and red. NBs delaminate from the neuroectoderm at stereotyped positions, each inheriting a unique molecular identity defined by its row (A–P) and column (D–V) coordinates. **B. Temporal patterning**. Each spatially distinct NB (here, NB5-6 and NB7-4 shown delaminating from neuroectodermal cell clusters) transitions through a series of temporal windows corresponding to sequentially expressed TTFs, Hb > Kr > Pdm > Cas > Grh. This allows NBs to generate different neurons over time. During the Hb temporal window (pink box), NB5-6 generates Chaise Lounge neurons while NB7-4 generates G neurons. By the end of the temporal cascade, each NB produces distinct lineages. STF expression relative to this cascade differs: Gsb expression in NB5-6 (blue bar) is transient and precedes the Hb window while En expression persists in NB7-4 (pink bar). **C–E. Models for spatiotemporal integration**. Two NBs (NB1, NB2) express different spatial transcription factors (STF1, STF2) but share the same TTF. In direct regulation (C), STFs and the TTF co-bind accessible enhancers of neuron type–specific terminal selector (TS) genes in each NB, requiring concurrent STF and TTF expression. In epigenetic regulation (D), STFs first bind and open chromatin in a NB-specific manner, creating sites of integration (SoIs) that are later bound by the TTF to activate TS enhancers; in this mode, STF expression may be transient. Distinct TS combinations (TS1/TS3 in NB1, TS2 in NB2) specify lineage-specific, time-appropriate neurons (E).

In the optic lobe, an analogous spatial logic partitions the neuroepithelium of the outer and inner proliferation centers (OPC/IPC) into A–P domains by Vsx, Optix, and Rx (split by Dpp and Wg and into D–V compartments by Spalt/Salm and Disco/Hh) (1, 19–23). These factors establish progenitor territories that seed distinct neuronal outputs across the medulla, lobula, and lobula plate (20–23). For example, Vsx-domain progenitors generate Pm3 neurons, dorsal and ventral Rx territories produce Pm1 and Pm2 respectively, whereas Optix-domain equivalents are eliminated by apoptosis (20). As in the VNC, manipulating spatial inputs reassigns territories and shifts lineage output. For example, expanding the Rx domain biases neural fates towards Pm1/2 output, and mutating *disco*, which specifies ventral fates, results in loss of ventral medulla neuron identities (20, 22).

NBs are also patterned in time. Each NB sequentially expresses a cascade of TTFs, creating successive competence windows during which different neurons are born (Figure 1B) (1, 2, 19, 24, 25). In the embryonic VNC, the canonical cascade proceeds through Hunchback (Hb) → Kruppel (Kr) → Pdm → Castor (Cas) → Grainyhead (Grh) (26–28). In the optic lobe, this cascade consists of 14 TTFs: Hth → Dmrt99B → Erm → Opa → L → Ey → Hbn → Scro → Slp1 → Slp2 → D → BarH1 → BarH2 →Tll (29–34). While the identity of the temporal factors varies across systems — including the central brain, type II lineages, and in the postembryonic VNC and brain — the strategy is conserved: sequential expression of TTFs subdivides neurogenesis into discrete temporal windows, each producing neuron types appropriate to that time point.

In keeping with this, manipulating temporal factors respecifies the temporal identity of the neurons. In the optic lobe, loss of function of Ey, Slp, or D stalls the temporal sequence and the neurons from those windows are lost; misexpressing any one of them biases production toward that window’s fates (31). For example, in the early Hth/Bsh window, *bsh* or *hth* loss eliminates Mi1 neurons, and Bsh overexpression induces Mi1-like neurons (31, 35, 36). Similarly, in the embryonic VNC, Hb, the first in the temporal cascade, is both necessary and sufficient for the earliest-born neurons in multiple NB lineages. Its misexpression prolongs early competence windows and induces supernumerary early-born neurons, while its loss abolishes these fates (26, 27, 37–40).

Importantly, the identity of a neuron from a given time window differs across lineages. For example, Hb+ neurons are: U1 in NB7-1, RP2 in NB4-2, EW1 in NB7-3, and CCAP in NB3-5-implying that temporal information alone is insufficient to specify neuronal identity (24). Instead, Hb – indeed any TTF – must act in a spatially defined context to specify neuronal fate.

The mechanism by which spatial and temporal cues are integrated to generate such context-dependent outcomes remains unclear. Two mechanistic models have been proposed (1, 41). In ***direct regulation***, STFs and TTFs act concurrently: both are present at the same time in the same NB, and co-occupy TS enhancers to drive activation. In ***epigenetic regulation***, the action is sequential: an STF first engages the enhancers to establish an accessible state, and only later does the TTF bind and activate transcription (Figure 1C–E).

Regardless of the model, we refer to the cis-regulatory loci that integrate the STF and TTF inputs as “sites of integration” (SoIs). Functionally, SoIs are enhancer elements that: (i) are specific to a particular NB lineage, and (ii) receive input from both the lineage-defining STF(s) and the TTF.

Here, we test these models using two well-characterized embryonic lineages — NB5-6 and NB7-4 — and their candidate STFs, Gooseberry (Gsb) and Engrailed (En), together with the first TTF, Hunchback (Hb). Gsb is expressed transiently in NB5-6 and precedes Hb (Figure 1B), suggesting that NB5-6 may rely on a chromatin-based memory of Gsb activity. En persists in NB7-4 during the Hb window (Figure 1B), so integration in this lineage could be epigenetic or direct. We therefore reasoned that the relationship between STF occupancy and chromatin accessibility could distinguish between these models. If integration is epigenetic, the lineage STF would be expected to engage chromatin independently of accessibility state to establish NB-specific SoIs — enhancers rendered competent for later TTF recruitment. If regulation is direct, the STF need not engage less-accessible chromatin; concurrent STF+TTF binding on pre-accessible enhancers suffices. Prior work suggested a role for Gsb in explaining lineage-specific Hb occupancy in NB5-6 (42). However, because Gsb occupancy had been inferred from whole-embryo ChIP-seq, the central prediction of the epigenetic model remained untested in NB5-6. We therefore profiled Gsb and En occupancy together with chromatin accessibility in the NB5-6 and NB7-4 lineages, asking whether either STF binds less-accessible chromatin and, if so, whether that binding can remodel accessibility and redirect Hb occupancy.

Our results indicate that in the NB7-4 lineage, En occupancy is strongly biased toward chromatin that is accessible during the Hb window, including NB7-4 SoIs. This is consistent with En acting with Hb at accessible enhancers, but does not by itself explain how the NB7-4 accessibility landscape is generated. In the NB5-6 lineage, Gsb binds both accessible and less-accessible chromatin, consistent with a role in chromatin priming; however, we find that Gsb does not act as a simple chromatin-opening factor. Instead, Gsb remodels chromatin at specific loci in both directions, with associated changes in Hb occupancy. These findings argue that neither candidate STF alone fully explains spatiotemporal integration in its lineage.

Together, these findings support a third possibility: a ***distributed STF-code model***. In this model, direct and epigenetic regulation are not alternate models, but separable molecular mechanisms distributed across members of a NB-specific STF code. Some code members shape the enhancer landscape available to Hb, while others bind with Hb at accessible SoIs to select and activate lineage-appropriate enhancers. This distributed code allows the same temporal factor to produce different transcriptional outputs in different lineages.

## Results

### Generation of functional Dam:Gsb and Dam:En to determine lineage-specific STF occupancy

To determine STF occupancy *in vivo* in a lineage-restricted manner, we used Targeted DamID (TaDa), which has been employed previously to measure the Hb TTF’s occupancy in neuroblasts (NBs) (42). We generated UAS-LT3-Dam:Gsb and UAS-LT3-Dam:En fly lines, in which an upstream mCherry ORF ensures low expression of Dam:Gsb or Dam:En from a downstream ORF (see Methods, Figure S1A). This configuration minimizes toxicity associated with high Dam levels (43). Expression of Dam alone (DamOnly), without a fused TF, methylates chromatin within accessible regions, providing a cell-type-specific readout of chromatin accessibility (CATaDa) (44).

We first confirmed that Dam:Gsb and Dam:En were not toxic to cells. Their ubiquitous expression under Da-Gal4 caused no embryonic lethality and no denticle belt defects. However, this lack of toxicity might reflect non-functional TFs when fused to Dam. We therefore tested the functionality of these fusion proteins. We performed whole-embryo Dam:Gsb profiling (Da-Gal4>UAS-LT3-Dam:Gsb) and compared it to published Gsb ChIP data (45). Three biological replicates were generated, alongside UAS-LT3-Dam controls (DamOnly), and processed as described (see Methods). Dam:Gsb and DamOnly replicates correlated strongly within their own sets but poorly with each other (Figure S1B), confirming reproducibility.

Dam:Gsb also reproducibly marked known Gsb targets (*wg, gsb-n, prd*) (Figure S1C), and signal enrichment was higher at Gsb-ChIP peaks than at ChIP peaks of other TFs (Figure S1D). Dam:Gsb peaks should be enriched for Gsb binding motifs. Although a canonical Gsb motif is not well-defined, Gsb contains both homeodomain and paired DNA-binding domains (46). A *de novo* motif search revealed enrichment of a paired-homeobox composite motif in Dam:Gsb peaks relative to background (Figure S1E). Similarly, Dam:En data were reproducible, showed occupancy at established En targets, and were enriched for the canonical En motif (Figure S1F–H).

Together, these experiments confirm that Dam:En and Dam:Gsb retain DNA-binding activity, are non-toxic to cells, and are reliable proxies for En and Gsb, providing validated tools to interrogate STF binding in the NB5-6 and NB7-4 lineages. Alongside the Dam:TF profiles, DamOnly signal serves as a proxy for chromatin accessibility in each lineage, allowing us to simultaneously profile STF occupancy and chromatin state within the same experimental framework.

### Assaying spatial TF occupancy in identified NB lineages

STFs such as Gsb and En are expressed broadly in the embryonic neuroectoderm, yet understanding their lineage-specific activities requires resolving their occupancy within individual NB lineages. This creates a major challenge for profiling their genomic occupancy: each lineage contains only a small number of cells during this early developmental window, and STF expression is often transient. Previous studies relied on whole-embryo ChIP data (47), which cannot resolve lineage-specific binding.

To overcome this limitation, we combined TaDa with lineage-specific Gal4 drivers to assay STF binding in two identified lineages: Dam:Gsb in NB5-6 and Dam:En in NB7-4 (Figure 2A–B). The driver lines were previously validated as being specific to these lineages (42). We collected data during stages 10–12, corresponding to the Hb competence period when SoIs are active. Despite sampling very few cells over this short time window, replicate Dam:En and Dam:Gsb profiles were highly reproducible and separated clearly from their DamOnly controls (Figure 2C–D). As in the whole-embryo data, these lineage-specific profiles marked known targets of En and Gsb, further validating these tools (Figure 2—figure supplement 1). In addition they also showed occupancy at lineage-associated marker genes (6) and at genes proximal to lineage-specific SoIs. For example, Dam:Gsb occupancy was observed at *gsb, lbl, lbe, eya, ap*, and *toy*, and Dam:En occupancy was observed at *en, acj6, unc-4*, and *d* (Figure 2E–F). These data suggest that these STFs may directly regulate loci linked to NB identity and terminal-selector programs.

**Fig. 2.**
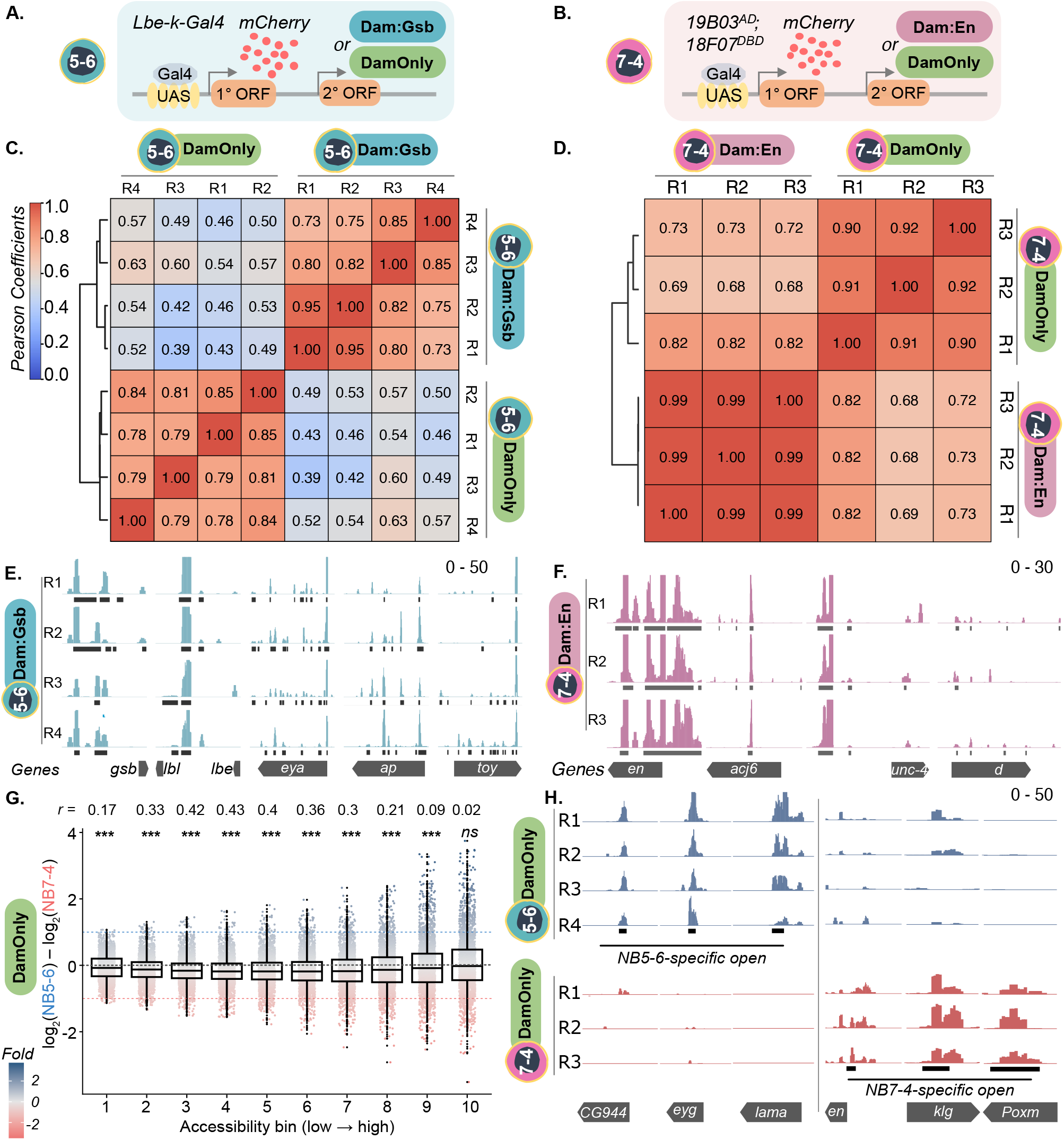
NB5-6 and NB7-4 TaDa profiling reveals STF occupancy and lineage-specific chromatin accessibility. **A–B. TaDa strategy for lineage-specific profiling.** In NB5-6 lineage, Lbe-k-Gal4 drives a UAS construct encoding mCherry in the primary ORF and Dam:Gsb (blue) or DamOnly (green) in the secondary ORF (A). In NB7-4 lineage, the split-Gal4 line 19B03[AD];18F07[DBD] similarly drives Dam:En (dark pink) or DamOnly (B). High mCherry expression reports driver specificity, while translation of Dam fusions from the secondary ORF maintains low Dam levels, minimising toxicity and preserving binding specificity. DamOnly profiles are used as a proxy for chromatin accessibility. **C. Reproducibility of NB5-6 DamOnly and Dam:Gsb profiles**. Pairwise Pearson correlation heatmap of NB5-6 DamOnly and Dam:Gsb replicates confirm dataset reproducibility and condition specificity. Hierarchical clustering separates the two conditions, with within-condition correlations higher than between-condition correlations. **D. Reproducibility of NB7-4 DamOnly and Dam:En profiles**. As in (C) for NB7-4 DamOnly and Dam:En replicates, confirming that NB7-4 profiles are reproducible and condition-specific. **E. Dam:Gsb occupancy at NB5-6-specific genes**. Genome browser snapshots show NB5-6 lineage Dam:Gsb (4 replicates) binding at NB5-6 marker genes *gsb, lbl, lbe, eya, ap and toy*. Data range: 0–50. **F. Dam:En occupancy at NB7-4-specific genes**. Genome browser snapshots show NB7-4 lineage Dam:En (3 replicates) binding at NB7-4 marker genes *en, acj6, unc-4, d*. Data range: 0–30. Black bars in (E) and (F) indicate peaks. **G. NB5-6 and NB7-4 show a broadly shared, but distinct chromatin landscape**. Chromatin accessibility (DamOnly signal) stratified into ten bins of increasing accessibility (see Methods, Figure 2—figure supplement 2). Accessibility bias, expressed as log_2_ fold-change in DamOnly signal from NB5-6 and NB7-4, is plotted across the ranked bins; positive values indicate NB5-6 bias, negative values NB7-4 bias, and values near 0 indicate shared accessibility. Each point represents one genomic window (*n* ≈ 2,700 per bin; RPGC-normalised, replicate-averaged), coloured by bias direction (blue, NB5-6-preferential; red, NB7-4-preferential; grey, shared). Boxes show median and interquartile range (IQR). Black dashed line marks zero bias. Blue and red dashed lines mark 2-fold threshold for NB5-6-preferential (log_2_ ≥1; 845 sites) and NB7-4-preferential loci (log_2_ ≤–1; 1067 sites), respectively. Across the spectrum, most regions show shared accessibility between both NBs, with bias distributed near zero. The spread of bias values indicates NB-specific differences at subsets of loci across all bins. Significance labels indicate that the median bias per bin deviates from zero (one-sample Wilcoxon signed-rank test, two-sided; ∗ ∗ ∗*p <* 0.001, ∗ ∗ *p <* 0.01, ∗*p <* 0.05, *ns* = not significant). Effect size r is shown above each bin. **H. Genome browser examples of differentially accessible loci**. Representative NB5-6-specific (blue) and NB7-4-specific (red) accessible sites are shown at the indicated loci. Data range 0–50. Genotypes for NB5-6-Gsb occupancy: *w-; Lbe-k-Gal4/+; +/UAS-Dam:Gsb*. NB5-6-chromatin accessibility: *w-; Lbe-k-Gal4/+; +/UAS-DamOnly*. NB7-4-En occupancy: *w-; 19B03[AD]/+; 18F07[DBD]/UAS-Dam:En*. NB7-4-chromatin accessibility: *w-; 19B03[AD]/+; 18F07[DBD]/UAS-DamOnly*.

We then used DamOnly profiles from the two NB lineages to investigate lineage-specific differences in chromatin accessibility. To do this, we first defined a common set of DamOnly-marked regions across NB5-6 and NB7-4 lineages and ordered these regions from least to most accessible. We then grouped them into ten bins spanning the full range of DamOnly signal, yielding 27,119 regions in total (see Methods, Figure 2—figure supplement 2A). Across these regions, DamOnly replicates separated by their lineage of origin, yet the two lineages showed broadly similar accessibility overall (Figure 2—figure supplement 2B). Since chromatin accessibility exists on a continuum rather than as binary open or closed states, this binned framework allowed us to compare lineage-specific accessibility differences across low, intermediate and high accessibility states Figure 2—figure supplement 2C–D). These data confirmed previous findings that NB5-6 and NB7-4 lineagews possess broadly shared but distinct chromatin landscapes, with lineage-specific accessibility differences present across the spectrum (Figure 2G–H). Together, these results show that TaDa can reliably profile Dam:Gsb and Dam:En occupancy in identified NB lineages. They also give us a way to use DamOnly signal to examine chromatin accessibility across the continuum of accessibility states in both lineages, allowing us to next ask whether Gsb and En differ in the kinds of chromatin they occupy.

### En preferentially binds pre-accessible chromatin in the NB7-4 lineage

The two models — ***direct*** or ***epigenetic* regulation** — make clear predictions for STF occupancy in the context of the chromatin state within NB lineages. While STF occupancy at less-accessible chromatin is necessary under the ***epigenetic regulation***, it is dispensable under ***direct regulation***.

To test these alternatives in NB7-4 lineage, we asked whether En occupancy is restricted to pre-accessible sites (consistent with conventional TF behaviour) or also engages less-accessible chromatin to establish SoIs (as predicted by epigenetic priming).

We first focussed our analyses on the previously generated SoIs (42). These sites were identified by performing a differential analysis on Hb binding in the NB5-6 and NB7-4 lineages, and were confirmed to be differentially accessible in the two lineages as previously reported (42) (see Methods, Figure 3A, Figure 3—figure supplement 1A–I). Annotating

**Fig. 3.**
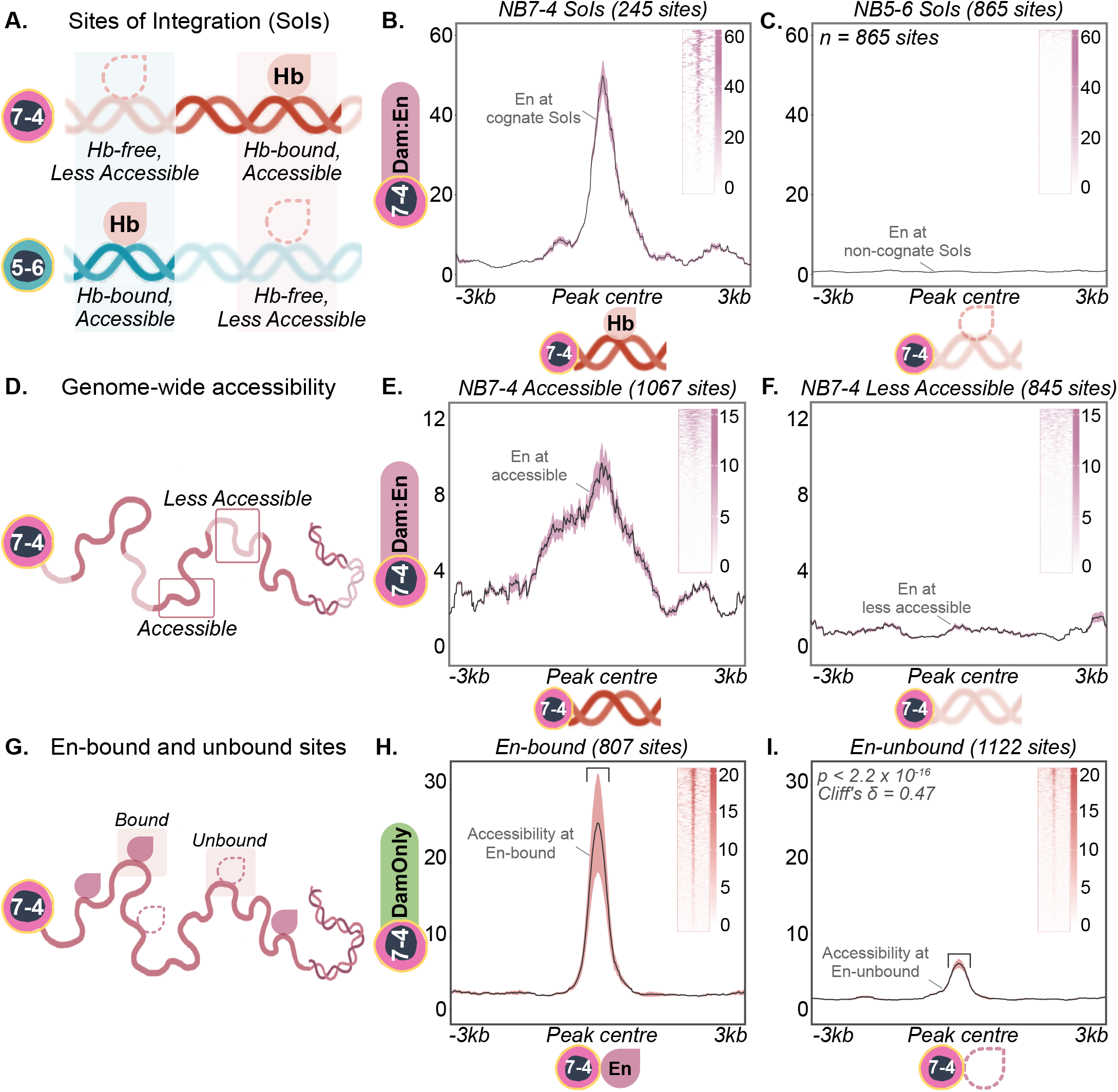
En preferentially binds pre-accessible chromatin. **A–C. En binding relative to Sites of Integration (SoIs).** Schematic (A) shows SoIs in each lineage, defined by differential Hb occupancy (FDR ≤ 0.01, FC ≥ 2; see Methods, Figure 3—figure supplement 1). NB7-4 SoIs (shaded pink box; 245 sites) are Hb-bound (filled drop) and accessible in NB7-4 (dark red strand), but Hb-free (hollow drop) and less accessible in NB5-6 (light blue strand). NB5-6 SoIs (shaded blue box; 865 sites) are Hb-bound (filled drop) and accessible in NB5-6 (dark blue strand), but Hb-free (hollow drop) and less accessible in NB7-4 (light red strand). In NB7-4 (A, top), En binds NB7-4 SoIs (B) but not NB5-6 SoIs (C). Plots show mean NB7-4-lineage Dam:En signal *±*SD (pink ribbon), *±*3kb from peak centres. Insets show per-site heatmaps of Dam:En signal at corresponding loci, averaged across three replicates. **D–F. En binding relative to genome-wide chromatin accessibility**. Schematic (D) shows accessible (1067 sites, dark red) and less accessible (845 sites, light red) loci in NB7-4 (see Methods, Figure 2G, Figure 3—figure supplement 2). En binds accessible sites (E), with little binding at less-accessible sites (F). Plots show mean NB7-4 Dam:En signal *±*SD (pink ribbon), *±*3kb from peak centres. Insets show per-site heatmaps of Dam:En signal at the corresponding loci, averaged across three replicates. **G–I. Chromatin accessibility relative to En-bound and En-unbound sites in NB7-4 lineage**. Schematic (G) shows En-bound (807 sites, dark pink drop) and En-unbound (1,122 sites, hollow drop) loci (see Methods, Figure 3—figure supplement 3). En-bound sites (H) are significantly more accessible than En-unbound sites (I), with signal compared *±*0.5kb around peak centres (square brackets; Mann–Whitney U, *p <* 2.2 *×* 10^−16^; Cliff’s *δ* = 0.47, medium effect). Plots show mean NB7-4 lineage DamOnly signal *±*SD (pink ribbon), *±*3kb from peak centres. Insets show per-site heatmaps of DamOnly signal at the corresponding loci, averaged across three replicates. Genotypes for NB7-4-Hb occupancy: *w-; 19B03[AD]/+; 18F07[DBD]/UAS-Dam:Hb*. NB5-6-Hb occupancy: *w-; Lbe-k-Gal4/+; +/UAS-Dam:Hb* (42). NB7-4-En occupancy: *w-; 19B03[AD]/+; 18F07[DBD]/UAS-Dam:En*. NB5-6-Gsb occupancy: *w-; Lbe-k-Gal4/+; +/UAS-Dam:Gsb*. NB7-4-chromatin accessibility: *w-; 19B03[AD]/+; 18F07[DBD]/UAS-DamOnly*. NB5-6-chromatin accessibility: *w-; Lbe-k-Gal4/+; +/UAS-DamOnly*.

TF-encoding genes near these sites recovered known lineage markers *lbl/lbe* for NB5-6 and *en* for NB7-4 suggesting that these sites are relevant for lineage identity (Figure 3—figure supplement 1J–K). As expected, En was enriched at NB7-4-specific SoIs, where Hb uniquely binds in this lineage (Figure 3B). However, there was no detectable En signal in NB7-4 at sites known to be NB5-6-specific SoIs (which remain relatively less accessible and Hb-free in NB7-4; Figure 3C). This difference is unlikely to be explained simply by En motif availability, since En motifs are similarly represented at NB7-4- and NB5-6-specific SoIs (Figure 3—figure supplement 1L). Thus, at SoIs, En occupancy is associated with the cognate, accessible NB7-4 SoIs, with little detectable occupancy at the non-cognate, relatively less accessible NB5-6 SoIs.

This SoI-level pattern was also reflected genome-wide, where we examined En occupancy relative to chromatin accessibility. For this comparison, we used the ranked accessibility bins defined above (Figure 2G). In general, En occupancy was found to be weighted towards higher accessibility bins (Figure 3—figure supplement 2A). Thresholding for sites that were at least two-fold differential between the two NBs yielded 1,067 loci preferentially accessible in NB7-4 lineage and 845 loci preferentially accessible in NB5-6 lineage, which are relatively less accessible in NB7-4 (see Methods, Figure 3D, Figure 3—figure supplement 2B–D). At these sites, En was strongly enriched at NB7-4 accessible loci (Figure 3E) and not at the less accessible ones (Figure 3F). Thus, consistent with the SoI-level analysis, En occupancy is associated primarily with chromatin that is accessible in the NB7-4 lineage.

To test this relationship more directly, we compared chromatin accessibility at En-bound versus En-unbound loci genome-wide. A differential analysis between En (NB7-4 lineage) and Gsb (NB5-6 lineage) yielded ‘En-bound’ sites in NB7-4, that were not Gsb-bound in NB5-6. Conversely, ‘En-unbound’ sites corresponded to those that were Gsb-bound in NB5-6, but not En-bound in NB7-4 (see Methods, Figure 3G, Figure 3—figure supplement 3A–D). En-bound regions displayed markedly higher accessibility than unbound ones (Figure 3H–I), suggesting that En-occupied sites tend to be associated with open chromatin. Consistent with this, ROC analysis, which asks how well chromatin accessibility separates En-bound from En-unbound sites across all possible thresholds, showed that accessibility predicts En occupancy better than chance (AUC = 0.74; Figure 3—figure supplement 3E). This pattern is consistent with En acting primarily at pre-accessible chromatin in NB7-4 lineage, rather than providing evidence that En itself establishes accessibility at these sites.

Taken together, these results show that En occupancy is strongly biased toward pre-accessible chromatin in NB7-4 lineage, including at NB7-4-specific SoIs. Thus, En is well placed to integrate spatial and temporal information through direct regulation at pre-accessible enhancers. However, because NB7-4 has a distinct accessibility landscape, the factors that establish this landscape remain unresolved. We therefore propose that NB7-4 may use a combinatorial STF-code logic, in which other lineage-specific factors help establish NB7-4-specific SoIs, while En occupies this pre-accessible landscape and likely collaborates with Hb to activate lineage-specific enhancers (tested below).

### Gsb binds both accessible and less-accessible chromatin in the NB5-6 lineage

Having established that En occupancy in the NB7-4 lineage is strongly biased toward pre-accessible chromatin, we next evaluated whether NB5-6 follows ***direct*** or ***epigenetic regulation*** by asking if Gsb, like En, is biased toward pre-accessible chromatin or instead engages both accessible and less-accessible regions to establish SoIs (epigenetic priming).

At NB5-6-specific SoIs, Gsb occupancy was enriched as expected (Figure 4A–B). However, Gsb signal in NB5-6 was also enriched at sites identified to be NB7-4-specific SoIs, which remain less accessible and Hb-free in NB5-6 lineage, indicating that Gsb binding alone does not determine where Hb will be recruited.(Figure 4C).

**Fig. 4.**
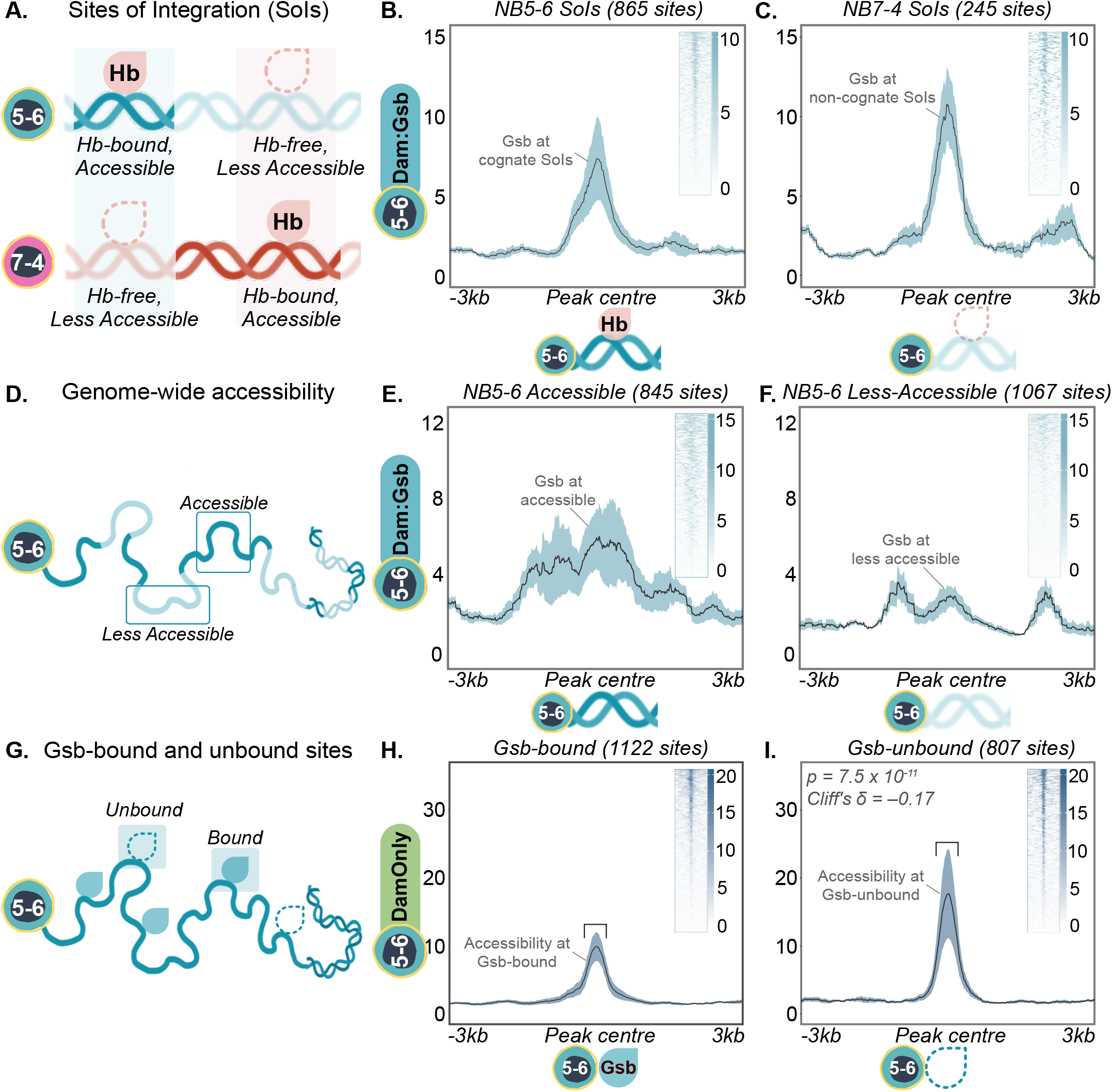
Gsb binds both accessible and less-accessible chromatin. **A–C. Gsb binding relative to Sites of Integration (SoIs).** Schematic (A) shows SoIs in each lineage, defined by differential Hb occupancy (FDR ≤ 0.01, FC ≥ 2; see Methods, Figure 3—figure supplement 1). NB5-6 SoIs (shaded blue box; 865 sites) are Hb-bound (filled drop) and accessible in NB5-6 (dark blue strand), but Hb-free (hollow drop) and less accessible in NB7-4 (light red strand). NB7-4 SoIs (shaded pink box; 245 sites) are Hb-bound (filled drop) and accessible in NB7-4 (dark red strand), but Hb-free (hollow drop) and less accessible in NB5-6 (light blue strand). In the NB5-6 lineage (A, top), Gsb binds both NB5-6 SoIs (B) and NB7-4 SoIs (C). Plots show mean NB5-6-lineage Dam:Gsb signal *±*SD (blue ribbon), *±*3kb from peak centres. Insets show per-site heatmaps of Dam:Gsb signal at corresponding loci, averaged across four replicates. **D–F. Gsb binding relative to genome-wide chromatin accessibility**. Schematic (D) shows accessible (845 sites, dark blue) and less-accessible (1067 sites, light blue) loci in the NB5-6 lineage (see Methods, Figure 2G, Figure 3—figure supplement 2). Gsb binds both accessible (E) and less-accessible (F) sites. Plots show mean NB5-6 Dam:Gsb signal *±*SD (blue ribbon), *±*3kb from peak centres. Insets show per-site heatmaps of Dam:Gsb signal at the corresponding loci, averaged across four replicates. **G–I. Chromatin accessibility relative to Gsb-bound and Gsb-unbound sites in the NB5-6 lineage**. Schematic (G) shows Gsb-bound (1,122 sites, filled blue drop) and Gsb-unbound (807 sites, hollow drop) loci (see Methods, Figure 3—figure supplement 3). Gsb-bound sites (H) are less accessible than Gsb-unbound sites (I), with signal compared *±*0.5kb around peak centres (square brackets; Mann–Whitney U, *p* = 7.5 *×* 10^−11^; Cliff’s *δ* = −0.17, small effect). Plots show mean NB5-6 lineage DamOnly signal *±*SD (blue ribbon), *±*3kb from peak centres. Insets show per-site heatmaps of DamOnly signal at the corresponding loci, averaged across four replicates. Genotypes for NB5-6-Hb occupancy: *w-; Lbe-k-Gal4/+; +/UAS-Dam:Hb*. NB7-4-Hb occupancy: *w-; 19B03[AD]/+; 18F07[DBD]/UAS-Dam:Hb* (42). NB5-6-Gsb occupancy: *w-; Lbe-k-Gal4/+; +/UAS-Dam:Gsb*. NB7-4-En occupancy: *w-; 19B03[AD]/+; 18F07[DBD]/UAS-Dam:En*. NB5-6-chromatin accessibility: *w-; Lbe-k-Gal4/+; +/UAS-DamOnly*. NB7-4-chromatin accessibility: *w-; 19B03[AD]/+; 18F07[DBD]/UAS-DamOnly*.

This suggested that Gsb has the ability to bind less-accessible chromatin. To test this genome-wide, we examined Gsb occupancy across the ranked accessibility bins. Gsb occupancy, like En, was biased towards higher accessibility bins, but some signal was also detected across the lower bins (Figure 3—figure supplement 2A). When examined in the context of the converse set of differentially accessible genomic loci, defined at the same two-fold cutoff as above (Figure 2G, Figure 3—figure supplement 2B–D), Gsb was enriched at the 845 loci preferentially accessible in the NB5-6 lineage as expected (Figure 4E). Importantly, we also detected some Gsb occupancy at the 1,067 loci that were relatively less accessible in the NB5-6 lineage (see Methods, Figure 4F). This distinguishes Gsb from En and is consistent with a capacity to engage chromatin across a broader range of accessibility states.

To test this relationship more directly, we compared chromatin accessibility at Gsb-bound versus unbound loci, which were determined as described earlier via differential analysis between Gsb (NB5-6 lineage) and En (NB7-4 lineage): enriched occupancy in the NB5-6 lineage yielded ‘Gsb-bound’ sites and ‘Gsb-unbound’ sites corresponded to those that were En-bound in the NB7-4 lineage, but not Gsb-bound in the NB5-6 lineage (see Methods, Figure 3—figure supplement 3A–D, Figure 4G). Genome-wide, accessibility was significantly lower at Gsb-bound regions relative to unbound ones (Figure 4H–I). An ROC analysis showed that, unlike En, accessibility failed to discriminate Gsb-bound from unbound sites better than chance (AUC = 0.41; Figure 3—figure supplement 3E), consistent with Gsb engaging both accessible and less-accessible chromatin.

Together, these results meet a key prediction of the epigenetic model: Gsb engages both accessible and relatively less-accessible chromatin. However, because many Gsb-bound sites, including NB7-4-specific SoIs, remain relatively less accessible and Hb-free in the NB5-6 lineage, Gsb binding is not sufficient to recruit Hb. Thus, as in NB7-4, a single STF does not fully explain lineage-specific SoI selection.

Rather, these results extend the combinatorial logic proposed above: Gsb provides a chromatin-engagement activity consistent with epigenetic priming, but productive Hb recruitment likely requires additional lineage-restricted factors that distinguish NB5-6 SoIs from non-cognate Gsb-bound sites.

### Co-binding with Hb marks highly accessible sites

Our analyses of Gsb in the NB5-6 lineage and En in the NB7-4 lineage suggest that chromatin engagement and Hb co-occupancy may represent separable steps in the proposed combinatorial STF-code logic. If productive spatiotemporal integration requires both lineage-specific STF engagement and TTF occupancy, then loci co-bound by Hb and an STF-code component should represent a highly accessible class of candidate regulatory sites.

To test this, we analyzed three genome-wide classes of loci — STF-only, Hb-only, and STF+Hb co-bound (Figure 5A– B) — and compared their chromatin accessibility. For each TF (Gsb in the NB5-6 lienage; En in the NB7-4 lineage; Hb in both), the top 5000 highest-confidence peaks covering known genomic targets were used. To define STF-only sites, we subtracted Hb-bound peaks that overlapped with STFs; the same approach was used to obtain Hb-only sites in each NB. Co-bound sites were defined as those with an overlap of at least 90%. In both NB5-6 and NB7-4, accessibility was lowest at Hb-only sites, intermediate at STF-only sites, and highest at STF+Hb co-bound loci (Figure 5C–H). This analysis shows that, in both lineages, STF+Hb co-occupancy marks the most accessible class among the tested loci. This supports the model proposed above in which chromatin engagement by lineage-specific STFs is not sufficient on its own; instead, productive integration is associated with Hb co-occupancy at the appropriate lineage-specific sites. Thus, STF+Hb co-binding marks a highly accessible class of candidate enhancers, while STF-only binding identifies sites that may remain non-productive without the appropriate temporal or lineage-restricted context.

**Fig. 5.**
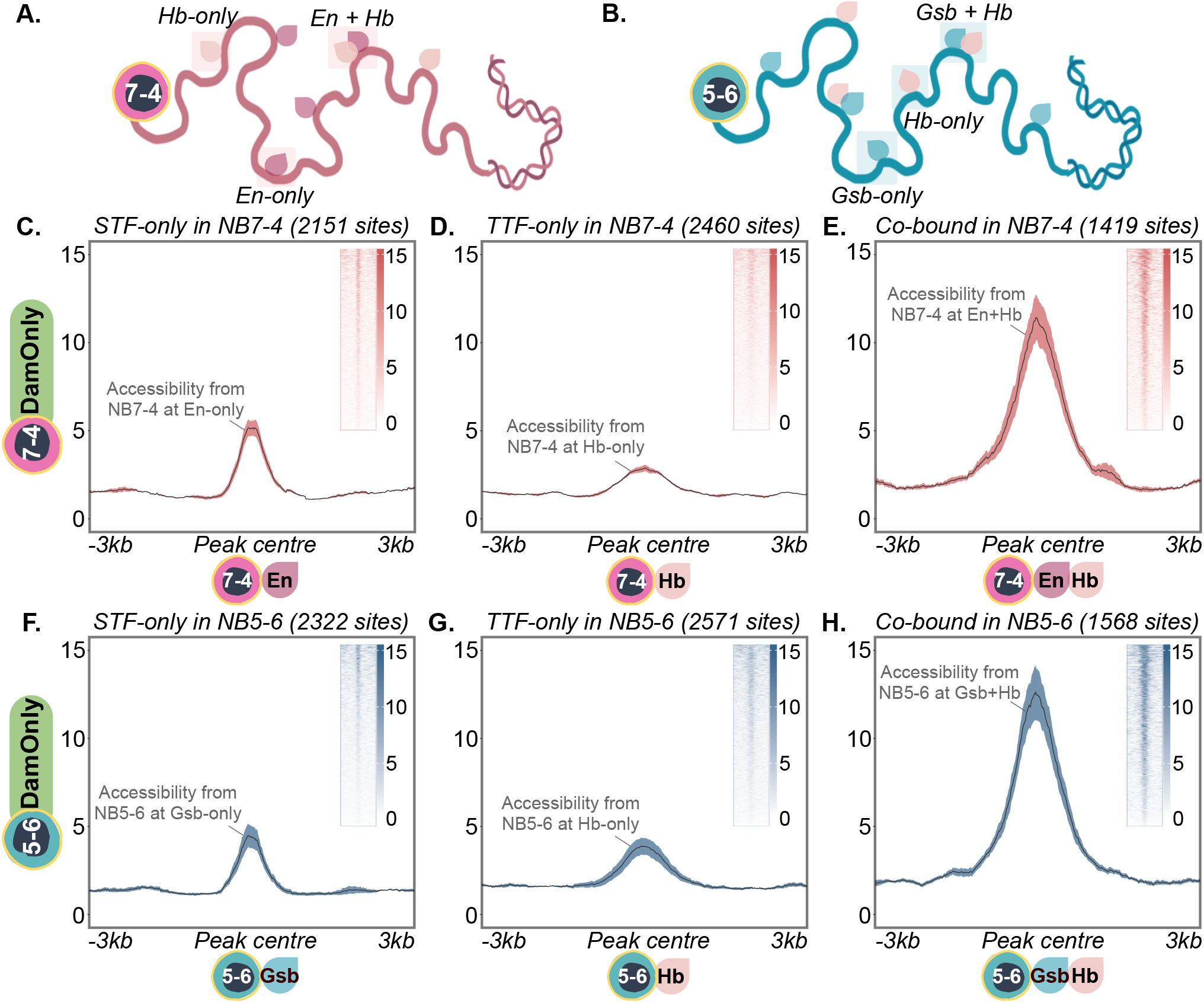
Co-binding with Hb marks sites of highest accessibility. **A–B. STF-only, Hb-only and STF+Hb co-bound sites.** Schematic (A) shows sites in the NB7-4 lineage bound by En only (dark pink drop; 2,151 sites), Hb only (light pink drop; 2,460 sites), or both (En+Hb; 1,419 sites). Schematic (B) shows sites in the NB5-6 lineage bound by Gsb only (blue drop; 2,322 sites), Hb only (light pink drop; 2,571 sites), or both (Gsb+Hb; 1,568 sites). STF-only and Hb-only sites were identified by excluding peaks that overlapped between the two datasets. Co-bound sites were defined by ≥90% peak overlap (see Methods). **C–E. Accessibility at NB7-4-lineage STF/Hb categories**. In the NB7-4 lineage, DamOnly signal is modest at En-only (C) and Hb-only (D) sites, and highest at En+Hb co-bound sites (E). **F–H. Accessibility at NB5-6 STF/Hb categories**. In the NB5-6 lineage, DamOnly signal is modest at Gsb-only (F) and Hb-only (G) sites, and highest at Gsb+Hb co-bound sites (H). Plots show mean DamOnly signal *±*SD (shaded ribbon; red, NB7-4; blue, NB5-6), *±*3kb from peak centres. Insets show per-site heatmaps of DamOnly signal at the corresponding loci, averaged across replicates (three for NB7-4, four for NB5-6). Genotypes for NB5-6-Hb occupancy: *w-; Lbe-k-Gal4/+; +/UAS-Dam:Hb*. NB7-4-Hb occupancy: *w-; 19B03[AD]/+; 18F07[DBD]/UAS-Dam:Hb* (42). NB5-6-Gsb occupancy: *w-; Lbe-k-Gal4/+; +/UAS-Dam:Gsb*. NB7-4-En occupancy: *w-; 19B03[AD]/+; 18F07[DBD]/UAS-Dam:En*. NB5-6-chromatin accessibility: *w-; Lbe-k-Gal4/+; +/UAS-DamOnly*. NB7-4-chromatin accessibility: *w-; 19B03[AD]/+; 18F07[DBD]/UAS-DamOnly*.

### Gsb remodels chromatin when ectopically expressed

Having established that Gsb in the NB5-6 lineage can bind both open and less-accessible chromatin, we next asked whether Gsb is sufficient to actively remodel chromatin accessibility. Binding to less-accessible loci is a hallmark of pioneers and chromatin-priming factors, but the simplest version of the epigenetic model makes a stronger prediction: that an STF should not only recognize compact chromatin but also open it to generate accessible sites competent for later TTF binding and activation. Addressing this prediction requires moving beyond correlative occupancy analyses to a functional test of whether Gsb can alter chromatin state when introduced into a heterologous lineage where it is not normally expressed, a standard approach for testing chromatin-remodelling capacity.

To this end, we ectopically expressed Gsb ubiquitously using a *hsGsb* construct during the Hb-competence window and assayed chromatin accessibility in the NB7-4 lineage, a lineage where Gsb is absent (see Methods, Figure 6A–B, Figure 6— figure supplement 1). This provided a heterologous context in which to assess whether Gsb could open less-accessible chromatin and create sites competent for Hb recruitment outside its endogenous NB5-6 setting. We found, however, that Gsb’s role in chromatin remodelling was more complex: ectopic Gsb remodelled chromatin bidirectionally with regions both losing and gaining accessibility upon Gsb misexpression. (Figure 6—figure supplement 2A–B, E). Thresholding for sites that showed at least two-fold change in accessibility yielded 1340 sites losing accessibility and 606 sites gaining accessibility (Figure 6C, F). These accessibility changes were accompanied by parallel changes in Hb occupancy: sites that lost accessibility showed reduced Hb occupancy, whereas sites that gained accessibility showed increased Hb occupancy (Figure 6D, G, Figure 6—figure supplement 2C– D, F).

**Fig. 6.**
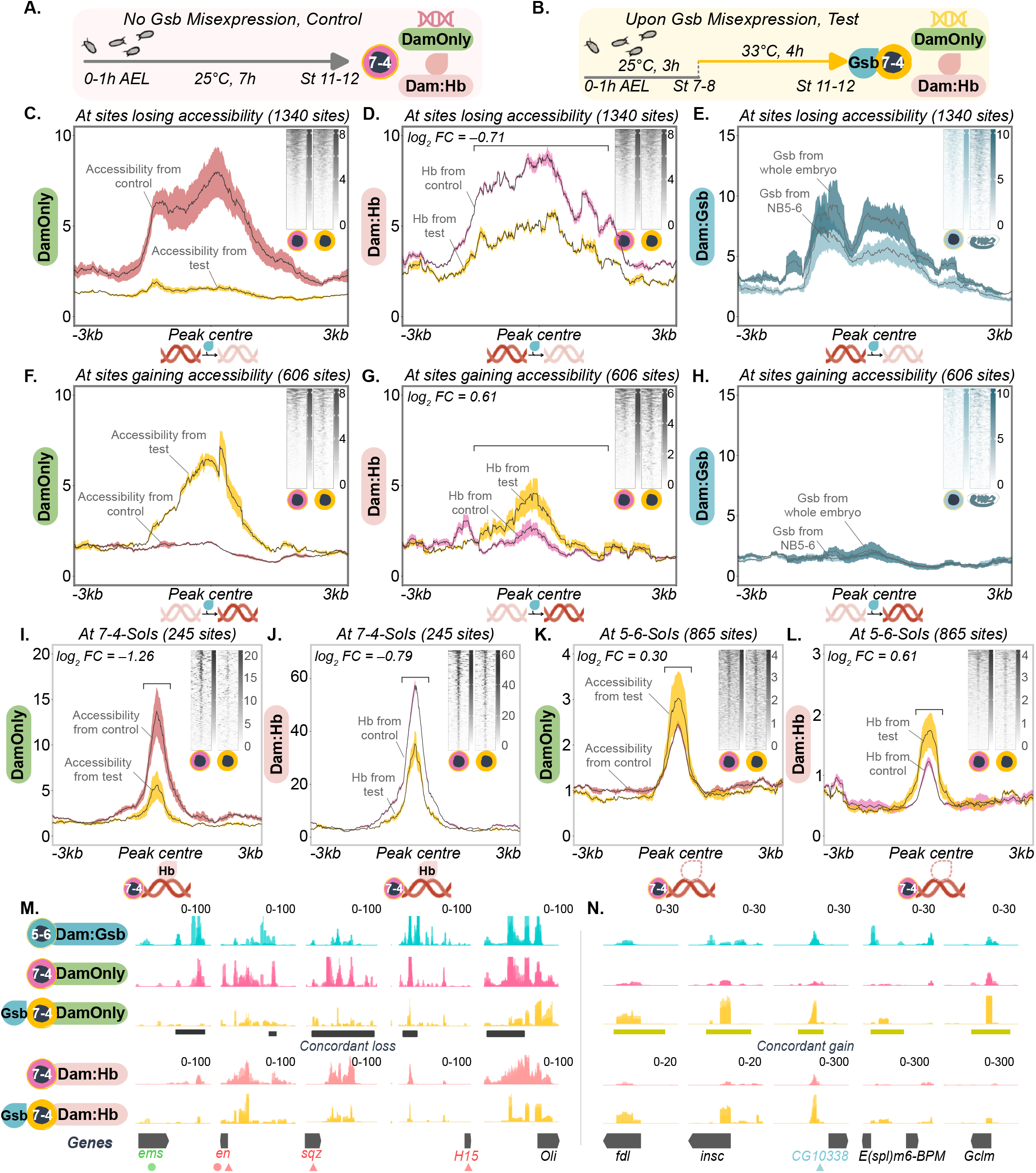
Gsb misexpression remodels chromatin accessibility and Hb occupancy at lineage-specific sites. **A–B. Experimental design.** Schematics show protocols for assaying chromatin accessibility (DamOnly) and Hb occupancy (Dam:Hb) in the NB7-4 lineage under control (A) and Gsb-misexpression (B) conditions. Control NB7-4 and its chromatin are shown in pink; Gsb-misexpressed NB7-4 and its chromatin in yellow. **C–E. Changes at sites losing accessibility**. At sites losing accessibility (1,340 sites), DamOnly signal is lower in test (yellow) than control (pink; C). Dam:Hb signal shows a corresponding loss from control (pink) to test (yellow; log_2_ FC = −0.71; D). Dam:Gsb occupancy at these sites is shown from two independent datasets: stage-matched NB5-6-lineage Gsb occupancy (stage 12, light blue) and cumulative whole-embryo Gsb occupancy (stage 17, dark blue; E). Loss sites were defined by the difference in log_2_-transformed DamOnly signal between test and control across the ranked accessibility bins, at a 2-fold cutoff (log_2_ FC ≤ −1; see Methods, Figure 6—figure supplement 2E). **F–H. Changes at sites gaining accessibility**. At sites gaining accessibility (606 sites), DamOnly signal is higher in test (yellow) than control (pink; F). Dam:Hb signal shows a corresponding gain from control (pink) to test (yellow; log_2_ FC = +0.61; G). Dam:Gsb occupancy at these sites is shown for the same two datasets as in (E; H); both Gsb datasets show weaker occupancy at gain sites than loss sites. Gain sites were defined at log_2_ FC ≥ +1 (see Methods, Figure 6—figure supplement 2E). Plots show mean DamOnly (C, F; three replicates each), Dam:Hb (D,G; three replicates each) or Dam:Gsb (E,H; four replicates from the NB5-6 lineage, three from whole-embryo) signal; *±*SD (shaded ribbon), *±*3kb from peak centres. Insets show per-site heatmaps, averaged across replicates, with rows ordered by descending signal in control. I–J. NB7-4 SoIs lose accessibility and Hb occupancy upon Gsb misexpression. Across NB7-4 SoIs (245 sites; accessible and Hb-bound in control), accessibility (DamOnly, I) and Hb occupancy (Dam:Hb, J) show a concordant loss (DamOnly log_2_ FC = −1.26; Dam:Hb log_2_ FC = −0.79, control to test; replicates non-overlapping in signal range). **K–L. NB5-6 SoIs gain accessibility and Hb occupancy upon Gsb misexpression**. Across NB5-6 SoIs (865 sites; less accessible and low Hb occupancy in control), accessibility (DamOnly, K) and Hb occupancy (Dam:Hb, L) show a small concordant gain (DamOnly log_2_ FC = 0.30; Dam:Hb log_2_ FC = 0.61, control to test; replicates non-overlapping in signal range). For I–L, plots show mean DamOnly (I, K) or Dam:Hb (J, L) signal *±*SD (shaded ribbon), *±*3kb from peak centres; log_2_ FC is computed from the change in signal from control to test, *±*0.5kb around peak centres. Insets show per-site heatmaps, averaged across replicates, with rows ordered by descending signal in control. **M–N. Genome browser examples of sites showing concordant changes in accessibility and Hb occupancy**. Representative sites undergoing concordant loss (M) and concordant gain (N) in accessibility and Hb occupancy. Triangles mark remodelled loci that are also SoIs (red, NB7-4 SoIs, M; blue, NB5-6 SoIs, N); circles mark remodelled loci that are also NB-specific markers (green, NB3-5; red, NB7-4). Tracks show, from top: NB5-6 Dam:Gsb occupancy; control and test NB7-4-lineage DamOnly signal; and control and test NB7-4-lineage Dam:Hb occupancy. Dark grey bars mark concordant-loss peaks (M); green bars mark concordant-gain peaks (N). Data range as indicated. Genotypes for NB5-6-Hb occupancy: *w-; Lbe-k-Gal4/+; +/UAS-Dam:Hb*. NB7-4-Hb occupancy: *w-; 19B03[AD]/+; 18F07[DBD]/UAS-Dam:Hb* (42). NB5-6-Gsb occupancy: *w-; Lbe-k-Gal4/+; +/UAS-Dam:Gsb*. NB5-6-chromatin accessibility: *w-; Lbe-k-Gal4/+; +/UAS-DamOnly*. NB7-4-En occupancy: *w-; 19B03[AD]/+; 18F07[DBD]/UAS-Dam:En*. NB7-4-chromatin accessibility: *w-; 19B03[AD]/+; 18F07[DBD]/UAS-DamOnly*. NB7-4-chromatin accessibility with Gsb misexpression: *hsGsb; 19B03[AD]/+; 18F07[DBD]/UAS-DamOnly*. NB7-4-Hb occupancy with Gsb misexpression: *hsGsb; 19B03[AD]/+; 18F07[DBD]/UAS-Dam:Hb*.

Although we did not directly profile Gsb occupancy in the NB7-4 lineage upon misexpression, we asked whether the sites that lose or gain accessibility differ in their propensity to be bound by Gsb. To do this, we mapped independent Gsb occupancy datasets onto these two classes of remodelled sites: stage-matched Dam:Gsb from the NB5-6 lineage, where Gsb is normally expressed, and whole-embryo Dam:Gsb, which provides a broader map of Gsb-compatible loci. In both datasets, sites that lost accessibility upon Gsb misexpression in the NB7-4 lienage showed strong Gsb occupancy, whereas sites that gained accessibility showed little or no Gsb signal (Figure 6E, H). Thus, the most direct consequence of ectopic Gsb is likely accessibility loss at loci that Gsb can bind, and this loss is accompanied by reduced Hb occupancy. By contrast, accessibility gains occur mainly at loci with little evidence of Gsb binding in independent datasets, suggesting that these gains are more likely indirect consequences of Gsb misexpression.

We next focussed our analysis on established SoIs in the NB7-4 lineage and asked how ectopic Gsb affects them. At endogenous NB7-4 SoIs, where chromatin is normally highly accessible and strongly bound by Hb in the NB7-4 lineage, ectopic Gsb led to a marked reduction in accessibility (Figure 6I). This was accompanied by a corresponding reduction in Hb occupancy at the same sites (Figure 6J). Thus, Gsb-mediated closing at native NB7-4 SoIs is coupled to dampened Hb recruitment. We then asked whether Gsb could conversely open the non-cognate NB5-6 SoIs in the NB7-4 lineage. At these loci, which are normally less accessible and Hb-free in the NB7-4 lineage, Gsb misexpression produced a modest increase in accessibility and Hb occupancy (Figure 6K–L). However, this gain was much weaker than the loss observed at endogenous NB7-4 SoIs. The SoI-level analysis therefore mirrors the genome-wide pattern: ectopic Gsb is most strongly associated with chromatin closing and reduced Hb occupancy, while its ability to open non-cognate sites and recruit Hb is limited. Genome browser examples of remodelled loci showing concordant loss or gain in accessibility and Hb occupancy are shown in Figure 6M–N.

Together, these experiments demonstrate that Gsb is sufficient to remodel chromatin when misexpressed, establishing its chromatin-remodelling capacity *in vivo*. In the heterologous NB7-4 lineage context, Gsb-associated remodelling is biased toward reducing accessibility, and this loss of accessibility is accompanied by reduced Hb occupancy both genome-wide and at endogenous NB7-4 SoIs. By contrast, ectopic Gsb produces only modest accessibility and Hb gains at non-cognate NB5-6 SoIs. Thus, Gsb’s chromatin-remodelling capacity is not equivalent to autonomous opening of lineage-appropriate enhancers. Rather, these data support a model in which Gsb contributes to chromatin priming as part of a broader NB-specific STF code, with additional lineage-specific inputs required for robust opening and Hb recruitment at cognate NB5-6 SoIs.

### Manipulating Gsb alters NB lineage identity

Having shown that ectopic Gsb remodels chromatin and alters Hb occupancy in the NB7-4 lineage, we next asked whether altering Gsb function changes NB lineage identity. The mis-expression experiments described above had already shown that, when induced at high levels, Gsb misexpression turned off the NB7-4-lineage Gal4 reporter, consistent with Gsb suppressing NB7-4 identity at sufficient levels. In the chromatin-profiling experiments above, this was addressed by inducing lower Gsb misexpression levels that preserved Gal4 expression (Figure 6—figure supplement 1). Together, these observations suggested that Gsb misexpression can influence lineage identity as well as chromatin state. However, since NB7-4 is not an endogenous Gsb-expressing context, we turned to NB5-6 to test the developmental consequences of altering Gsb function.

We first tested whether Gsb is necessary for NB5-6 identity by depleting Gsb in the early embryo using Vasa-Cas9, which is expressed in somatic cells of the early embryo (48), together with a *gsb*-targeted guideRNA. Gsb-depleted embryos were embryonic lethal, showed axonal defects consistent with previously described *gsb* loss, and sequencing confirmed a high mosaic editing rate (Figure 7—figure supplement 1).

**Fig. 7.**
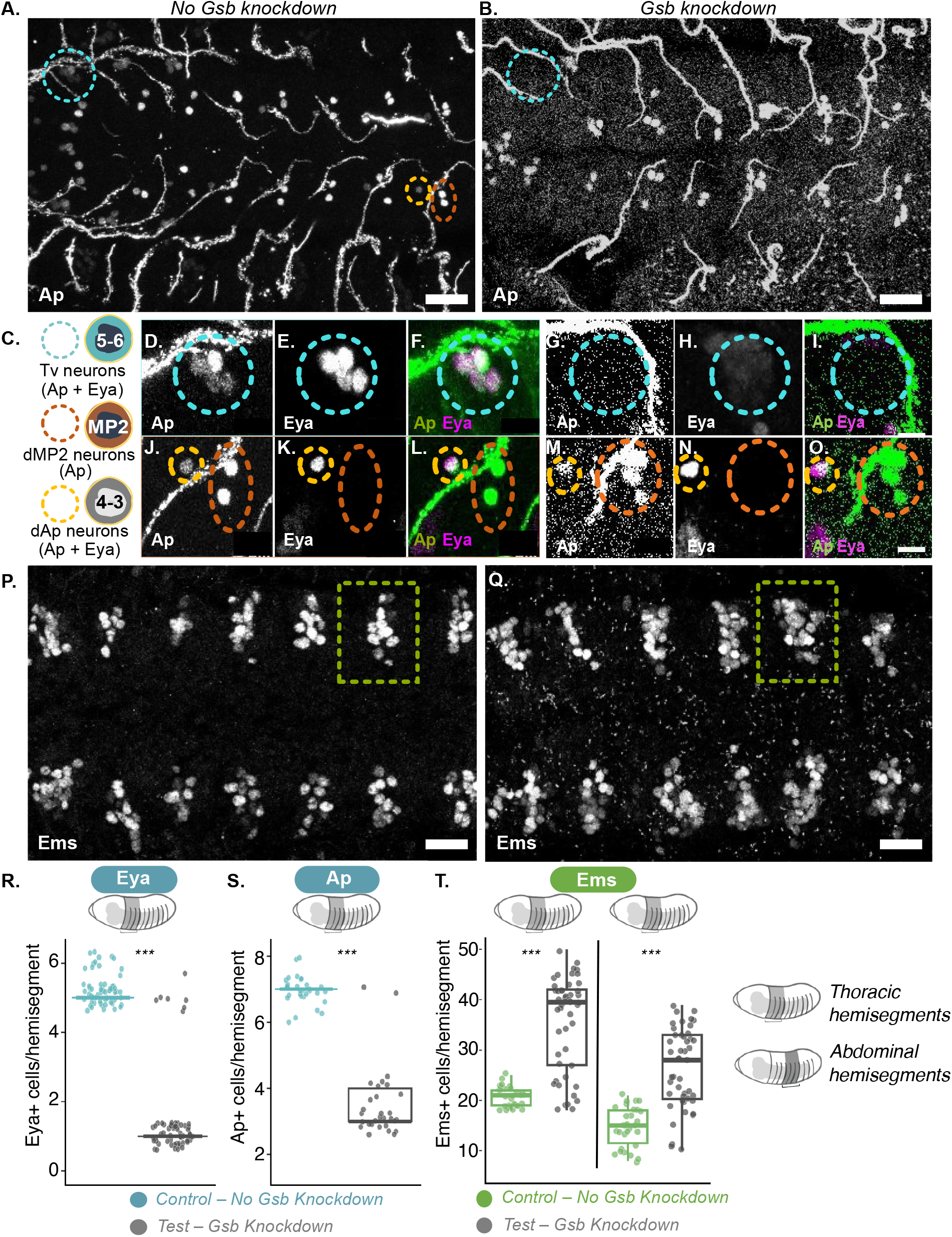
Broad loss of Gsb function results in loss of NB5-6 identity and gain of NB3-5 identity. **A–B. Gsb knockdown results in loss of Ap-positive Tv neurons born from NB5-6.** Control embryos show Ap-positive neurons born from NB5-6 (cyan circle), dMP2 neurons from MP2 (orange circle), and dAp neurons from NB4-3 (yellow circle) (A). Test embryos with Gsb knockdown show loss of Ap-positive Tv neurons from NB5-6 (cyan circle) (B). Scale bar 10 *µ*m. **C. Schematic of Ap-positive neuron populations**. Tv neurons (Ap+Eya) are born from NB5-6; dMP2 neurons (Ap-positive, Eya-negative) are born from MP2; dAp neurons (Ap+Eya) are born from NB4-3. **D–I. Tv neurons are lost upon Gsb knockdown**. Control embryos show a cluster of four Ap-positive Tv neurons per thoracic hemisegment (D), co-expressing Eya (E); merge shown in (F). Upon Gsb knockdown, Tv neurons are absent (cyan circle; G–I). **J–O. Gsb knockdown results in a concomitant gain in dMP2-like neurons**. Control embryos show one dMP2 neuron per hemisegment (orange circle), expressing Ap (J) but not Eya (K); merge shown in (L). dAp neurons (yellow circle) co-express Ap and Eya in control (J–L). Upon Gsb knockdown, dAp neurons remain unaffected (yellow circle; M–O), but dMP2 neurons show an increase — two pairs of Ap-positive neurons are present instead of one (orange circle; M–O). Scale bar 5 *µ*m. **P–Q. Gsb knockdown results in gain of Ems expression**. Control embryos show baseline Ems expression (P). Upon Gsb knockdown, Ems expression is increased (Q; green dashed box indicates example hemisegment). Scale bar 10 *µ*m. **R–T. Quantification of lineage markers under control and test conditions**. Each point represents a single hemisegment; boxplots show median and IQR. Quantification of Eya (R) and Ap (S) is shown for thoracic hemisegments; Ems (T) is shown for thoracic and abdominal hemisegments. Statistical significance was assessed using two-tailed Mann–Whitney U tests (∗ ∗ ∗*p <* 0.001, ∗ ∗ *p <* 0.01, ∗*p <* 0.05, *ns* = not significant). Genotype for control embryos: *Vasa Cas9/+;;*. For test embryos: *Vasa Cas9/+; gsb-guideRNA/+*.

We then asked how Gsb loss affects NB5-6 identity. In wild-type embryos, NB5-6-born Tv neurons are identified by co-expression of Ap and Eya (49) (Figure 7A, C–F). In Gsb-depleted embryos, these markers, along with the NB5-6-specific Lbe-K enhancer reporter, were lost, demonstrating that Gsb is required for NB5-6 identity (Figure 7B, G– I). Ap also labels neurons outside the NB5-6 lineage, providing an internal readout for effects on neighbouring lineages. Ap-positive neurons are also born from MP2 in row 3 and from NB4-3, named dMP2 and dAp neurons, respectively (50) (Figure 7A, C, J–L). These populations responded differently to Gsb loss: dMP2 neurons were approximately doubled in several hemisegments, whereas dAp neuron numbers were unaffected (Figure 7M–O). Thus, Gsb loss did not cause a general disruption of Ap-positive neurons, but instead produced a biased gain of a row-3-associated neuronal population. Consistent with this, Gsb-depleted embryos also showed ectopic expression of Ems, a row-3/NB3-5-associated marker (Figure 7P–Q).

Quantification across hemisegments confirmed this shift. Because NB5-6-specific Eya- and Ap-positive cells arise only in thoracic hemisegments, we limited quantification of these markers to thoracic hemisegments. These analyses confirmed loss of NB5-6 markers and gain of row-3/NB3-5-associated markers (Figure 7R–T), demonstrating that Gsb is required for NB5-6 identity and for suppression of row-3-associated fates. These findings are consistent with earlier reports that Gsb specifies row 5 and suppresses row 3 fates (10).

Having found that Gsb loss removes NB5-6 markers and permits row-3/NB3-5-associated markers, we next asked whether Gsb gain drives the opposite shift. We used the *hs-Gsb* misexpression system to increase Gsb levels at the time of NB specification, and assayed NB5-6 markers together with row-3/NB3-5 markers (Figure 8A). Based on our Gsb loss-of-function data, we expected that Gsb gain of function might produce the reciprocal phenotype: an increase in NB5-6 identity, possibly at the expense of row-3 identity.

**Fig. 8.**
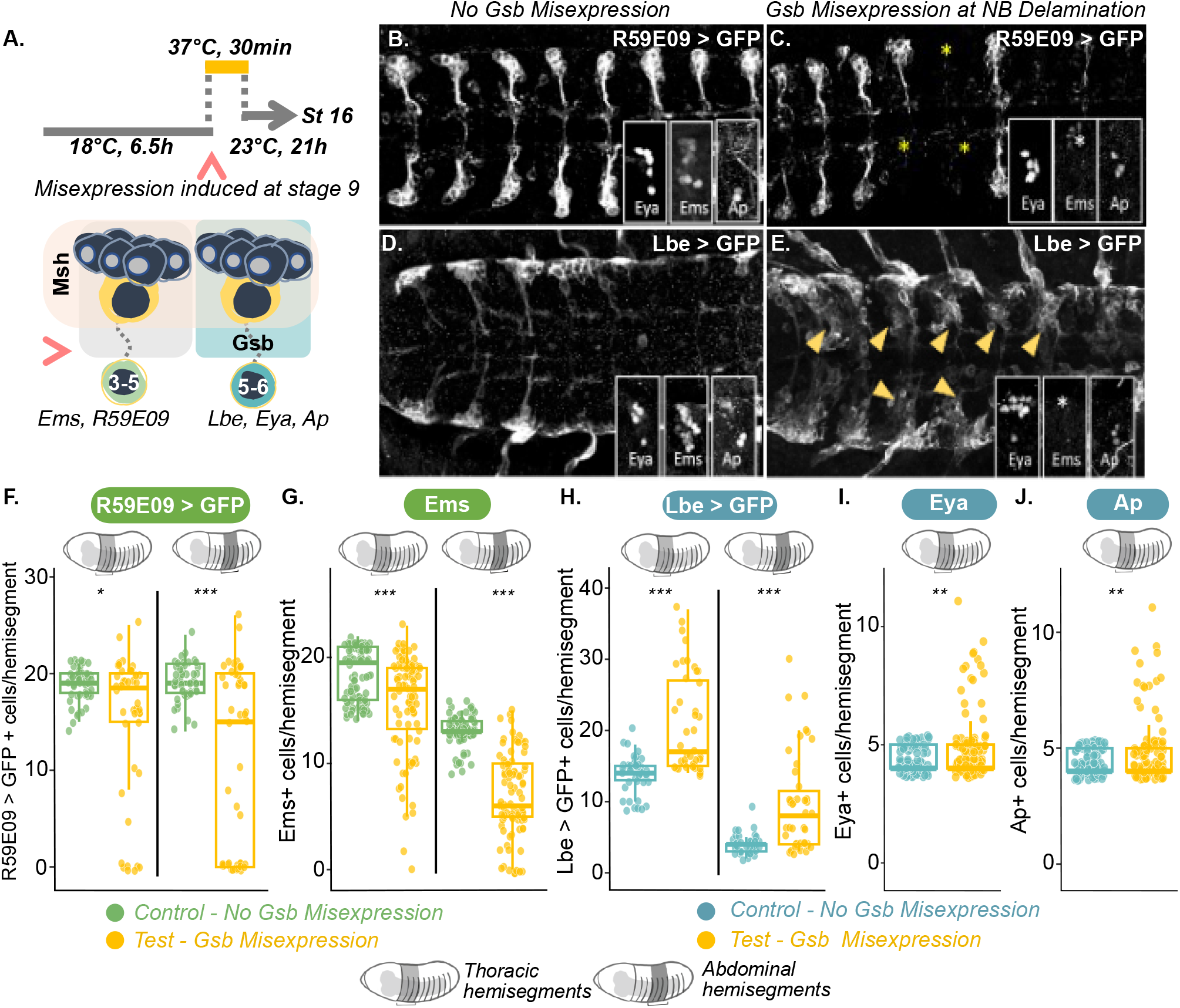
Broad misexpression of Gsb results in loss of NB3-5 identity, but partial gain of NB5-6 identity. **A. Experimental protocol used to induce broad Gsb misexpression.** (Top) Schematic represents heat shock protocol used to induce broad Gsb misexpression at NB delamination (red arrowhead). (Bottom) Both NB3-5 and NB5-6 arise in the Msh domain; however, only NB5-6 normally expresses Gsb. NB3-5 identity is assessed using R59E09>GFP and Ems; NB5-6 identity is assessed using Lbe>GFP, Eya, and Ap. **B–C. Gsb misexpression results in loss of NB3-5 markers**. Control embryos with NB3-5 lineages labelled by R59E09>GFP (B). Insets show wild-type expression of Eya, Ems and Ap. Test embryos show missing NB3-5 lineages (yellow asterisks) and a concomitant loss of Ems (white asterisk); expression of Eya and Ap remains largely unaffected (insets). **D–E. Broad Gsb misexpression results in partial gain of NB5-6 markers**. Control embryos with NB5-6 lineage labelled by Lbe>GFP (D). Insets show wild-type expression of Eya, Ems and Ap. Test embryos with broad Gsb misexpression show ectopic Lbe>GFP expression (E, yellow arrowheads). Insets show a concomitant gain of Eya, but not Ap, and loss of Ems. Loss of NB3-5 identity is not always accompanied by a gain in NB5-6 identity. **F–J. Quantification of lineage markers under control vs test conditions**. Each point represents a single hemisegment; boxplots show median and IQR. Quantification of GFP, Ems, Eya and Ap is shown separately in thoracic as well as abdominal segments. Among thoracic hemisegments, 27% showed loss of NB3-5 lineages with concomitant loss of Ems, and 4% additionally showed gain of Eya (n = 48); 38% of abdominal hemisegments showed loss of NB3-5 lineages with loss of Ems (n = 48). Statistical significance was assessed using two-tailed Mann–Whitney U tests (∗ ∗ ∗*p <* 0.001, ∗ ∗ *p <* 0.01, ∗*p <* 0.05, *ns* = not significant). Genotypes for both control embryos (no heat shock) and test embryos (with heat shock): *hsGsb/w-; Lbe-k-Gal4, UAS-mCD8-GFP* and *hsGsb/w-;;R59E09,UAS-myrGFP*.

Indeed, broad Gsb misexpression produced a partial gain of NB5-6 identity. In several thoracic hemisegments, we observed ectopic Lbe>GFP and supernumerary Eya- and Ap-positive neurons, consistent with a gain of NB5-6-born Tv-like neurons (Figure 8D–E). Concomitantly, it also suppressed NB3-5 identity: a proportion of hemisegments showed complete loss of the NB3-5 reporter R59E09>GFP and Ems, two independent markers of NB3-5 identity (Figure 8B–C). Since Ems also marks NB3-3, another row-3 NB, complete loss of Ems from an affected hemisegment suggests a broader disruption of row-3 identity following ectopic Gsb expression, consistent with previous findings (10). This loss of Ems expression also provides a locus-level link to the chromatin data above: in the heterologous NB7-4 lineage assay, Gsb misexpression resulted in loss of chromatin accessibility and Hb occupancy at the *ems* locus (Figure 6M).

Together, the gain- and loss-of-marker phenotypes indicate a reciprocal but incomplete shift in lineage identity. Only a small proportion of hemisegments showed both gain of NB5-6 identity and concomitant loss of NB3-5 identity. Quantification confirmed that while Gsb is sufficient to repress row-3/NB3-5-associated identity, it is not sufficient on its own to reliably impose the full NB5-6 lineage program (Figure 8F– J).

This incomplete reciprocity is important: it argues against Gsb acting as a stand-alone determinant of NB5-6 fate. Instead, it supports a combinatorial STF-code model in which Gsb contributes to chromatin remodelling and row-identity bias, while complete lineage specification requires additional spatial inputs and temporal factors such as Hb.

## Discussion

Two models have been proposed to explain how neural stem cells integrate spatial and temporal information to produce lineage-appropriate, time-specific neurons. In the direct model, STFs and TTFs together bind and regulate terminal-selector gene enhancers to drive lineage-specific effector programs. In the epigenetic model, STFs first establish a lineage-specific chromatin landscape that later permits TTF binding and enhancer activation. Our results instead support a third possibility: a ***distributed STF-code*** model in which direct and epigenetic activities are partitioned across different members of a neuroblast-specific STF code. In this model, the TFs that constitute a NB’s code contribute to shaping that NB’s chromatin landscape: opening lineage-appropriate loci, restricting inappropriate loci, and enabling productive TTF recruitment (Figure 9A–B).

**Fig. 9.**
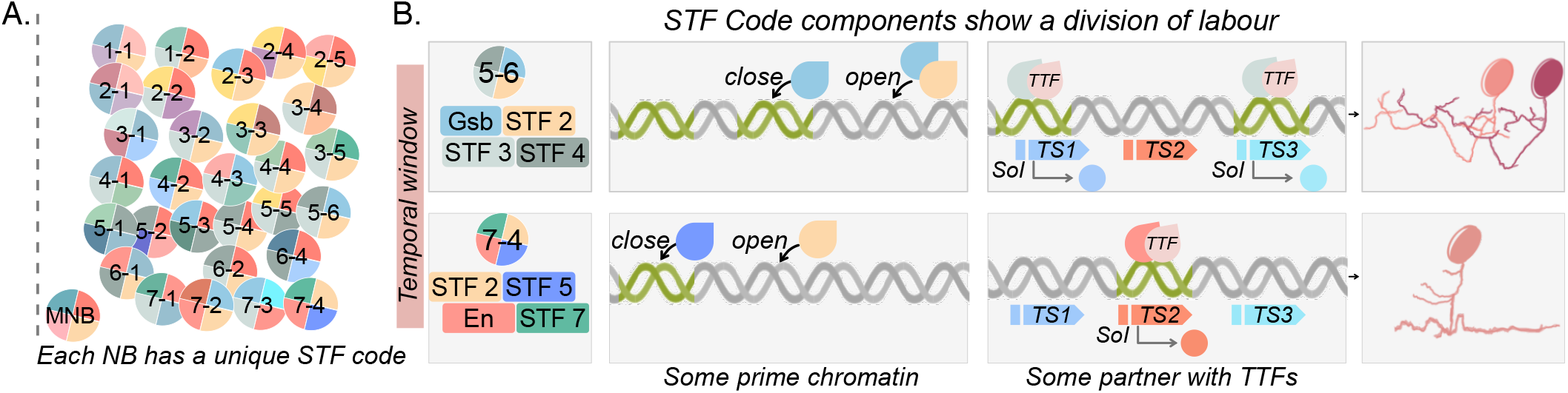
The distributed STF code model for spatiotemporal integration to specify lineage-specific neuron identities. **A. Each NB expresses a unique STF code.** Schematic of NBs in one hemisegment of the *Drosophila* embryonic VNC, adapted from (63). Each NB is shown as a pie whose coloured sectors represent the STFs comprising its code. Dashed line indicates midline. **B. Proposed mechanism**. NB5-6 and NB7-4 are shown as illustrative examples. Each has a distinct set of STFs including Gsb (blue) in NB5-6 and En (red) in NB7-4. Within a shared temporal window (pink bar), code members show a division of labour. Some prime the chromatin landscape, closing loci that would otherwise be available and opening others, thereby establishing lineage-specific accessibility (green DNA, accessible; grey DNA, less accessible). Other code components partner with TTFs at these primed loci to activate terminal selector (TS) genes. Lineage-specific sites integrating STF and TTF inputs are called Sites of Integration (SoIs). Because the two lineages differ in which loci are accessible, the same TTF engages different SoIs and activates a different TS repertoire (TS1 and TS3 in NB5-6; TS2 in NB7-4; indicated as coloured circles), which specifies the identity of the neurons each NB produces (right), linking upstream STF codes to final neuronal fates. STFs and TTFs are drawn as coloured drops.

Several observations in the NB5-6 lineage point to distributed chromatin-priming activity across an STF code, rather than assigning this activity to Gsb alone. Gsb binds both cognate NB5-6 SoIs and non-cognate NB7-4 SoIs, yet only the cognate NB5-6 SoIs become accessible and co-bound by Hb. The non-cognate NB7-4 SoIs remain relatively inaccessible and Hb-free despite Gsb occupancy. Similarly, when Gsb is ectopically introduced into the NB7-4 lineage, it only modestly increases accessibility at non-cognate NB5-6 SoIs and does not install a full NB5-6-like SoI landscape. The remodelling data further suggest that Gsb’s opening and closing effects might be mechanistically distinct: losses of accessibility are strongly associated with Gsb occupancy in independent datasets, whereas gains of accessibility are not. Thus, Gsb can engage loci and directly restrict chromatin accessibility at bound sites, but its ability to promote accessibility and Hb recruitment likely requires additional NB5-6-specific inputs (Figure 9B). This points to a distributed STF-code model in which different code members contribute distinct steps in SoI selection and activation.

The NB7-4-lineage data point to a similar division of labour across a distributed STF code. En occupies accessible NB7-4 SoIs with Hb, but does not bind the non-cognate NB5-6 SoIs in the NB7-4 lineage, even though En motifs are present at these loci. Thus, En can participate in direct STF–Hb engagement at NB7-4 SoIs, but on its own cannot explain how those SoIs are made accessible. As in NB5-6, the full regulatory logic of chromatin priming and TTF integration therefore cannot be assigned to a single candidate STF alone. Instead, it points to additional NB7-4-specific inputs that establish or maintain the chromatin landscape that En and Hb subsequently engage (Figure 9B).

### Gsb and En in the context of known chromatin-associated TF activities

This division of labour is consistent with what is known about these TF classes. Gsb is a Pax3/7-class TF required for row-5/6 NB identity, and vertebrate Pax3/7 proteins can engage relatively inaccessible or nucleosomal chromatin, but their ability to generate productive enhancer opening is strongly cofactor-dependent (10, 51–55). This parallels our finding that Gsb can engage less-accessible loci in the NB5-6 lineage, but productive accessibility and Hb recruitment require additional NB5-6-specific inputs.

En provides a contrasting example. En can both activate and repress its target genes, often via chromatin-modifying cofactors. As a repressor, En’s eh1 motif recruits Groucho/TLE co-repressors, which in turn recruit HDAC/Rpd3, providing a route to local deacetylation and compaction (56–59). As an activator, En can partner with CBP/p300 (Nejire) at a regulatory element *in vivo*, linking En-dependent activation to histone acetylation (60). En can also act as a *bona fide* activator at specific targets, such as *polyhomeotic* (60–62). This fits a role for En in modulating already competent enhancers in the NB7-4 lineage, rather than establishing the accessibility landscape itself.

### The STF code for each NB

The distributed STF-code model we propose here also provides a mechanistic explanation for classic maps of NB identity (Figure 9A). Foundational studies in the VNC and brain showed that individual NBs are identified by unique combinations of transcription factors and molecular markers, set by orthogonal A–P and D–V patterning systems (63, 64). By cataloguing these combinations, these studies effectively defined NB-specific TF codes, although how such codes act at the level of enhancer control was not yet clear. Our data provide one possible mechanistic interpretation of these codes.

A NB’s STF code can be thought of as the minimal combination of STFs sufficient to confer lineage identity by shaping the chromatin landscape and specifying how the temporal cascade engages it — whether through STF-driven chromatin priming, direct STF–TTF co-occupancy on pre-accessible DNA, restriction of inappropriate enhancers, or, in some contexts, TTF-led chromatin remodelling with STFs selecting targets. In this view, direct and epigenetic regulation are not competing lineage-level models, but separable molecular activities that can be distributed across members of the same STF code (Figure 9B).

The corollary is testable: closely related NBs should be most easily interconverted by swapping the relevant code components. An extreme example of this was shown in the central brain, where manipulation of a single TF, *otd*, in a pair of closely related NB lineages could completely change their identity, rewiring both morphology and function (65). One possibility is that *otd* represents the principal differing component between two otherwise similar STF codes. Outside such rare cases, single-factor perturbations more often bias or disrupt lineage programs than produce clean identity swaps: in olfactory projection neurons, individual TF perturbations commonly affect targeting, survival, marker expression, or wiring specificity (66–69), whereas predictable, morphologically and transcriptionally coherent conversions in the optic lobe have emerged from single-cell-atlas-guided manipulation of terminal-selector codes (70). Our Gsb manipulations fit this broader pattern: Gsb loss and gain shift NB identity in reciprocal directions, but neither perturbation fully interconverts NB5-6 and neighbouring row identities. More complete identity changes are therefore likely to require co-manipulation of multiple code members.

Several candidate code components now become important to test. Msh is a shared D–V input in both NB5-6 and NB7-4 and may provide a common component whose activity is interpreted differently depending on accompanying A–P factors. For NB5-6, motif enrichment at lineage-specific SoIs points to additional neuroectodermal patterning factors, including Iroquois-complex members, as candidate code components. For NB7-4, enriched motifs at NB7-4 SoIs may mark factors that establish or maintain the accessibility landscape that En later engages. These candidates remain hypotheses, but they provide a route toward testing code mechanism.

NB6-4 will be a particularly informative future test because it co-expresses both Gsb and En. This lineage provides a natural setting in which a putative chromatin-priming factor and a Hb partner are present in the same NB. Profiling Gsb, En, Hb, and chromatin accessibility in NB6-4 would reveal whether Gsb-occupied sites, En-occupied sites, and Hb-bound SoIs overlap at NB6-4 SoIs. Perturbing Gsb and En individually and in combination would then test whether the two factors do in fact act within the NB6-4 STF code to determine lineage identity. Such systematic code mapping across NBs — extending the classic NB maps with perturbation and chromatin readouts — should reveal how changes in code composition generate the observed diversity of neuronal lineages.

### Scope and future directions

In this work we have used TaDa to profile STF occupancy and chromatin state in identified *Drosophila* NB lineages *in vivo* and propose that NBs integrate spatiotemporal information by using NB-specific STF codes that shape the chromatin landscape to guide TTF binding. Yet the methods used and the biology of the system have important limitations that are worth mentioning. First, the lineage-specific Gal4 drivers also label the immediate progeny of each NB; thus, although profiling was restricted to the early Hb competence window, when only a small number of GMCs and early-born neurons are present, the resulting profiles are not exclusively derived from the NB itself. DamOnly provides a useful *in vivo* proxy for chromatin accessibility in rare, genetically defined NB lineages, and GATC sites are frequent in the *Drosophila* genome, with reported median spacings of approximately 190–200 bp (44, 71); nevertheless, the effective resolution of TaDa-based accessibility is limited by the frequency and local distribution of these sites, and does not provide nucleotide-or nucleosome-scale information. Furthermore, it does not report instantaneous binding or nucleosome position, and instead reports cumulative Dam accessibility over the interval between Gal4 activation, DamOnly expression, and DNA extraction and digestion. Likewise, Gsb misexpression tests chromatin-remodelling capacity outside the native NB5-6 context, and the current tools in this experimental system did not allow us to test the chromatin effects of Gsb loss of function directly in NB5-6. Downstream of these mechanisms, the SoIs remain to be functionally validated, for example by targeted enhancer perturbations while assaying terminal-selector expression and lineage identity. Finally, it will be important to identify the other members of the minimal STF code for NB5-6 and NB7-4 in particular, but more generally for all NBs. New approaches such as spatial transcriptomics should make it possible to identify candidate code members in a less biased way, especially when combined with perturbation and chromatin readouts.

### Relation to vertebrate neural tube

Shared principles are seen in vertebrates. In the spinal cord, opposing morphogen gradients, including ventral Shh and dorsal BMP/WNT signals, establish discrete progenitor domains along the dorsoventral axis, while Hox genes diversify neuronal identities along the anteroposterior axis (72). Temporal patterning, first defined in the developing cortex, has more recently been described across vertebrate CNS regions, including the spinal cord, where temporal transcriptional programs coordinate the sequential production of neuronal subtypes (73).

These vertebrate studies raise the same central question addressed here: how are spatial and temporal cues integrated at cis-regulatory elements? A recent study proposed a “chronotopic” strategy in vertebrate neural progenitors, in which temporal regulators drive a global temporal chromatin program over developmental time, and spatial determinants act on the regulatory elements made available in each progenitor domain (74). This is an interesting comparison to the model proposed here, where spatial STF codes help shape the chromatin landscape that Hb engages. However, the two studies differ in experimental scale and context. Our assays resolve transcription factor occupancy and chromatin accessibility in single identified stem cell lineages *in vivo*, whereas the vertebrate work examines regulatory logic at the level of progenitor domains, across time-windows, *in vitro*. Apparent differences in the relative contributions of spatial and temporal factors may therefore reflect biological differences, differences in developmental scale, or differences in assay design.

For now, the most useful conclusion is that chromatin provides a common substrate for integrating spatial and temporal information. Whether spatial factors first establish competence, temporal factors broadly remodel competence, or both occur in different contexts will require more direct comparisons across systems and scales. Even within *Drosophila*, other NBs or temporal windows may use different implementations of this logic, including TTF-led chromatin remodelling, as suggested by the pioneer activity of Grainy head (75). In the embryonic NBs studied here, our data suggest that integration is implemented through a NB-specific STF code that shapes the enhancer landscape available to Hb. More broadly, combinatorial transcription factor codes may provide a general strategy for generating lineage-specific neuronal outputs from shared temporal programs.

## Materials and Methods

All fly stocks are listed in Table 1. All antibodies used are listed in Table 2.

**Table 1.**
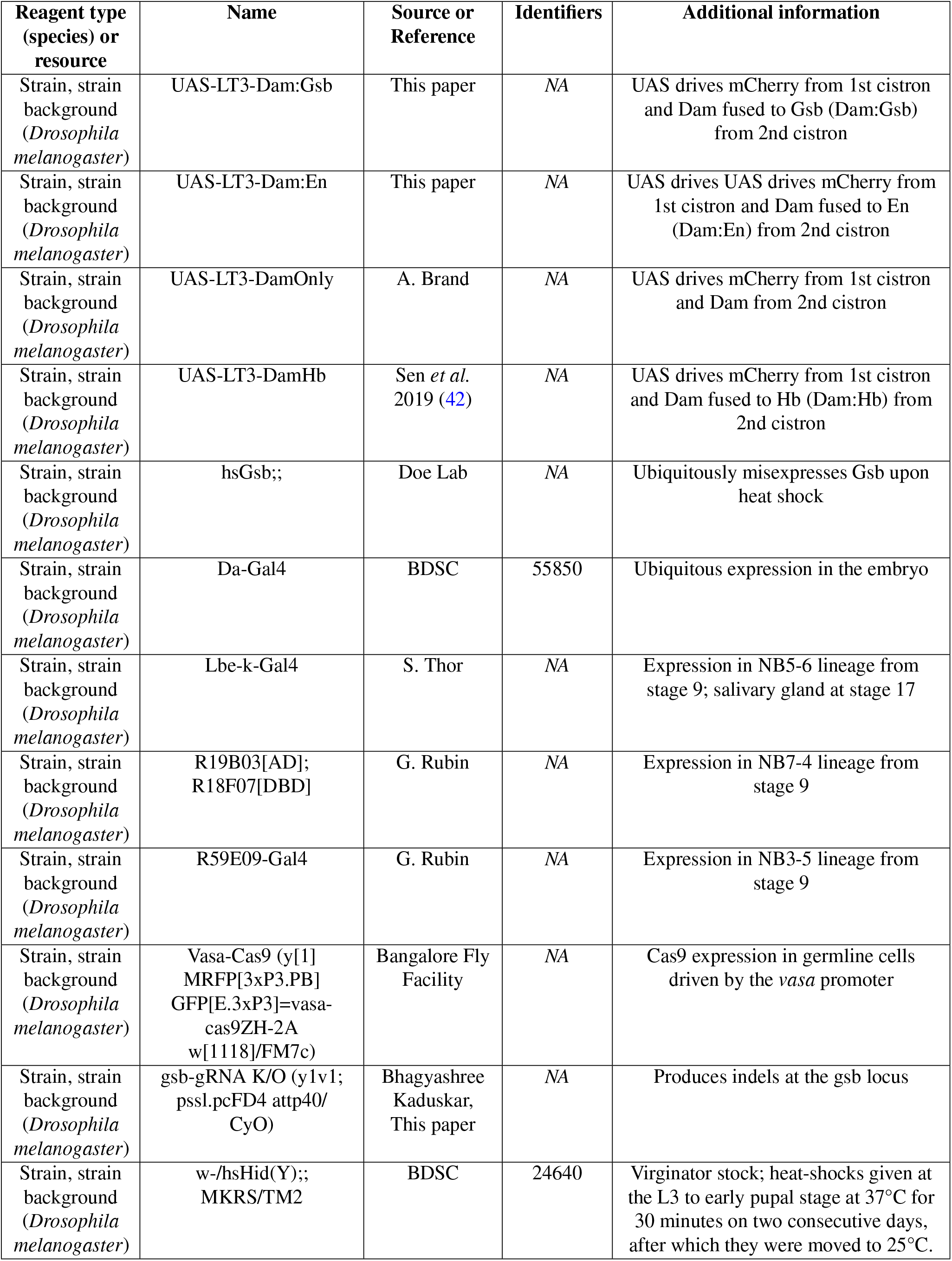
Resource Table: List of fly lines used.

**Table 2.**
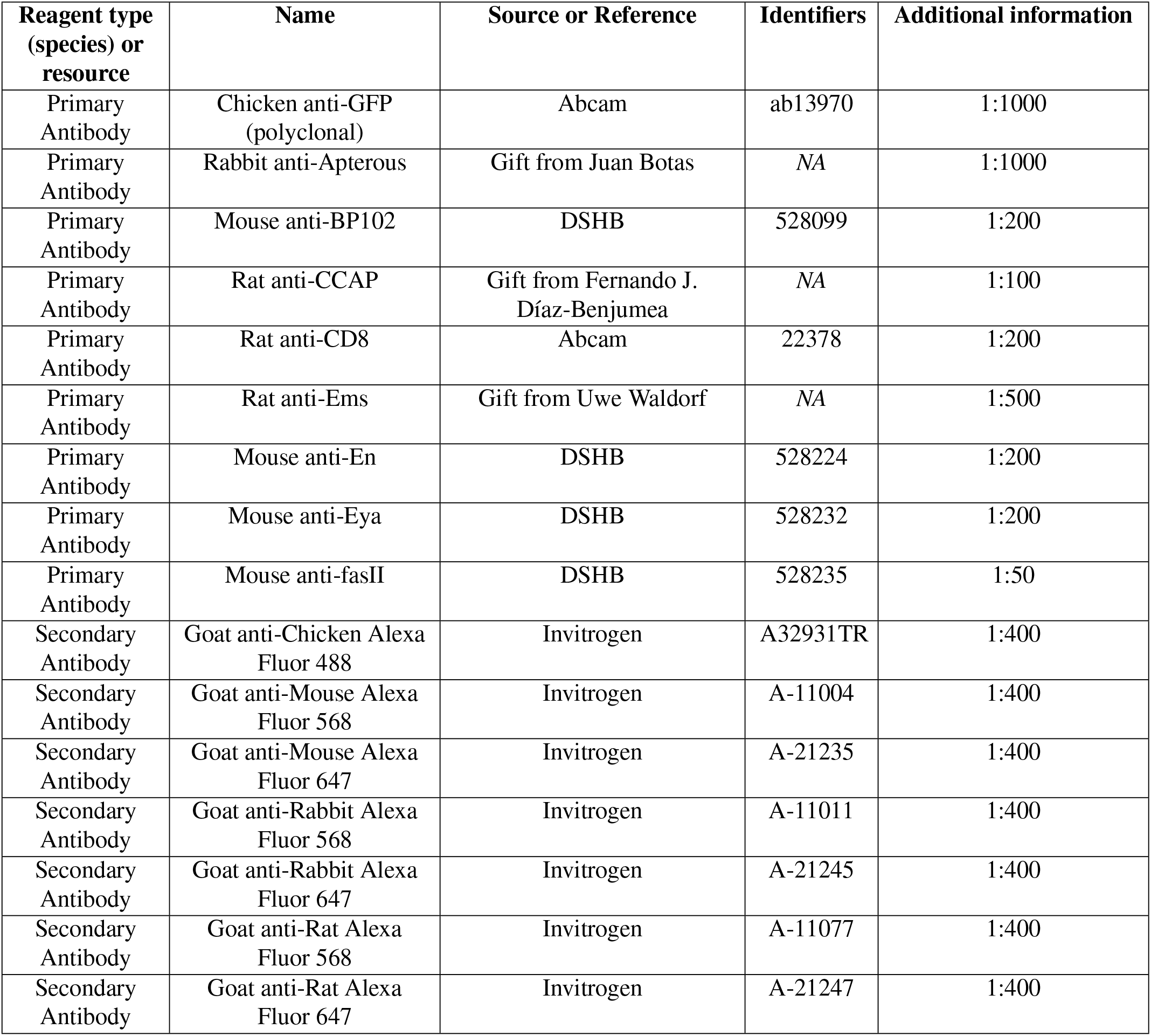
Resource Table-Antibodies Used.

### Fly Stock Maintenance

Fly stocks were maintained on a standard cornmeal medium. The recipe consisted of the following per litre: 80g corn flour, 20g glucose, 40g sugar, 15g yeast extract, 4mL propionic acid, 5mL p-hydroxybenzoic acid methyl ester (in ethanol), and 5mL ortho-butyric acid. Flies were raised at 25°C under a 12h:12h light-dark cycle. The virginator *hsHid(Y)* and *hsGsb* stocks were grown at room temperature.

### Generation of gsb-guideRNA fly lines

To generate the gsb-guideRNA line, *Drosophila gsb*-guideRNA sequences were cloned into the tandem guideRNA expression vector pCFD4 (76). The gRNA target sequences were amplified via PCR using *gsb*-specific primers, and the resulting PCR product was assembled into the BbsI-digested pCFD4 backbone using Gibson assembly. Transgenic lines were established by standard balancing. To validate editing efficiency, *gsb*-guideRNA lines were crossed to *Vasa-Cas9* flies. Genomic DNA was extracted from embryos and the *gsb* target locus was amplified by PCR and submitted for Sanger sequencing. Sequencing chromatograms were analysed using the Synthego ICE (Inference of CRISPR Edits) online tool to estimate editing efficiency (Figure 7—figure supplement 1C–D).

### TaDa and CATaDa Experiments

#### Verification of Dam:Gsb and Dam:En

To assess whole-embryo occupancy of Gsb and En, Dam fusion constructs were expressed using the ubiquitous Da-Gal4 driver. 1000– 1600 females of *w-;;Da-Gal4* were crossed to a similar number of males of *w-;;UAS-LT3-Dam:Gsb* for Gsb occupancy, *w-;;UAS-LT3-Dam:En* for En occupancy, and *w-;;UAS-LT3-DamOnly* for chromatin accessibility.

After 16–24h of mating, adults were moved to embryo collection cages. Collection plates were prepared using 2% agar and 2% sucrose in 60–100mm Petri dishes, and yeast paste was applied to stimulate egg laying. Embryos were collected every 4h (0–4h AEL) and then aged for 12h (12–16h AEL; Stages 15–16) at 25°C to match the temporal window of publicly available ChIP-seq datasets used for validation.

Following ageing, embryos were rinsed with distilled water and dechorionated in bleach for 2min (or until chorion removal was visually confirmed). Embryos were then washed thoroughly with distilled water, dried on filter paper, weighed, and stored in 1.5mL tubes at –20°C. A minimum of 30mg per replicate was collected. This procedure was performed as two independent biological runs, yielding three replicates each of Dam:Gsb and DamOnly, and three each for Dam:En and DamOnly, profiled across the whole embryo.

#### Determination of NB-specific STF occupancy and chromatin accessibility

To determine Gsb occupancy and chromatin accessibility in NB5-6, ~ 4000 males of *w-;Lbe-k-Gal4* were crossed to a similar number of females of *w-;;UAS-LT3-Dam:Gsb* or *w-;;UAS-LT3-DamOnly*. Similarly, to determine En binding and chromatin accessibility in NB7-4, ~ 4000 males of *w-;19B03[AD];18F07[DBD]* were crossed to a similar number of females of *w-;;UAS-LT3-Dam:En* or *w-;;UAS-LT3-DamOnly*. Embryos were collected for 2h (0– 2h AEL) and then aged for 6h (6h–8h AEL, Stage 11–12) at 25°C. We collected replicates with at least 30mg of embryos for each condition. This procedure was performed as two independent biological runs, yielding four Dam:Gsb and four DamOnly replicates for NB5-6 and three Dam:En and three DamOnly replicates for NB7-4.

#### Determination of NB-specific Hb occupancy

NB-specific Hb occupancy data (Stage 11–12) were obtained from (42).

#### Determination of NB7-4-specific chromatin accessibility and Hb occupancy, upon broad Gsb misexpression

To assay chromatin accessibility and Hb occupancy in NB7-4 under conditions of broad Gsb misexpression, ~ 5000 females each of *hsGsb;;UAS-LT3-DamOnly* and *hsGsb;;UAS-LT3-Dam:Hb* were crossed to a similar number of males of *w-;19B03[AD];18F07[DBD]*. Embryos were collected every 1h (0–1h AEL), incubated at 25°C for 3h (3–4h AEL, Stage 7–8), and then moved to 33°C just before NB delamination for 4h (7–8h AEL, Stage 11–12), inducing Gsb misexpression. Three replicates of ≥ 30mg were collected per condition.

#### TaDa/CATaDa Experimental Pipeline

TaDa and CATaDa replicates were processed as per (77). Briefly, embryos were homogenised with motorized pestles, and genomic DNA was extracted using the QIAamp DNA Micro Kit (Qiagen, 56304; 50µL final volume) using wide-bore tips throughout to minimise shearing. DNA integrity was confirmed on a 0.8% agarose gel. Genomic DNA was digested with DpnI (NEB, R0176S) to cut Dam/Dam:TF-methylated sites, followed by DamID adaptor ligation and DpnII digestion (NEB, R0543S) to fragment unmethylated DNA. Adaptor-ligated fragments were PCR-amplified (21 cycles; MyTaq HS DNA Polymerase, Bioline BIO-21112). Libraries were prepared at the Next Generation Genomics Facility (NGGF), BLiSC, using the NEBNext® Ultra™ II DNA Library Prep with Sample Purification Beads (E7103L), and sequenced on Illumina HiSeq2500 (50bp single-end, 20–60 million reads/sample). Gsb-misexpression TaDa/CATaDa samples were sequenced on NovaSeq 6000 (50bp paired-end, 50–60 million reads/sample). Quality metrics associated with all sequencing experiments are summarised in Supplementary sheet 1.

### Data processing and analysis

#### Alignment and filtering

Reads were aligned to the *Drosophila* genome (dm6) using `Bowtie2` (78). The resulting .*sam* files were converted to .*bam* format and sorted using `SAMtools` (79). Reads were then filtered using `NGSUtils` (80) to exclude those mapping to ENCODE blacklisted regions in dm6 or to the mitochondrial genome. Reads with mapping quality below MAPQ 5 were further removed using `SAMtools`. Filtered .*bam* files were indexed using `SAMtools` and used for all downstream analyses.

Overall replicate consistency was assessed by pairwise Pearson correlation of .*bam* files using `multiBamSummary` and `plotCorrelation` from the `deepTools` suite (81) (Figure S1B, F, Figure 2C–D).

#### Processing of published ChIP-seq data

Raw ChIP and input FASTQ files for Gsb (GSE256559 (82)), Twist (GSE256656 (83)), and Bicoid (GSE257099 (84)), generated by the modENCODE consortium (45), were downloaded and aligned to the *Drosophila* genome (dm6) using `Bowtie2`. Resulting .*sam* files were converted to .*bam* format, sorted, and indexed using `SAMtools`. Peaks were called for each ChIP dataset using `MACS2` (85), with the corresponding input as control (`-m 3 30 -q 0.0001`). Whole-embryo Dam:Gsb signal was plotted at ChIP peak summits using `computeMatrix reference-point`, allowing comparison of TaDa-derived Gsb occupancy at published Gsb-ChIP, Twi-ChIP, and Bcd-ChIP peaks (Figure S1D).

#### Motif discovery in whole embryo STF-TaDa data

De novo and known motif enrichment analyses were performed on Dam:TF peak sets using `findMotifsGenome.pl` from `HOMER` (86), using the dm6 genome with default parameters. Top enriched motifs were identified by ranked *p*-value, with percentage of target and background peaks containing each motif reported (Figure S1E,H).

#### Peak calling for Dam:TF samples

Narrow peaks were called on Dam:TF .*bam* files using `MACS2` (v2.2.6) (85), with concatenated DamOnly replicates from the corresponding condition as control. The `-nomodel` and `-nolambda` parameters were used to disable fragment size modelling and dynamic local background estimation, respectively. Read extension was set to 300bp (`-extsize 300`) to account for fragment size. Duplicate reads were handled using `-keep-dup auto`, which retains duplicates up to the threshold expected by chance given library size, based on a binomial distribution. An FDR of ≤ 0.01 was used as the significance threshold.

This approach was applied consistently across all Dam:TF datasets, including NB-specific Hb occupancy data from (42), for which raw data were re-processed using the same pipeline to ensure comparability.

#### Differential analysis on Dam:TF data

Differential analyses were performed on Dam:TF peaks using DiffBind (87, 88).

#### (i) Determining Sites of Integration (SoIs)

SoIs – loci that are Hb-bound and accessible in one NB but not the other – mark candidate sites of STF–TTF convergence. These were defined using differential analysis in NB-specific Hb-TaDa data from (42). Peaks were considered differential at FDR ≤ 0.01 and fold change ≥ 2 between NB5-6 and NB7-4, yielding 865 NB5-6-SoIs and 245 NB7-4-SoIs. Shared Hb-bound sites were identified using `bedtools intersect` with a minimum overlap of ≥ 50%, yielding 1,140 shared sites. SoI definitions were validated by plotting Dam:Hb and DamOnly signal at these site sets, confirming differential Hb occupancy and accessibility (Figure 3—figure supplement 1A–F). Genomic coordinates and associated signal values are summarised in Supplementary sheet 3.

#### (ii) Determining genome-wide STF-bound and -unboundsites

NB5-6 Dam:Gsb and NB7-4 Dam:En peaks were called at a stricter threshold of FDR ≤ 1 *×* 10−5. STF-bound and -unbound sites were determined by differential analysis between these peak sets, considered differential at FDR ≤ 0.01 and fold change ≥ 2. This yielded 1,122 Gsb-bound/En-unbound sites and 807 En-bound/Gsb-unbound sites. Sites bound by both STFs were identified using `bedtools intersect` with a minimum overlap of ≥ 50%, yielding 740 shared sites. Site definitions were validated by plotting Dam:Gsb and Dam:En signal at these site sets, confirming differential STF occupancy (Figure 3—figure supplement 3A–D). Genomic coordinates and associated signal values are summarised in Supplementary sheet 4.

#### Chromatin accessibility as a predictor of STF binding

To test whether chromatin accessibility predicts STF binding, DamOnly signal (NB5-6: four replicates; NB7-4: three replicates) was quantified at the genome-wide STF-bound and -unbound site sets defined above (1,122 Gsb-bound/En-unbound and 807 En-bound/Gsb-unbound sites; the 740 shared sites were excluded) using `computeMatrix` (`deepTools`).

For each site, signal was averaged across bins within a central window (bins 250–350, *±* 500bp from site centre) and across replicates to yield one accessibility value per site. Accessibility was then compared between the corresponding bound and unbound sets using a two-tailed Mann–Whitney U test, with effect size and direction quantified by Cliff’s *δ* (Figure 3G–I, Figure 4G–I). Classifier performance was assessed by ROC analysis (`pROC`, R), with bound sites as the positive class and per-site accessibility as the predictor; AUC was reported as the probability that a randomly selected bound site has higher accessibility than a randomly selected unbound site (Figure 3—figure supplement 3E).

#### Determining STF-only, TTF-only, and STF+TTF co-bound sites

The top 5,000 `MACS2` FDR-ranked peaks were used for each TF: Gsb in NB5-6, En in NB7-4, and Hb in both NB5-6 and NB7-4. For each TF, peaks were visually inspected on IGV genome browser and confirmed to include known target loci. STF-only and TTF-only sites were identified using `bedtools intersect` with the `-v` parameter to exclude overlapping peaks. Co-bound (STF+TTF) sites were defined using `bedtools intersect` with ≥ 90% overlap, yielding 1,568 Gsb+Hb sites in NB5-6, 1,419 En+Hb sites in NB7-4, 2,322 Gsb-only sites, 2,151 En-only sites, 2,571 NB5-6 Hb-only sites, and 2,460 NB7-4 Hb-only sites. These site sets were used to assess chromatin accessibility at co-bound versus singly-bound loci (Figure 5). Genomic coordinates and associated signal values are summarised in Supplementary sheet 5.

#### Annotating genes near SoIs

SoI peaks (NB5-6 and NB7-4) were annotated to nearby TF-encoding genes using `ChIPpeakAnno` (89) with the *Drosophila* dm6 gene annotation (`TxDb.Dmelanogaster.UCSC.dm6.ensGene`). `annotatePeakInBatch` was run with `output =` “`both`” to report all genes overlapping or flanking each peak within a specified distance, rather than only the single nearest gene. A 10kb maximum distance threshold was selected following a sensitivity analysis testing 10, 25, and 50kb windows. This range was informed by REDfly-curated (90) enhancer-to-TSS distances for known lineage markers (*lbe, ap, en, acj6*: 0.5–50kb). Recovery of known markers did not improve at larger thresholds, so 10kb was used as the most conservative window supporting marker recovery. Gene symbols were assigned using `org.Dm.eg.db`, mapping FlyBase IDs to gene symbols (91). Genes were classified as TF-encoding based on membership in the FlyBase Transcription Factors gene group (FBgg0000745). For each TF-encoding gene near an SoI, fold change and FDR values from the corresponding differential Hb occupancy analysis (see above) were assigned by genomic coordinate overlap. Candidates were ranked using a combined score incorporating fold change, statistical confidence, and distance to the nearest TSS:

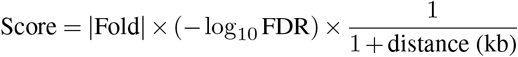

This score prioritises genes with strong differential Hb occupancy, high statistical confidence, and proximity to the SoI, while not excluding more distal candidates. Duplicate gene entries were collapsed to retain the highest-scoring occurrence per gene. The top 20 candidates per lineage were visualised; known lineage markers (*lbe* for NB5-6, *en* for NB7-4) were included regardless of rank if not already present in the top 20 (Figure 3—figure supplement 1J–K).

#### En binding motif occurrence at SoIs

To test whether the absence of En binding at NB5-6-SoIs in NB7-4 reflects a lack of appropriate binding sequence context, we assessed En motif occurrence at NB5-6-SoIs and NB7-4-SoIs using `FIMO` from the MEME suite (92), with the En position weight matrix obtained from JASPAR 2026 (93). The percentage of peaks containing at least one motif occurrence was calculated for each site set.

To establish a background expectation, 500 size- and composition-matched shuffled peak sets were generated using `bedtools shuffle` and scanned identically. The empirical *p*-value for each real site set was calculated as the fraction of shuffled sets with motif occurrence equal to or greater than the observed value, with a minimum reportable *p*-value of 1*/*500 = 0.002 given the number of permutations. NB7-4-SoIs, which are En-bound, served as a positive control (Figure 3—figure supplement 1L).

#### Defining the ranked chromatin accessibility spectrum

Since accessibility is continuous rather than binary open/closed, we defined an accessibility landscape with regions relevant to both NBs: Dam-bound broad peaks reproducible in at least two replicates in either NB were considered. These regions were binned into ranked deciles by mean accessibility using DamOnly signal from both NBs, yielding a representation of different accessibility states (bin 1, least accessible; bin 10, most accessible) (Figure 2—figure supplement 2A). Genomic coordinates and associated signal values are summarised in Supplementary sheet 2.

#### (i)DamOnly peak calling

1. Since DamOnly binding is diffuse, broad peaks were called on individual DamOnly .*bam* files using `MACS2` with the following parameters: `-broad -broad-cutoff 0.1 -nomodel -extsize 300`, without controls, FDR ≤ 0.01.

#### (ii)Defining the accessible landscape

- Reproducible peaks (present in ≥ 2 replicates) were identified per NB, then merged across NBs (`bedtools merge`, 500bp max gap) into a union peakset. Since broadpeak lengths varied widely (mean 4,296bp, median 3,157bp, range 300–52,891bp), peaks were standardised to a 3kb maximum, reflecting median length. Peaks <3kb were retained and longer peaks split into 3kb windows (`bedtools makewindows`), yielding 27,119 regions defining the accessible landscape used throughout.

#### (iii) Binning the accessible universe into a ranked spectrum

DamOnly signal across all 27,119 regions was quantified for each replicate using `multiBigwigSummary` from `deepTools` (81), using RPGC-normalised .*bw* files. Signal values were log_2_(*x* + 1) transformed to stabilise variance. For each NB, mean accessibility was calculated as the average of log-transformed replicate values. This combined mean accessibility score was computed across both NBs and used to rank all 27,119 regions. Regions were stratified into ten ranked deciles (~ 2,700 regions per bin), from least (bin 1) to most (bin 10) accessible.

#### NB-bias in accessibility across the defined accessibility landscape

NB-specific bias was calculated as the difference in mean log-transformed DamOnly signal (NB5-6 NB7-4), i.e. a log_2_ fold-change (positive = NB5-6-preferential, negative = − NB7-4-preferential). Replicate consistency was confirmed by pairwise Pearson correlation (`multiBigwigSummary`; visualised in R; Figure 2— figure supplement 2B).

Bias across the ranked spectrum was plotted per bin (jitter/boxplot, points = regions coloured by bias direction, boxes = median/IQR). A one-sample two-sided Wilcoxon signed-rank test assessed deviation from zero per bin; effect size 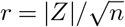 (Figure 2G).

Regions with accessibility bias| ≥1 were extracted from the 27,119-region accessibility landscape, corresponding to a ≥ 2-fold difference in DamOnly signal between the two NBs. This yielded 845 NB5-6-preferential sites (NB56 bias ≥ − 1) and 1,067 NB7-4-preferential sites (NB56 bias ≤ 1). NB-preferential sites were exported as .*bed* files for downstream analyses and genome browser visualisation. NB5-6 preferential sites were considered accessible in NB5-6 but less accessible in NB7-4, and vice versa. This definition was validated by plotting DamOnly signal back at these NB-preferential sites Figure 3—figure supplement 2B–D).

#### Profiling STF occupancy across the defined accessibility landscape

STF occupancy across the 27,119 regions was quantified, transformed, and averaged as for DamOnly above. Sina plots showed per-site mean occupancy per bin (mean *±* SEM overlaid; Figure 3—figure supplement 2A).

#### Determining changes in chromatin accessibility and Hb occupancy in NB7-4 upon Gsb misexpression

DamOnly signal from control and Gsb-misexpressed NB7-4 was quantified, log_2_ transformed, and averaged across replicates per condition as above, within the 27,119-region universe. Accessibility or Hb occupancy change (test control, log_2_ fold-change; positive = gain, negative = loss) was confirmed reproducible by Pearson correlation (Figure 6—figure supplement 2A, C). Change across the ranked spectrum was assessed per bin by one-sample Wilcoxon signed-rank test 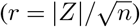 and summarised as mean ≥ SEM per bin (Figure 6—figure supplement 2E, F). Regions with accessibility change 1 yielded 606 gain and 1,340 loss sites, exported as .*bed* files. To confirm that these accessibility changes were accompanied by corresponding changes in Hb occupancy, DamOnly and Dam:Hb signal from control and Gsb-misexpression conditions were mapped at gain and loss sites using `computeMatrix` (*±* 1.5kb around site centres, 10bp bins) (Figure 6 C–D, F–G). Hb occupancy change was calculated as log_2_ fold-change (test − control) across this window, following the same convention as accessibility change above. To test whether native Gsb occupancy is associated with sites undergoing remodelling in accessibility, Dam:Gsb signal from NB5-6 (stage 11–12) and whole embryo (stage 16– 17) was mapped at gain and loss sites (Figure 6 E,H). Gsb occupancy was associated with sites undergoing accessibility loss, but not sites undergoing gain.

#### Signal metaplots and heatmaps

Signal files were generated from filtered .*bam* files using `bamCoverage` (RPGC-normalised) and visualised in IGV (94, 95). `computeMatrix` (`deepTools`) generated per-bin signal matrices (±3kb around peak centres, 10bp bins); mean signal across replicates was plotted in R with *±* SD ribbons. Heatmaps show mean replicate signal ordered by decreasing signal in the relevant control/reference condition unless otherwise stated.

#### Gsb knockdown and misexpression experiments

##### (i) Gsb knockdown

For Gsb loss-of-function experiments, *Vasa-Cas9* flies were crossed to *y[1]v[1]; gsb-guideRNA* flies. Embryos were collected at the appropriate stage and stained with rabbit anti-Ap, mouse anti-Eya, and rat anti-Ems antibodies (see Table 2). Marker-positive cells per hemisegment were counted from confocal images.

##### (ii) Gsb misexpression

For Gsb misexpression experiments, *hsGsb* flies were crossed to *w-; Lbe-k-Gal4, UAS-mCD8-GFP* (to visualise the NB5-6 lineage) or *w-;; R59E09, UAS-myrGFP* (to visualise the NB3-5 lineage). Embryos were collected (0–1h AEL) and incubated at 18°C for 6.5h, heat shocked at 37°C for 30min, and subsequently incubated at 23°C for 21h to reach Stage 16. Embryos were stained with chicken anti-GFP, mouse anti-Eya, rabbit anti-Ap, and rat anti-Ems antibodies (see Table 2). Marker-positive cells per hemisegment were counted from confocal images.

##### (iii) Quantification and statistics

In all cases, marker-positive cells per hemisegment were counted and plotted as box and whisker plots in R. Each point represents one hemisegment. Statistical significance was assessed using two-tailed Mann– Whitney U tests (∗∗∗ *p <* 0.001, ∗∗*p <* 0.01, ∗*p <* 0.05, *ns* = not significant). Raw quantification of marker-positive cells per hemisegment across experimental conditions are summarised in Supplementary sheet 7.

##### Immunohistochemistry and imaging

Appropriately staged embryos were washed with distilled water and dechorionated in bleach for 2–3 minutes. After thorough rinsing with distilled water, embryos were fixed by nutating in a 1:1 mixture of 4% PFA and heptane for 20–25min. Vitelline membranes were removed by vigorous shaking for 1min in a 1:1 mixture of heptane and methanol. Embryos were then washed in 0.5% PTX (0.5% Triton X-100 in 1 *×* PBS), incubated overnight at 4°C with primary antibodies, washed again in 0.5% PTX, and incubated overnight at 4°C with secondary antibodies. Following additional washes in 0.5% PTX, embryos were cleared in 90% glycerol with Vectashield.

Imaging was performed at the Central Imaging and Flow Cytometry Facility (CIFF), Bangalore Life Science Cluster (BLiSC), using an Olympus FV3000 system with a 60 *×* oil-immersion objective. Images were acquired at 512 *×* 512 or 1024 *×* 1024 resolution, viewed and manually analysed in FIJI (ImageJ) (96), and are presented as maximum-intensity projections of z-stacks.

All statistical analyses and data visualisations were performed in R. All figures were assembled using Affinity Designer v2.

## Supporting information

Supplementary sheet 1

Supplementary sheet 2

Supplementary sheet 3

Supplementary sheet 4

Supplementary sheet 5

Supplementary sheet 6

Supplementary sheet 7

## Data availability

Raw fastq files have been deposited at the National Center for Biotechnology Information (NCBI) Sequence Read Archive (SRA), under the BioProject ID PRJNA1372925. All scripts used in analysis and figure generation are available on Github.

## Author contributions

AB conceptualised the project, performed and analysed all experiments, wrote and reviewed the manuscript. HR conceptualised, performed, and analysed experiments related to the functional analysis of Gsb and reviewed the manuscript. SQS conceptualised and supervised the project and wrote and reviewed the manuscript.

## ACKNOWLEDGEMENTS

This work was supported by the Ramalingaswami Fellowship to SS, UGC-CSIR Fellowship to AB, and TIGS. We acknowledge the Fly Facility at NCBS. All sequencing was done at the NGGF sequencing facility at the Bangalore Life Science Cluster. All imaging was done at the Central Imaging and Flow Cytometry Facility (CIFF) at the Bangalore Life Science Cluster. We thank Keiko Hirono and Bhagyashree Kaduskar for their help in generating fly lines for this project. We are grateful to Vishaka Gopalan, Sridhar Hannenhalli, Sabarinathan Radhakrishnan, Bhavana Muralidharan, Chris Q Doe, Sachin Chanchani and Anton Iyer for invaluable discussions. We are also grateful to Chris Doe and K. VijayRaghavan, for their critical comments on the manuscript. Finally, we thank Sen Lab members for discussions and critical feedback.

## Supplementary Data

### Supplementary files (as attachments)

- **Supplementary sheet 1**: Summary of TaDa and CATaDa sequencing quality metrics.
- **Supplementary sheet 2**: Summary across regions of the accessibility landscape, with or without Gsb misexpression.
- **Supplementary sheet 3**: Summary at sites of integration (SoIs) in NB5-6 and NB7-4.
- **Supplementary sheet 4**: Summary at STF-bound and unbound loci in NB5-6 and NB7-4.
- **Supplementary sheet 5**: Summary at sites that are bound by STF only, Hb only, and co-bound by both STF and Hb in NB5-6 and NB7-4.
- **Supplementary sheet 6**: Genomic coordinates of site sets used in analyses.
- **Supplementary sheet 7**: Raw quantification of marker-positive cells per hemisegment (Ap, Ems, Eya, GFP-reporters) across all experimental conditions. This sheet contains the complete dataset used for all quantifications shown in Figures 7–8.

**Supplementary Figure 1.**
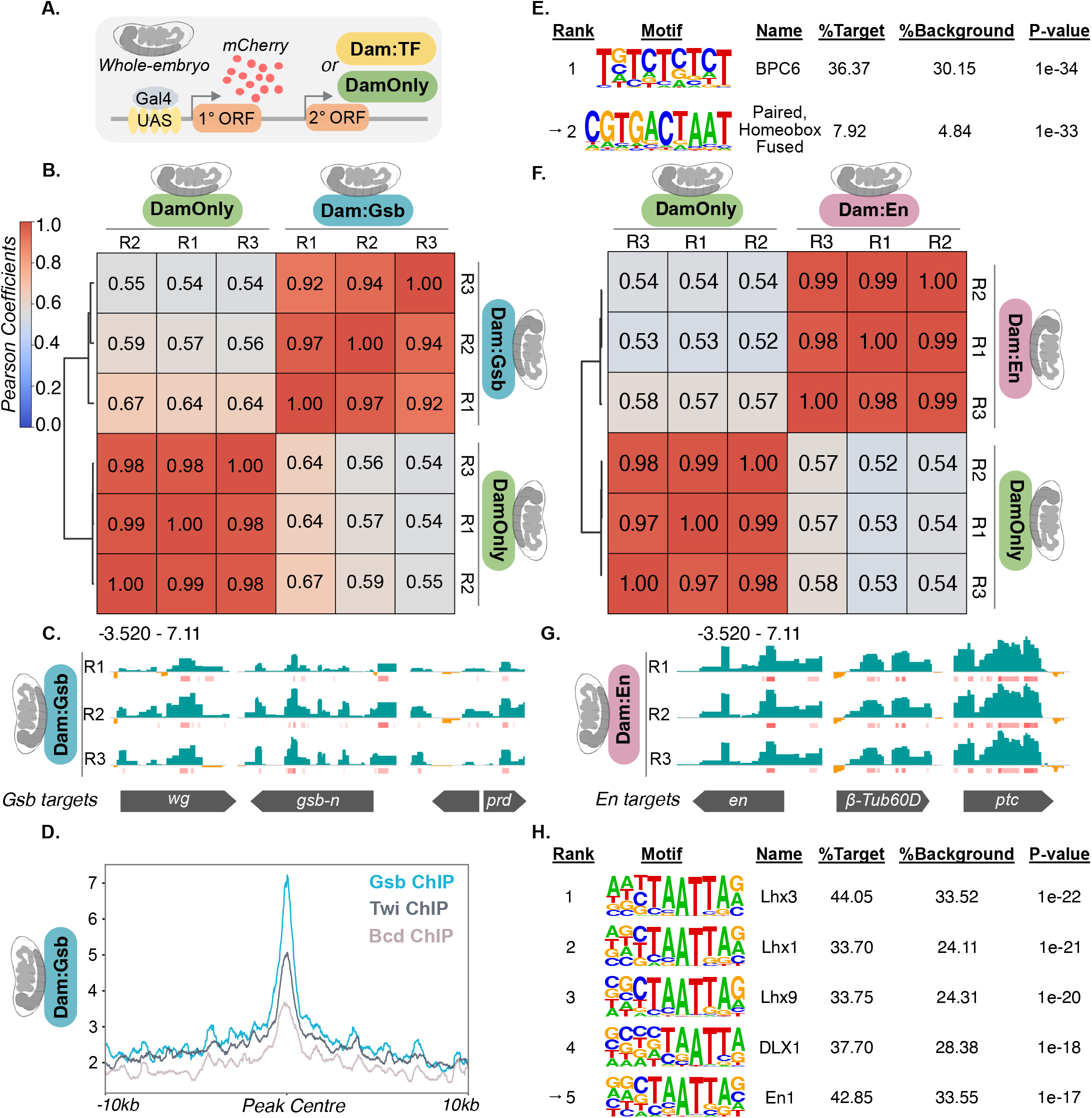
Verification of TaDa Tools. **A. Design of TaDa and CATaDa tools.** The ubiquitous *Da-Gal4* driver expresses mCherry from the primary ORF and either DamOnly (green) or Dam:TF (yellow) from the secondary ORF. High mCherry expression reports driver specificity, while translation of Dam fusions from the secondary ORF keeps Dam levels low, minimizing toxicity and preserving binding specificity. **B. Whole-embryo DamOnly and Dam:Gsb bind reproducibly**. Heatmap shows pairwise Pearson correlations between DamOnly and Dam:Gsb replicates. Hierarchical clustering separates DamOnly and Dam:Gsb samples, with stronger correlations within than between conditions. **C. Binding at known Gsb targets**. Genome browser snapshots of log_2_ ratio files (Dam:Gsb/DamOnly) are shown (3 replicates), Dam:Gsb in blue positive tracks, DamOnly in orange negative tracks. Replicates show occupancy at *wg, gsb-n, prd*. Data range –3.520–7.11. Pink bars indicate peaks, colour intensity indicates peak score. **D. TaDa-Gsb correlates with published Gsb ChIP-seq data**. Dam:Gsb signals are most strongly enriched at Gsb-ChIP peaks relative to Twi-ChIP or Bcd-ChIP sites. **E. Motif enrichment at Dam:Gsb sites**. *De novo* motif analysis at Dam:Gsb peaks reveals a fused Paired–Homeobox motif (arrow, rank 2). **F. Whole-embryo DamOnly and Dam:En bind reproducibly:** Heatmap shows pairwise Pearson correlations between replicates of DamOnly and Dam:En. Hierarchical clustering separates DamOnly and Dam:En samples, with stronger correlations within than between conditions. **G. Binding at known En targets**. Genome browser snapshots of log_2_ ratio files (Dam:En/DamOnly) are shown (3 replicates), Dam:En in blue positive tracks, DamOnly in orange negative tracks. Replicates show occupancy at *en, β-Tub60D* and *ptc*. Data range –3.520–7.11. Pink bars indicate peaks, colour intensity indicates peak score. **H. Motif enrichment at Dam:En sites**. *De novo* motif analysis at Dam:En peaks recovers homeodomain motifs sharing the canonical TAAT core (Lhx/DLX family, Ranks 1–4), with the Engrailed motif recovered at Rank 5 (arrow). Genotypes for embryo-wide Gsb occupancy: *w-;; Da-Gal4/UAS-Dam:Gsb*. For embryo-wide En occupancy: *w-;; Da-Gal4/UAS-Dam:En*. For embryo-wide chromatin accessibility: *w-;; Da-Gal4/UAS-DamOnly*.

**Figure 2—Figure Supplement 1.**
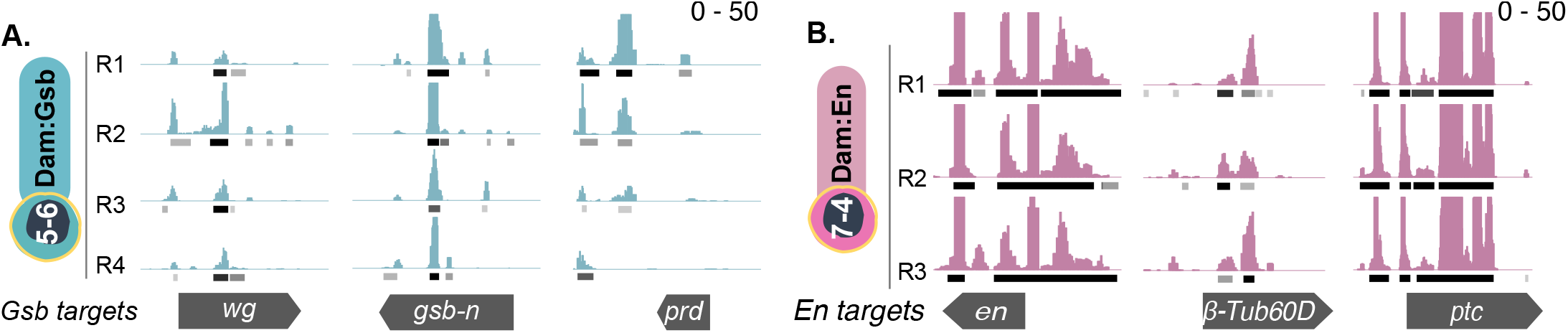
NB-specific Dam:TF at known targets. **A. Binding at known Gsb targets.** Genome browser snapshots of four replicates of NB5-6-Dam:Gsb show binding at known Gsb targets – *wg, gsb-n, prd*. **B. Binding at known En targets**. Genome browser snapshots of three replicates of NB7-4-Dam:En show binding at en, *β*-*Tub60D, ptc*. For all browser snapshots, data range 0-50. Black bars indicate peaks, colour intensity indicates peak score. Genotypes for NB5-6 Gsb occupancy: *w-; Lbe-k-Gal4/+; +/UAS-Dam:Gsb*. NB7-4 En occupancy: *w-; 19B03[AD]/+; 18F07[DBD]/UAS-Dam:En*.

**Figure 2—Figure Supplement 2.**
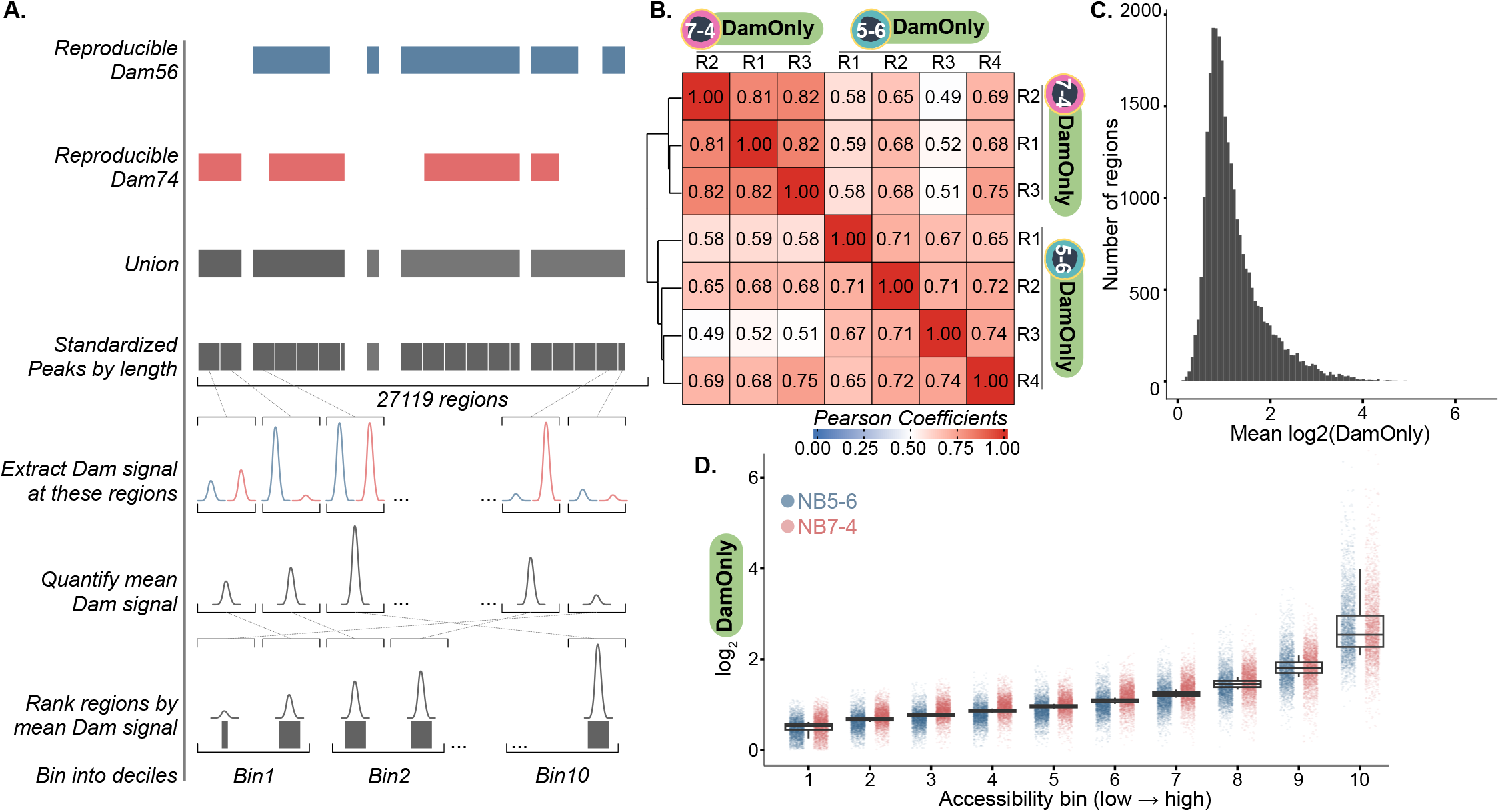
Defining the ranked accessibility spectrum. **A. Construction of the accessibility landscape.** DamOnly peaks from NB5-6 (blue) and NB7-4 (red), reproducibly identified in at least two biological replicates, were merged into a union peakset (dark grey). Peak lengths were standardised to a maximum of 3kb, the median peak length. Longer peaks were split into 3kb windows and shorter peaks retained, yielding 27,119 regions. Regions were ranked by mean accessibility (average DamOnly signal from both NBs) and stratified into deciles (~2,700 regions per bin), from bin 1 (least accessible) to bin 10 (most accessible). **B. Reproducibility of DamOnly replicates across the accessibility landscape.** Pairwise Pearson correlation heatmap of NB5-6 and NB7-4 DamOnly replicates across all 27,119 regions shows overall similarity in DamOnly signal between the two NBs. Hierarchical clustering separates NBs, indicating condition-specificity despite overall similarity. **C. Chromatin accessibility is continuous across the landscape.** Histogram of mean log_2_ DamOnly distribution (averaged across both NBs) across all 27,119 regions shows a unimodal, right-skewed distribution. **D. Ranked bins represent increasing accessibility states.** DamOnly signal increases from bin 1 (least accessible) to bin 10 (most accessible) within the constructed accessibility spectrum. Each dot represents log_2_-transformed DamOnly signal in a given region within the bin, averaged across replicates, coloured blue for NB5-6 and red for NB7-4. Boxes show median and IQR. Genotypes for NB5-6 chromatin accessibility: *w-; Lbe-k-Gal4/+; +/UAS-DamOnly*. NB7-4 chromatin accessibility: *w-; 19B03[AD]/+; 18F07[DBD]/UAS-DamOnly*.

**Figure 3—Figure Supplement 1.**
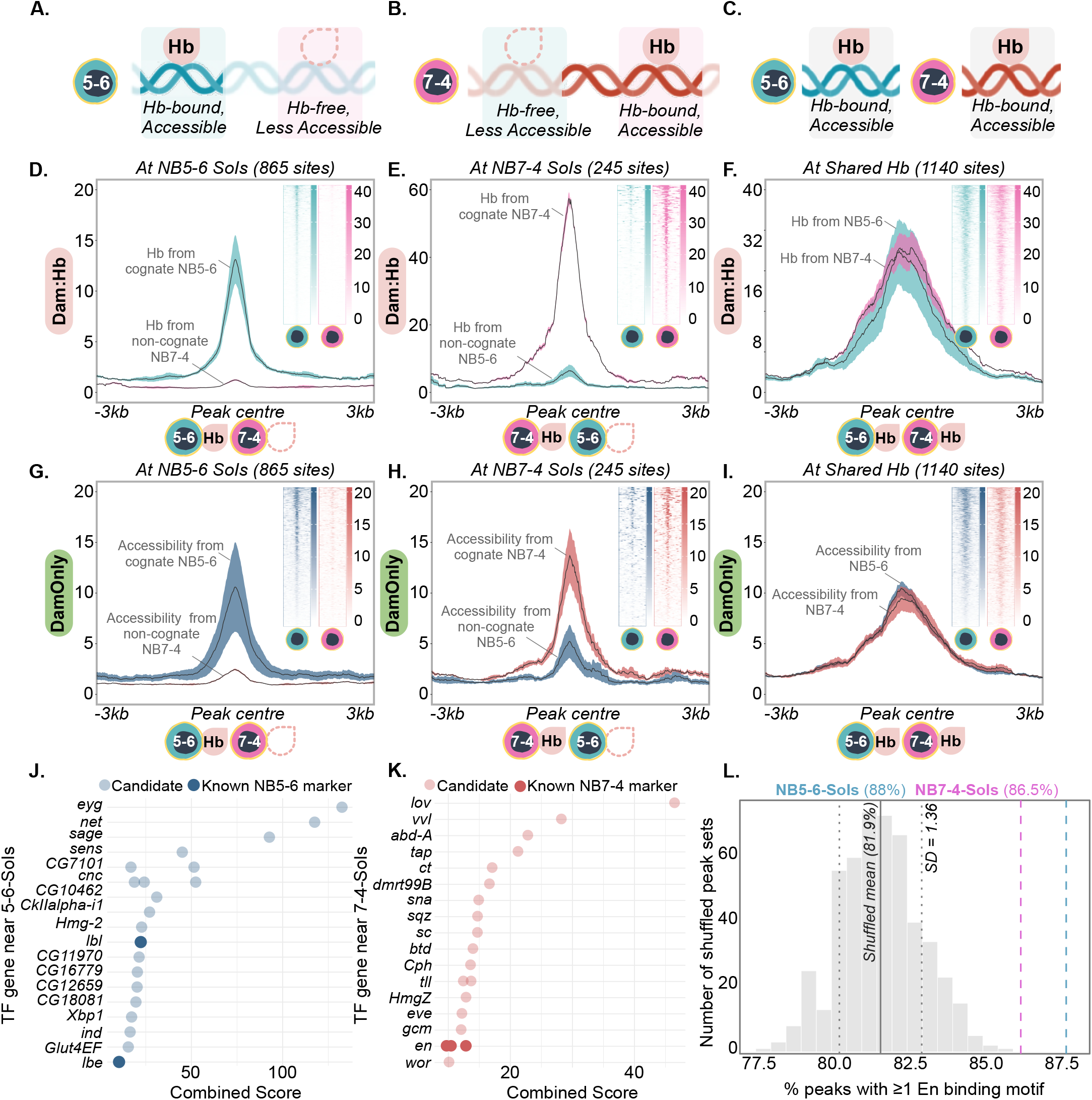
NB-specific sites of integration (SoIs) are differentially Hb-bound and differentially accessible. **A–C. Schematics defining NB-specific and shared Hb-bound sites.** NB5-6 SoIs (865 sites; shaded blue boxes in A and B) are defined as loci that are Hb-bound and accessible in NB5-6 (A), but Hb-free and less accessible in NB7-4 (B). NB7-4 SoIs (245 sites; shaded pink boxes in A and B) are Hb-bound and accessible in NB7-4 (A), while the corresponding loci in NB5-6 are Hb-free and less accessible (B). Sites identified by differential analysis of Hb occupancy in NB5-6 and NB7-4 (FDR ≤ 0.01, FC ≥ 2, see Methods). Shared Hb-bound sites (1,140 sites; grey shaded boxes) are Hb-bound and accessible in both NB5-6 and NB7-4 (C), identified by ≥ 50% peak overlap. **D–F. Hb occupancy validates NB-specific SoI definitions**. At NB5-6 SoIs (D), NB5-6 Hb signal (blue) exceeds NB7-4 Hb signal (pink). At NB7-4 SoIs (E), NB7-4 Hb signal exceeds NB5-6 Hb signal. At shared Hb-bound loci (*n* = 1,140), Hb signal is comparable between both NBs (F). Plots show mean Dam:Hb signal, *±* SD (shaded ribbon), *±*3kb from peak centre. Insets show per-site signal heatmaps, averaged across three replicates from each NB. **G–I. Chromatin accessibility validates NB-specific SoI definitions**. At NB5-6 SoIs, NB5-6 DamOnly signal (dark blue) exceeds NB7-4 DamOnly signal (red; G). At NB7-4 SoIs, NB7-4 DamOnly signal exceeds NB5-6 DamOnly signal (H). At shared Hb-bound sites, chromatin is similarly accessible in both NBs (I). Plots show mean DamOnly signal, *±* SD (shaded ribbon), *±*3kb from peak centre. Insets show per-site signal heatmaps, averaged across four replicates from NB5-6 and three from NB7-4. **J–K. Genes encoding lineage-specific TFs are recovered near SoIs**. Annotation of TF-encoding genes near NB5-6 SoIs recovers known NB5-6 markers including *lbe* and *lbl* (J). Annotation near NB7-4 SoIs recovers *en* (K). Known lineage markers (dark dots) are distinguished from candidates (light dots). Candidates were ranked by a combined score incorporating fold change of Hb occupancy, statistical confidence (− log_10_ FDR), and distance to the nearest TSS (see Methods). **L. Lack of En binding at NB5-6 SoIs despite presence of binding motif**. En binding motif occurrence is reported as the percentage of peaks containing at least one motif, using size- and composition-matched shuffled peaksets as background (mean 81.9%, SD = 1.36; see Methods). NB7-4 SoIs (86.5%; pink dashed line), which are shown to be En-bound, serve as a positive control. NB5-6 SoIs (88%; blue dashed line) show comparable motif occurrence to NB7-4 SoIs. Genotypes for NB5-6 Hb occupancy: *w-; Lbe-k-Gal4/+; +/UAS-Dam:Hb*. NB7-4 Hb occupancy: *w-; 19B03[AD]/+; 18F07[DBD]/UAS-Dam:Hb* (42). NB5-6 chromatin accessibility: *w-; Lbe-k-Gal4/+; +/UAS-DamOnly*. NB7-4 chromatin accessibility: *w-; 19B03[AD]/+; 18F07[DBD]/UAS-DamOnly*.

**Figure 3—Figure Supplement 2.**
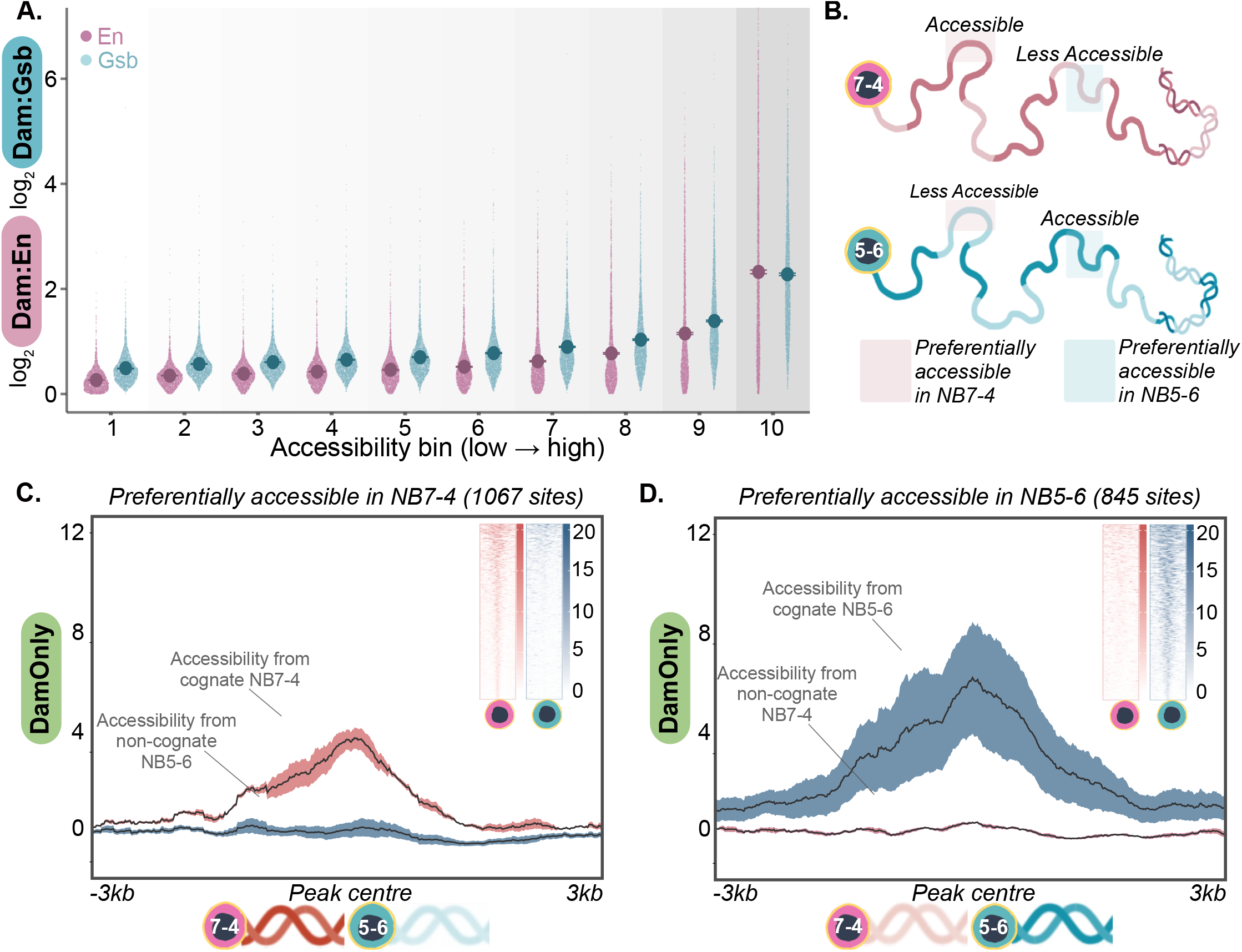
Validation of NB-specific accessible sites. **A. STF occupancy across the ranked accessibility bins.** Side-by-side sina plots show log2-transformed Dam:En (pink) and Dam:Gsb (blue) signal across ranked accessibility bins (see Methods, Figure 2—figure supplement 2), with mean ±SEM indicated per bin. Background shading indicates bin 1 (least accessible; lightest grey) to bin 10 (most accessible; darkest grey). Each point represents one genomic window (*n* ≈ 2,700 per bin; RPGC-normalised, averaged across three Dam:En and four Dam:Gsb replicates). **B. Defining accessible and less-accessible loci**. Schematic shows accessible and less-accessible loci in NB7-4 (top; dark/light red) and NB5-6 (bottom; dark/light blue), defined from the ranked accessibility bins at a 2-fold cutoff (see Methods, Figure 2G). NB7-4-preferential sites are accessible in NB7-4 but less accessible in NB5-6 (log2 ≤–1; 1067 sites; shaded pink boxes). NB5-6-preferential sites are accessible in NB5-6 but less accessible in NB7-4 (log2 ≥1; 845 sites; shaded blue boxes). **C–D. DamOnly signal validates NB-specific differentially accessible sites**. At NB7-4-preferential sites (C), DamOnly signal from NB7-4 (red) exceeds NB5-6 (blue); at NB5-6-preferential sites (D), NB5-6 signal exceeds NB7-4. Plots show mean DamOnly signal ±SD (shaded ribbon; red, NB7-4; blue, NB5-6), ±3kb from peak centre, averaged across replicates (three from NB7-4, four from NB5-6). Insets show per-site signal heatmaps at the corresponding loci, averaged across replicates. Genotypes for NB5-6-chromatin accessibility: *w-; Lbe-k-Gal4/+; +/UAS-DamOnly*. NB7-4-chromatin accessibility: *w-; 19B03[AD]/+; 18F07[DBD]/UAS-DamOnly*. NB5-6-Gsb occupancy: *w-; Lbe-k-Gal4/+; +/UAS-Dam:Gsb*. NB7-4-En occupancy: *w-; 19B03[AD]/+; 18F07[DBD]/UAS-Dam:En*.

**Figure 3—Figure Supplement 3.**
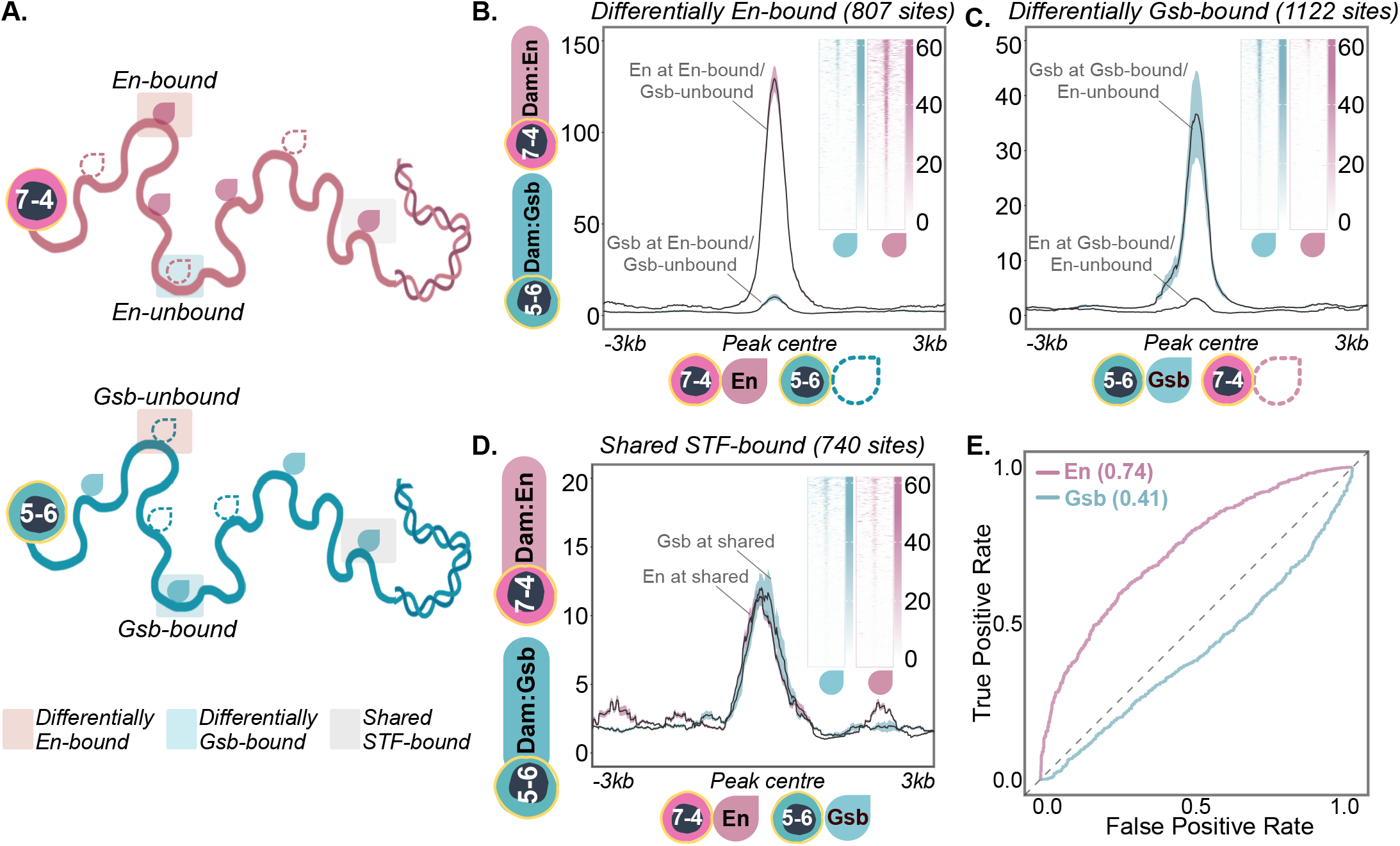
Validation of computationally determined STF-bound and -unbound sites. **A**. Schematic shows En-bound and -unbound sites in NB7-4 (top) and Gsb-bound and -unbound sites in NB5-6 (bottom), defined by differential STF occupancy analysis (FC ≥ 2, FDR ≤ 0.01; see Methods). Sites differentially bound by En are ‘En-bound’ in NB7-4 (filled pink drop) and ‘Gsb-unbound’ in NB5-6 (hollow blue drop), shown in shaded pink boxes. Sites differentially bound by Gsb are ‘Gsb-bound’ in NB5-6 (filled blue drop) and ‘En-unbound’ in NB7-4 (hollow pink drop), shown in shaded blue boxes. Sites co-bound by Gsb in NB5-6 and En in NB7-4, defined by ≥50% overlap between Dam:Gsb and Dam:En peaks, are shown in shaded grey boxes. **B–D. Dam:STF signal validates differential STF binding**. At En-bound/Gsb unbound sites (B; 807 sites), Dam:En signal (pink) exceeds Dam:Gsb (blue). At Gsb-bound/En-unbound sites (C; 1,122 sites), Dam:Gsb signal exceeds Dam:En. At co-bound sites (D; 740 sites), Dam:En and Dam:Gsb signal are comparable. Plots show mean Dam:STF signal *±*SD (shaded ribbon; pink, Dam:En; blue, Dam:Gsb), *±*3kb from peak centre, averaged across replicates (three Dam:En, four Dam:Gsb). Insets show per-site signal heatmaps at the corresponding loci, averaged across replicates. **E. Chromatin accessibility predicts En but not Gsb binding**. Receiver operating characteristic (ROC) curves for NB-specific accessibility classifying STF-bound versus -unbound sites: En (pink, AUC = 0.74), Gsb (blue, AUC = 0.41). Dashed line indicates chance (AUC = 0.5). Genotypes for NB5-6-Gsb occupancy: *w-; Lbe-k-Gal4/+; +/UAS-Dam:Gsb*. NB7-4-En occupancy: *w-; 19B03[AD]/+; 18F07[DBD]/UAS-Dam:En*. NB5-6-chromatin accessibility: *w-; Lbe-k-Gal4/+; +/UAS-DamOnly*. NB7-4-chromatin accessibility: *w-; 19B03[AD]/+; 18F07[DBD]/UAS-DamOnly*.

**Figure 6—Figure Supplement 1.**
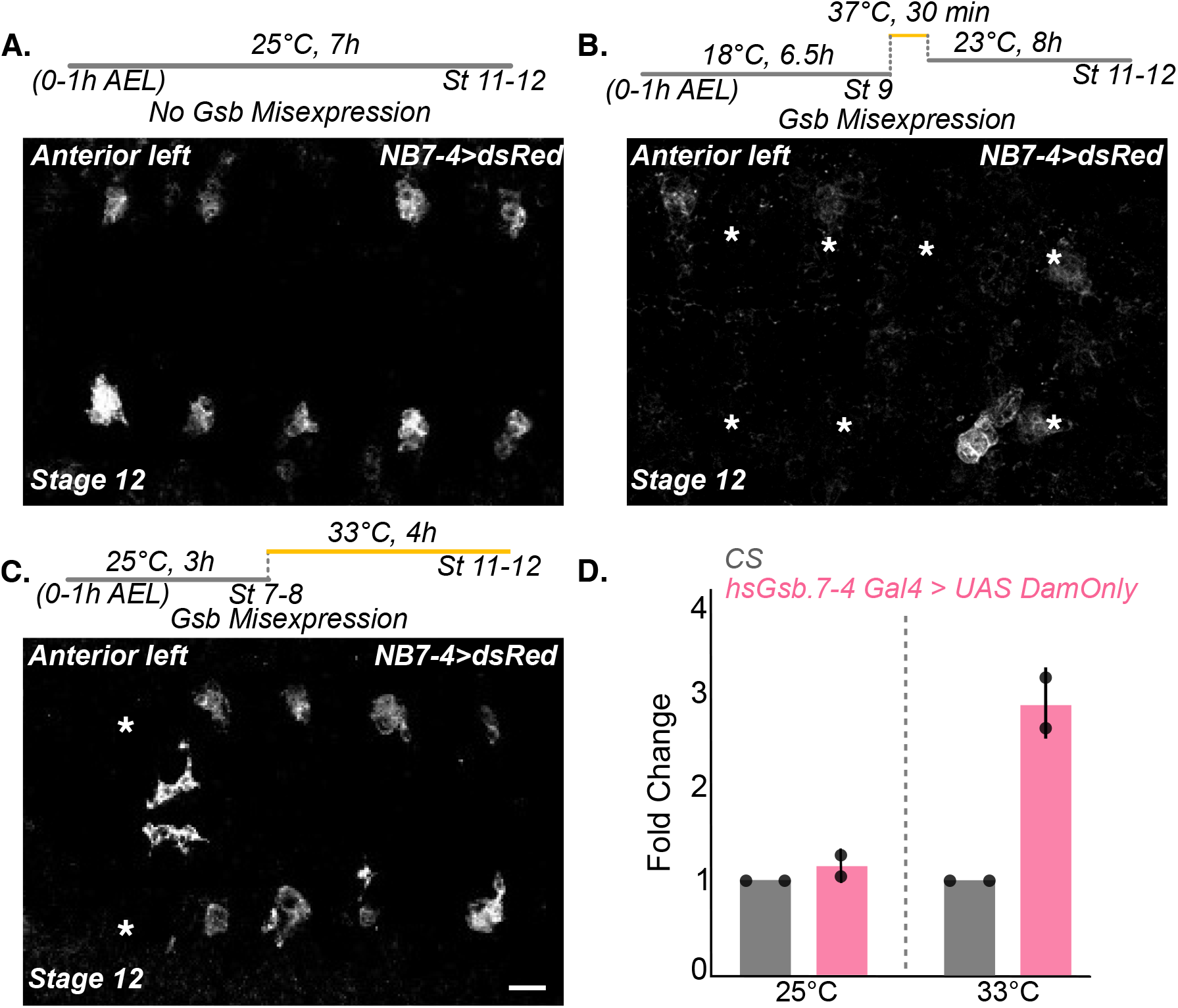
Validation of heat shock regime for Gsb misexpression and assaying accessibility and Hb occupancy from NB7-4. **A–C. Calibra tion of the heat-shock regime for Gsb misexpression in NB7-4.** Embryos of genotype *hsGsb; 19B03[AD]/UAS-mCD8-dsRed; 18F07[DBD]* were collected (0–1h AEL), and NB7-4*>*dsRed expression was assessed at Stage 12. In control embryos, incubated at 25^°^C throughout with no heat shock (A), NB7-4 lineages are consistently present. A heat shock at 37^°^C for 30min at Stage 9 disrupted NB7-4-Gal4 activity, with lineages absent in most hemisegments (asterisks, B). A calibrated, lower-temperature regime (33^°^C for 4h, from Stage 7–8 to Stage 11–12) maintained Gal4 activity comparably to control (C) and was used for assaying accessibility and Hb occupancy from NB7-4. White asterisks (B–C) mark missing NB7-4 lineages. Scale bar, 10 *µ*m. **D. Gsb transcripts are increased under the calibrated heat-shock regime**. qPCR confirms elevated Gsb transcript levels in *hsGsb; 19B03[AD]/+; 18F07[DBD]/UAS-DamOnly* embryos following the calibrated 33^°^C protocol (C) relative to wild-type CS embryos, but not at 25^°^C, confirming heat shock-dependent induction of ectopic Gsb while Gal4 activity is preserved. Bars show mean fold change; dots represent individual biological replicates (n = 2). Grey bars, wild-type CS; pink bars, heat-shocked *hsGsb* embryos. No-heat-shock (left) and heat-shocked (right) conditions are separated by the dashed line. Genotypes for NB7-4*>*dsRed (A–C): *hsGsb; 19B03[AD]/UAS-mCD8-dsRed; 18F07[DBD]*. Gsb transcript qPCR (D): *hsGsb; 19B03[AD]/+; 18F07[DBD]/UAS-DamOnly*; wild-type control, Canton-S.

**Figure 6—Figure Supplement 2.**
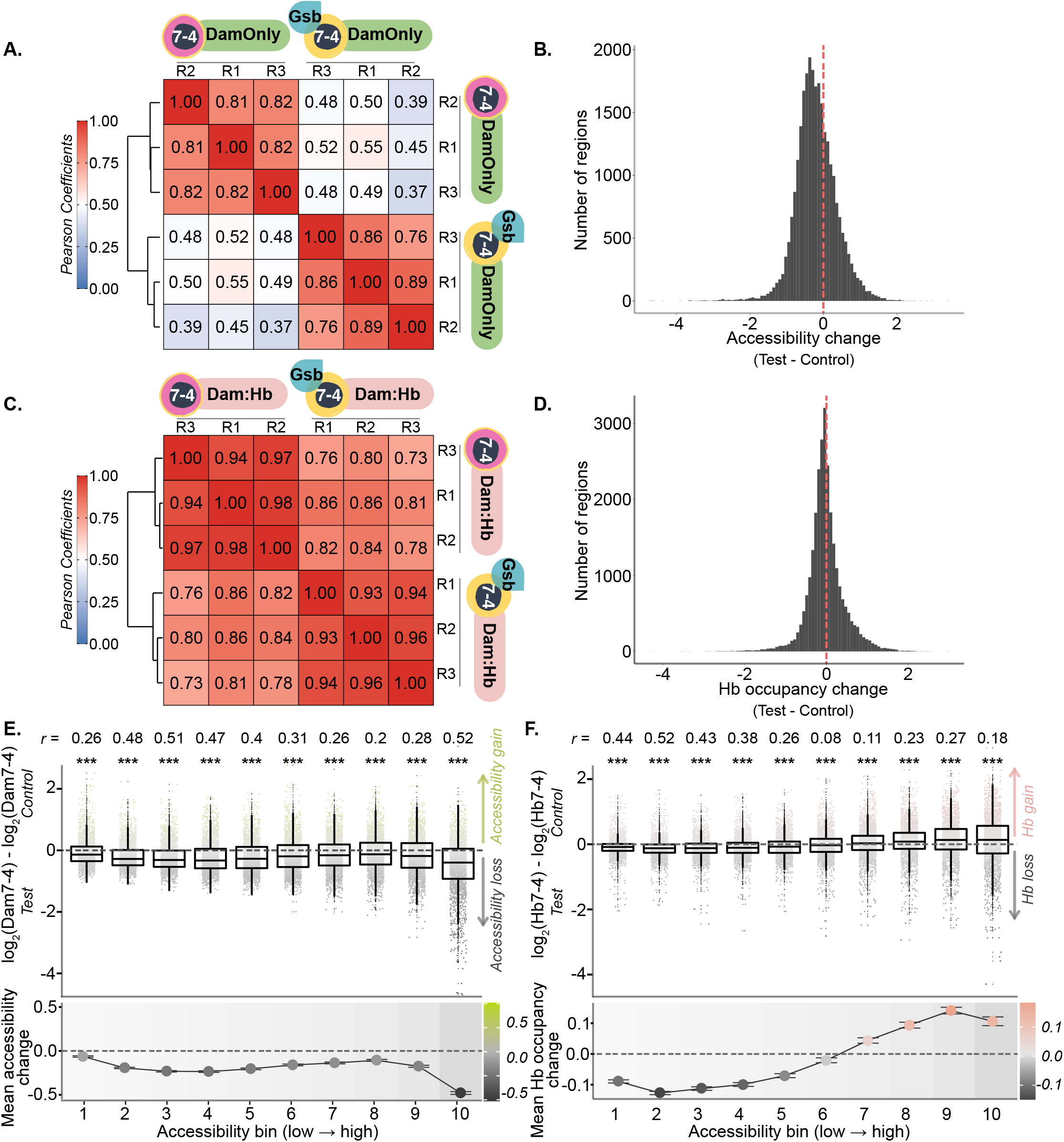
Genome-wide changes in accessibility and Hb occupancy upon Gsb misexpression. **A. Reproducibility of control and test DamOnly replicates across the accessibility landscape.** Pairwise Pearson correlation heatmap of control and Gsb-misexpressed (test) NB7-4 DamOnly replicates across all 27,119 regions of the accessibility landscape (see Methods, Figure 2—figure supplement 2). Hierarchical clustering separates control from test, with within-condition correlations exceeding between-condition correlations. **B. Bidirectional accessibility changes at subsets of loci**. Histogram of per-region accessibility change (log_2_ DamOnly, test−control) shows both losses (negative tail) and gains (positive tail). The distribution is skewed slightly towards loss. Red dashed line marks zero. **C. Reproducibility of control and test Dam:Hb replicates across the accessibility landscape**. Pairwise Pearson correlation heatmap of control and test NB7-4 Dam:Hb replicates across all 27,119 regions shows overall similarity in Dam:Hb signal between control and test. Hierarchical clustering separates conditions, despite overall similarity. **D. Bidirectional Hb occupancy changes at small subsets of loci**. Histogram of per-region Hb occupancy change (log_2_ Dam:Hb, test−control) shows both losses (negative tail) and gains (positive tail) at a small subset of loci. The distribution is centred near zero. Red dashed line marks zero. *(Figure 6—figure supplement 2 legend continued on next page*.) E. Accessibility gains and losses across the ranked accessibility spectrum. (Top) Chromatin accessibility (DamOnly signal) stratified into ten bins of increasing accessibility (see Methods, Figure 2—figure supplement 2). Accessibility change, expressed as log_2_fold-change in DamOnly signal from control to test NB7-4, is plotted across the ranked bins; positive values indicate gain (green), negative values loss (dark grey), and values near zero indicate unchanged accessibility (light grey). Each point represents one genomic window (*n* ≈ 2,700 per bin; RPGC-normalised, replicate-averaged), coloured by direction of change (green, gain; dark grey, loss; light grey, unchanged). Boxes show median and IQR. Black dashed line marks zero change. Green and grey dashed lines mark the 2-fold threshold for gain (log_2_ ≥ 1; 606 sites) and loss loci (log_2_ ≤ −1; 1,340 sites), respectively. Most regions show little change between control and test, with values distributed near zero but skewed negative towards loss across all bins. The spread of values indicates gains and losses at subsets of loci across all bins. Significance labels indicate that the median change per bin deviates from zero (one-sample Wilcoxon signed-rank test, two-sided; ∗ ∗ ∗*p <* 0.001, ∗ ∗ *p <* 0.01, ∗*p <* 0.05, *ns* = not significant), with effect size r shown above each bin. (Bottom) Mean accessibility change per bin shows the net effect of reduced accessibility upon Gsb misexpression. Background shading indicates bin 1 (least accessible; lightest grey) to bin 10 (most accessible; darkest grey). **F. Hb occupancy gains and losses across the ranked accessibility spectrum**. (Top) As in (E, Top), for Hb occupancy change (log_2_ Dam:Hb, test − control); positive values indicate Hb gain (pink), negative values Hb loss (dark grey), values near zero indicate unchanged Hb occupancy (light grey). (Bottom) As in (E, Bottom), mean change in Hb occupancy shows small losses in lower accessibility bins and gains in higher accessibility bins. Genotypes for NB7-4-Hb occupancy: *w-; 19B03[AD]/+; 18F07[DBD]/UAS-Dam:Hb* (42). NB7-4-chromatin accessibility: *w-; 19B03[AD]/+; 18F07[DBD]/UAS-DamOnly*. NB7-4-chromatin accessibility with Gsb misexpression: *hsGsb; 19B03[AD]/+; 18F07[DBD]/UAS-DamOnly*. NB7-4-Hb occupancy with Gsb misexpression: *hsGsb; 19B03[AD]/+; 18F07[DBD]/UAS-Dam:Hb*.

**Figure 7—Figure Supplement 1.**
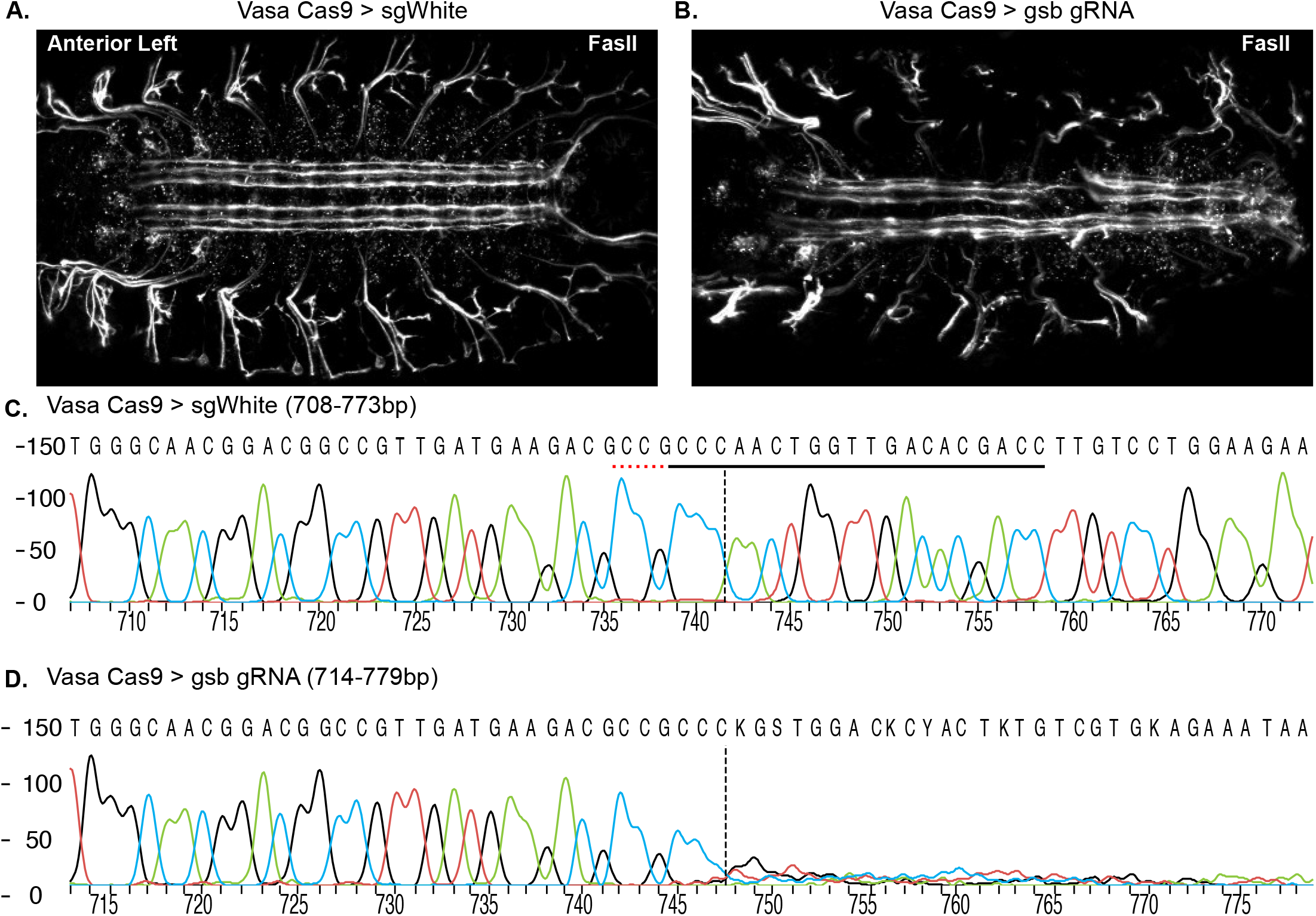
Validation of *gsb*-guideRNA. **A–B. FasII staining reveals axonal tract defects upon *gsb* editing.** In control embryos (Vasa-Cas9 *>* sgWhite), FasII staining shows intact axonal tracts (A). In test embryos (Vasa-Cas9 *> gsb*-gRNA), axonal tract defects are seen, consistent with loss of Gsb (B). **C–D. Sanger sequencing confirms Cas9-mediated editing at the *gsb* locus**. Sequencing chromatograms of control (Vasa-Cas9 *>* sgWhite; C) and edited (Vasa-Cas9 *> gsb*-gRNA; D) embryos were analysed using the Synthego ICE (Inference of CRISPR Edits) online tool. The predicted Cas9 cut site (dashed line) corresponds to position 747bp. The control trace shows clean base calls with no evidence of indels (C), whereas mixed peaks downstream of the cut site in the edited sample indicate indel formation at the *gsb* locus (D), consistent with efficient CRISPR editing (ICE-predicted efficiency, 75%). Genotypes for control embryos: *Vasa-Cas9/+; sgWhite/+*. For edited embryos: *Vasa-Cas9/y[1]v[1]; gsb-guideRNA/+*.

## Bibliography

1. Asif Bakshi, Khaled Ben El Kadhi, and Claude Desplan. Decoding neuronal diversity: Mechanisms governing neural cell fate in Drosophila. Current Opinion in Neurobiology, 93:103061, August 2025. ISSN 09594388. doi: 10.1016/j.conb.2025.103061.

2. Anthony M. Rossi, Shadi Jafari, and Claude Desplan. Integrated Patterning Programs During Drosophila Development Generate the Diversity of Neurons and Control Their Mature Properties. Annual Review of Neuroscience, 44(1):153–172, July 2021. ISSN 0147-006X, 1545-4126. doi: 10.1146/annurev-neuro-102120-014813.

3. Sonia Q. Sen. Generating neural diversity through spatial and temporal patterning. Seminars in Cell & Developmental Biology, 142:54–66, June 2023. ISSN 10849521. doi: 10.1016/j.semcdb.2022.06.002.

4. Aloisia Schmid, Akira Chiba, and Chris Q. Doe. Clonal analysis of Drosophila embryonic neuroblasts: Neural cell types, axon projections and muscle targets. Development, 126(21): 4653–4689, November 1999. ISSN 0950-1991, 1477-9129. doi: 10.1242/dev.126.21.4653.

5. Hartmut Schmidt, Christof Rickert, Torsten Bossing, Olaf Vef, Joachim Urban, and Gerhard M. Technau. The embryonic central nervous system lineages of Drosophila melanogaster. Developmental Biology, 189(2):186–204, 1997. ISSN 0012-1606. doi: 10.1006/dbio.1997.8660.

6. Haluk Lacin and James W Truman. Lineage mapping identifies molecular and architectural similarities between the larval and adult Drosophila central nervous system. eLife, 5: e13399, March 2016. ISSN 2050-084X. doi: 10.7554/eLife.13399.

7. Torsten Bossing, Gerald Udolph, Chris Q. Doe, and Gerhard M. Technau. The embryonic central nervous system lineages ofDrosophila melanogaster: I. Neuroblast lineages derived from the ventral half of the neuroectoderm. Developmental Biology, 179(1):41–64, 1996. ISSN 0012-1606. doi: 10.1006/dbio.1996.0240.

8. Matthias Landgraf, Torsten Bossing, Gerd M. Technau, and Michael Bate. The Origin, Location, and Projections of the Embryonic Abdominal Motorneurons of Drosophila. Journal of Neuroscience, 17(24):9642–9655, December 1997. ISSN 0270-6474, 1529-2401. doi: 10.1523/JNEUROSCI.17-24-09642.1997.

9. J. B. Skeath, G. Panganiban, J. Selegue, and S. B. Carroll. Gene regulation in two dimensions: The proneural achaete and scute genes are controlled by combinations of axis-patterning genes through a common intergenic control region. Genes & Development, 6 (12b):2606–2619, January 1992. ISSN 0890-9369, 1549-5477. doi: 10.1101/gad.6.12b.2606.

10. James B. Skeath, Yu Zhang, Robert Holmgren, Sean B. Carroll, and Chris Q. Doe. Specification of neuroblast identity in the Drosophila embryonic central nervous system by gooseberry-distal. Nature, 376(6539):427–430, August 1995. ISSN 1476-4687. doi: 10.1038/376427a0.

11. Quynh Chu-LaGraff and Chris Q. Doe. Neuroblast Specification and Formation Regulated by wingless in the Drosophila CNS. Science, 261(5128):1594–1597, September 1993. ISSN 0036-8075, 1095-9203. doi: 10.1126/science.8372355.

12. Martha J. Lundell, Quynh Chu-LaGraff, Chris Q. Doe, and Jay Hirsh. The engrailed and huckebein genes are essential for development of serotonin neurons in the Drosophila CNS. Molecular and Cellular Neuroscience, 7(1):46–61, 1996. ISSN 1044-7431. doi: 10.1006/mcne.1996.0004.

13. Nirupama Deshpande, Rainer Dittrich, Gerhard M. Technau, and Joachim Urban. Successive specification of Drosophila neuroblasts NB 6-4 and NB 7-3 depends on interaction of the segment polarity genes wingless, gooseberry and naked cuticle. Development, 128(17):3253–3261, September 2001. ISSN 1477-9129, 0950-1991. doi: 10.1242/dev.128.17.3253.

14. Nathan L.Q. Anderson, Sen Lin Lai, and Chris Q. Doe. Vnd and en are expressed in orthogonal stripes and act in a brief competence window to combinatorially specify nb7-1 and its early lineage. Developmental Biology, 532:73–82, 4 2026. ISSN 0012-1606. doi: 10.1016/J.YDBIO.2026.01.008.

15. Takako Isshiki, Masatoshi Takeichi, and Akinao Nose. The role of the msh homeobox gene during Drosophila neurogenesis: Implication for the dorsoventral specification of the neuroectoderm. Development, 124(16):3099–3109, August 1997. ISSN 0950-1991, 1477-9129. doi: 10.1242/dev.124.16.3099.

16. Joseph B. Weiss, Tonia Von Ohlen, Dervla M. Mellerick, Gregory Dressler, Chris Q. Doe, and Matthew P. Scott. Dorsoventral patterning in the Drosophila central nervous system: The intermediate neuroblasts defective homeobox gene specifies intermediate column identity. Genes & Development, 12(22):3591–3602, November 1998. ISSN 0890-9369, 1549-5477. doi: 10.1101/gad.12.22.3591.

17. Hsin Chu, Carlos Parras, Kalpana White, and Fernando Jiménez. Formation and specification of ventral neuroblasts is controlled by vnd in Drosophila neurogenesis. Genes & Development, 12(22):3613–3624, November 1998. ISSN 0890-9369, 1549-5477. doi: 10.1101/gad.12.22.3613.

18. Jocelyn A. McDonald and Chris Q. Doe. Establishing neuroblast-specific gene expression in the Drosophila CNS: Huckebein is activated by Wingless and Hedgehog and repressed by Engrailed and Gooseberry. Development, 124(5):1079–1087, March 1997. ISSN 0950-1991, 1477-9129. doi: 10.1242/dev.124.5.1079.

19. Yu Zhang and Xin Li. Development of the Drosophila Optic Lobe. Cold Spring Harbor Protocols, 2024(3):pdb.top108156, January 2024. ISSN 1940-3402, 1559-6095. doi: 10.1101/pdb.top108156.

20. Ted Erclik, Xin Li, Maximilien Courgeon, Claire Bertet, Zhenqing Chen, Ryan Baumert, June Ng, Clara Koo, Urfa Arain, Rudy Behnia, Alberto Del Valle Rodriguez, Lionel Senderowicz, Nicolas Negre, Kevin P. White, and Claude Desplan. Integration of temporal and spatial patterning generates neural diversity. Nature, 541(7637):365–370, January 2017. ISSN 0028-0836, 1476-4687. doi: 10.1038/nature20794.

21. Katrina S. Gold and Andrea H. Brand. Optix defines a neuroepithelial compartment in the optic lobe of the Drosophila brain. Neural Development, 9(1):18, July 2014. ISSN 1749-8104. doi: 10.1186/1749-8104-9-18.

22. Priscilla Valentino and Ted Erclik. Spalt and disco define the dorsal-ventral neuroepithelial compartments of the developing Drosophila medulla. Genetics, 222(3):iyac145, November 2022. ISSN 1943-2631. doi: 10.1093/genetics/iyac145.

23. Holger Apitz and Iris Salecker. Spatio-temporal relays control layer identity of direction-selective neuron subtypes in Drosophila. Nature Communications, 9(1):2295, June 2018. ISSN 2041-1723. doi: 10.1038/s41467-018-04592-z.

24. Chris Q. Doe. Temporal Patterning in the Drosophila CNS. Annual Review of Cell and Developmental Biology, 33:219–240, October 2017. ISSN 1530-8995. doi: 10.1146/annurev-cellbio-111315-125210.

25. Cédric Maurange. Temporal patterning in neural progenitors: From Drosophila development to childhood cancers. Disease Models & Mechanisms, 13(7):dmm044883, July 2020. ISSN 1754-8403. doi: 10.1242/dmm.044883.

26. Takako Isshiki, Bret Pearson, Scott Holbrook, and Chris Q. Doe. Drosophila Neuroblasts Sequentially Express Transcription Factors which Specify the Temporal Identity of Their Neuronal Progeny. Cell, 106(4):511–521, August 2001. ISSN 0092-8674, 1097-4172. doi: 10.1016/S0092-8674(01)00465-2.

27. Tanja Novotny, Regina Eiselt, and Joachim Urban. Hunchback is required for the specification of the early sublineage of neuroblast 7-3 in the Drosophila central nervous system. Development, 129(4):1027–1036, February 2002. ISSN 0950-1991. doi: 10.1242/dev.129.4.1027.

28. Thomas Brody and Ward F Odenwald. Programmed Transformations in Neuroblast Gene Expression during Drosophila CNS Lineage Development. Developmental Biology, 226(1): 34–44, October 2000. ISSN 0012-1606. doi: 10.1006/dbio.2000.9829.

29. Nikolaos Konstantinides, Katarina Kapuralin, Chaimaa Fadil, Luendreo Barboza, Rahul Satija, and Claude Desplan. Phenotypic Convergence: Distinct Transcription Factors Regulate Common Terminal Features. Cell, 174(3):622–635.e13, July 2018. ISSN 0092-8674. doi: 10.1016/j.cell.2018.05.021.

30. Takumi Suzuki, Masako Kaido, Rie Takayama, and Makoto Sato. A temporal mechanism that produces neuronal diversity in the Drosophila visual center. Developmental Biology, 380(1):12–24, August 2013. ISSN 0012-1606. doi: 10.1016/j.ydbio.2013.05.002.

31. Xin Li, Ted Erclik, Claire Bertet, Zhenqing Chen, Roumen Voutev, Srinidhi Venkatesh, Javier Morante, Arzu Celik, and Claude Desplan. Temporal patterning of Drosophila medulla neuroblasts controls neural fates. Nature, 498(7455):456–462, June 2013. ISSN 1476-4687. doi: 10.1038/nature12319.

32. Nikolaos Konstantinides, Isabel Holguera, Anthony M. Rossi, Aristides Escobar, Liébaut Dudragne, Yen-Chung Chen, Thinh N. Tran, Azalia M. Martínez Jaimes, Mehmet Neset Özel, Félix Simon, Zhiping Shao, Nadejda M. Tsankova, John F. Fullard, Uwe Walldorf, Panos Roussos, and Claude Desplan. A complete temporal transcription factor series in the fly visual system. Nature, 604(7905):316–322, April 2022. ISSN 1476-4687. doi: 10.1038/s41586-022-04564-w.

33. Hailun Zhu, Sihai Dave Zhao, Alokananda Ray, Yu Zhang, and Xin Li. A comprehensive temporal patterning gene network in Drosophila medulla neuroblasts revealed by single-cell RNA sequencing. Nature Communications, 13(1):1247, March 2022. ISSN 2041-1723. doi: 10.1038/s41467-022-28915-3.

34. Makoto Sato and Takumi Suzuki. Cutting edge technologies expose the temporal regulation of neurogenesis in the Drosophila nervous system. Fly, 16(1):222–232, December 2022. ISSN 1933-6934. doi: 10.1080/19336934.2022.2073158.

35. Eri Hasegawa, Masako Kaido, Rie Takayama, and Makoto Sato. Brain-specific-homeobox is required for the specification of neuronal types in the Drosophila optic lobe. Developmental Biology, 377(1):90–99, May 2013. ISSN 0012-1606. doi: 10.1016/j.ydbio.2013.02.012.

36. Eri Hasegawa, Yusuke Kitada, Masako Kaido, Rie Takayama, Takeshi Awasaki, Tetsuya Tabata, and Makoto Sato. Concentric zones, cell migration and neuronal circuits in the Drosophila visual center. Development, 138(5):983–993, March 2011. ISSN 0950-1991. doi: 10.1242/dev.058370.

37. Makoto I. Kanai, Masataka Okabe, and Yasushi Hiromi. Seven-up Controls Switching of Transcription Factors that Specify Temporal Identities of Drosophila Neuroblasts. Developmental Cell, 8(2):203–213, February 2005. ISSN 1534-5807. doi: 10.1016/j.devcel.2004.12.014.

38. Marta Moris-Sanz, Alicia Estacio-Gómez, Javier Álvarez-Rivero, and Fernando J. Díaz-Benjumea. Specification of neuronal subtypes by different levels of Hunchback. Development, 141(22):4366–4374, November 2014. ISSN 0950-1991. doi: 10.1242/dev.113381.

39. Julie Broadus, James B. Skeath, Eric P. Spana, Torsten Bossing, Gerhard Technau, and Chris Q. Doe. New neuroblast markers and the origin of the aCC/pCC neurons in the Drosophila central nervous system. Mechanisms of Development, 53(3):393–402, November 1995. ISSN 0925-4773. doi: 10.1016/0925-4773(95)00454-8.

40. Khoa D. Tran and Chris Q. Doe. Pdm and Castor close successive temporal identity windows in the NB3-1 lineage. Development (Cambridge, England), 135(21):3491–3499, November 2008. ISSN 0950-1991. doi: 10.1242/dev.024349.

41. Yen-Chung Chen and Nikolaos Konstantinides. Integration of spatial and temporal patterning in the invertebrate and vertebrate nervous system. Frontiers in Neuroscience, 16: 854422, 2022. ISSN 1662-453X. doi: 10.3389/fnins.2022.854422.

42. Sonia Q Sen, Sachin Chanchani, Tony D Southall, and Chris Q Doe. Neuroblast-specific open chromatin allows the temporal transcription factor, Hunchback, to bind neuroblast-specific loci. eLife, 8:e44036, January 2019. ISSN 2050-084X. doi: 10.7554/eLife.44036.

43. Tony D. Southall, Katrina S. Gold, Boris Egger, Catherine M. Davidson, Elizabeth E. Caygill, Owen J. Marshall, and Andrea H. Brand. Cell-Type-Specific Profiling of Gene Expression and Chromatin Binding without Cell Isolation: Assaying RNA Pol II Occupancy in Neural Stem Cells. Developmental Cell, 26(1):101–112, July 2013. ISSN 15345807. doi: 10.1016/j.devcel.2013.05.020.

44. Gabriel N Aughey, Alicia Estacio Gomez, Jamie Thomson, Hang Yin, and Tony D Southall. CATaDa reveals global remodelling of chromatin accessibility during stem cell differentiation in vivo. eLife, 7:e32341, February 2018. ISSN 2050-084X. doi: 10.7554/eLife.32341.

45. The modENCODE Consortium, Sushmita Roy, Jason Ernst, Peter V. Kharchenko, Pouya Kheradpour, Nicolas Negre, Matthew L. Eaton, Jane M. Landolin, Christopher A. Bristow, Lijia Ma, Michael F. Lin, Stefan Washietl, Bradley I. Arshinoff, Ferhat Ay, Patrick E. Meyer, Nicolas Robine, Nicole L. Washington, Luisa Di Stefano, Eugene Berezikov, Christopher D. Brown, Rogerio Candeias, Joseph W. Carlson, Adrian Carr, Irwin Jungreis, Daniel Mar-bach, Rachel Sealfon, Michael Y. Tolstorukov, Sebastian Will, Artyom A. Alekseyenko, Carlo Artieri, Benjamin W. Booth, Angela N. Brooks, Qi Dai, Carrie A. Davis, Michael O. Duff, Xin Feng, Andrey A. Gorchakov, Tingting Gu, Jorja G. Henikoff, Philipp Kapranov, Ren-hua Li, Heather K. MacAlpine, John Malone, Aki Minoda, Jared Nordman, Katsutomo Oka-mura, Marc Perry, Sara K. Powell, Nicole C. Riddle, Akiko Sakai, Anastasia Samsonova, Jeremy E. Sandler, Yuri B. Schwartz, Noa Sher, Rebecca Spokony, David Sturgill, Marijke van Baren, Kenneth H. Wan, Li Yang, Charles Yu, Elise Feingold, Peter Good, Mark Guyer, Rebecca Lowdon, Kami Ahmad, Justen Andrews, Bonnie Berger, Steven E. Brenner, Michael R. Brent, Lucy Cherbas, Sarah C. R. Elgin, Thomas R. Gingeras, Robert Grossman, Roger A. Hoskins, Thomas C. Kaufman, William Kent, Mitzi I. Kuroda, Terry Orr-Weaver, Norbert Perrimon, Vincenzo Pirrotta, James W. Posakony, Bing Ren, Steven Russell, Peter Cherbas, Brenton R. Graveley, Suzanna Lewis, Gos Micklem, Brian Oliver, Peter J. Park, Susan E. Celniker, Steven Henikoff, Gary H. Karpen, Eric C. Lai, David M. MacAlpine, Lincoln D. Stein, Kevin P. White, Manolis Kellis, David Acevedo, Richard Auburn, Galt Barber, Hugo J. Bellen, Eric P. Bishop, Terri D. Bryson, Aurelien Chateigner, Jia Chen, Hiram Clawson, Charles L. G. Comstock, Sergio Contrino, Leyna C. DeNapoli, Queying Ding, Alex Dobin, Marc H. Domanus, Jorg Drenkow, Sandrine Dudoit, Jackie Dumais, Thomas Eng, Delphine Fagegaltier, Sarah E. Gadel, Srinka Ghosh, Francois Guillier, David Hanley, Gregory J. Hannon, Kasper D. Hansen, Elizabeth Heinz, Angie S. Hinrichs, Martin Hirst, Sonali Jha, Lichun Jiang, Youngsook L. Jung, Helena Kashevsky, Cameron D. Kennedy, Ellen T. Kephart, Laura Langton, Ok-Kyung Lee, Sharon Li, Zirong Li, Wei Lin, Daniela Linder-Basso, Paul Lloyd, Rachel Lyne, Sarah E. Marchetti, Marco Marra, Nicolas R. Mattiuzzo, Sheldon McKay, Folker Meyer, David Miller, Steven W. Miller, Richard A. Moore, Carolyn A. Morrison, Joseph A. Prinz, Michelle Rooks, Richard Moore, Kim M. Rutherford, Peter Ruzanov, Douglas A. Scheftner, Lionel Senderowicz, Parantu K. Shah, Gregory Shanower, Richard Smith, E. O. Stinson, Sarah Suchy, Aaron E. Tenney, Feng Tian, Koen J. T. Venken, Huaien Wang, Robert White, Jared Wilkening, Aarron T. Willingham, Chris Zaleski, Zheng Zha, Dayu Zhang, Yongjun Zhao, and Jennifer Zieba. Identification of Functional Elements and Regulatory Circuits by Drosophila modENCODE. Science, 330(6012): 1787–1797, December 2010. doi: 10.1126/science.1198374.

46. S. Baumgartner, D. Bopp, M. Burri, and M. Noll. Structure of two genes at the gooseberry locus related to the paired gene and their spatial expression during Drosophila embryogenesis. Genes & Development, 1(10):1247–1267, January 1987. ISSN 0890-9369, 1549-5477. doi: 10.1101/gad.1.10.1247.

47. Nathalie Bonneaud, Sophie Layalle, Sophie Colomb, Christophe Jourdan, Alain Ghysen, Dany Severac, Christelle Dantec, Nicolas Nègre, and Florence Maschat. Control of nerve cord formation by Engrailed and Gooseberry-Neuro: A multi-step, coordinated process. Developmental Biology, 432(2):273–285, December 2017. ISSN 0012-1606. doi: 10.1016/j.ydbio.2017.10.018.

48. Andrew D. Renault. Vasa is expressed in somatic cells of the embryonic gonad in a sex-specific manner in Drosophila melanogaster. Biology Open, 1(10):1043–1048, August 2012. ISSN 2046-6390. doi: 10.1242/bio.20121909.

49. Magnus Baumgardt, Daniel Karlsson, Javier Terriente, Fernando J. Díaz-Benjumea, and Stefan Thor. Neuronal Subtype Specification within a Lineage by Opposing Temporal Feed-Forward Loops. Cell, 139(5):969–982, November 2009. ISSN 0092-8674. doi: 10.1016/j.cell.2009.10.032.

50. Hugo Gabilondo, Johannes Stratmann, Irene Rubio-Ferrera, Irene Millán-Crespo, Patricia Contero-García, Shahrzad Bahrampour, Stefan Thor, and Jonathan Benito-Sipos. Neuronal Cell Fate Specification by the Convergence of Different Spatiotemporal Cues on a Common Terminal Selector Cascade. PLOS Biology, 14(5):e1002450, May 2016. ISSN 1545-7885. doi: 10.1371/journal.pbio.1002450.

51. Gregory K. Davis, Joseph A. D’Alessio, and Nipam H. Patel. Pax3/7 genes reveal conservation and divergence in the arthropod segmentation hierarchy. Developmental Biology, 285 (1):169–184, September 2005. ISSN 0012-1606. doi: 10.1016/j.ydbio.2005.06.014.

52. L Xue and M Noll. The functional conservation of proteins in evolutionary alleles and the dominant role of enhancers in evolution. The EMBO Journal, 15(14):3722–3731, July 1996. ISSN 0261-4189.

53. Alexandre Mayran, Konstantin Khetchoumian, Fadi Hariri, Tomi Pastinen, Yves Gauthier, Aurelio Balsalobre, and Jacques Drouin. Pioneer factor Pax7 deploys a stable enhancer repertoire for specification of cell fate. Nature Genetics, 50(2):259–269, February 2018. ISSN 1546-1718. doi: 10.1038/s41588-017-0035-2.

54. Alexandre Mayran, Kevin Sochodolsky, Konstantin Khetchoumian, Juliette Harris, Yves Gauthier, Amandine Bemmo, Aurelio Balsalobre, and Jacques Drouin. Pioneer and non-pioneer factor cooperation drives lineage specific chromatin opening. Nature Communications, 10(1):3807, August 2019. ISSN 2041-1723. doi: 10.1038/s41467-019-11791-9.

55. Lionel Budry, Aurélio Balsalobre, Yves Gauthier, Konstantin Khetchoumian, Aurore L’Honoré, Sophie Vallette, Thierry Brue, Dominique Figarella-Branger, Björn Meij, and Jacques Drouin. The selector gene Pax7 dictates alternate pituitary cell fates through its pioneer action on chromatin remodeling. Genes & Development, 26(20):2299–2310, October 2012. ISSN 0890-9369. doi: 10.1101/gad.200436.112.

56. Elena N. Tolkunova, Miki Fujioka, Masatomo Kobayashi, Deepali Deka, and James B. Jaynes. Two Distinct Types of Repression Domain in Engrailed: One Interacts with the Groucho Corepressor and Is Preferentially Active on Integrated Target Genes. Molecular and Cellular Biology, 18(5):2804–2814, May 1998. ISSN 0270-7306. doi: 10.1128/mcb.18.5.2804.

57. Richard R. Copley. The EH1 motif in metazoan transcription factors. BMC Genomics, 6(1): 169, November 2005. ISSN 1471-2164. doi: 10.1186/1471-2164-6-169.

58. G. Chen, J. Fernandez, S. Mische, and A. J. Courey. A functional interaction between the histone deacetylase Rpd3 and the corepressor Groucho in Drosophila development. Genes & Development, 13(17):2218–2230, September 1999. ISSN 0890-9369. doi: 10.1101/gad.13.17.2218.

59. Aamna Kaul, Eugene Schuster, and Barbara H. Jennings. The Groucho Co-repressor Is Primarily Recruited to Local Target Sites in Active Chromatin to Attenuate Transcription. PLOS Genetics, 10(8):e1004595, August 2014. ISSN 1553-7404. doi: 10.1371/journal.pgen.1004595.

60. Lichao Luo, Chia Keng Siah, and Yu Cai. Engrailed acts with Nejire to control decapentaplegic expression in the Drosophila ovarian stem cell niche. Development, 144(18):3224– 3231, September 2017. ISSN 1477-9129, 0950-1991. doi: 10.1242/dev.145474.

61. Cyrille Alexandre and Jean-Paul Vincent. Requirements for transcriptional repression and activation by Engrailed in Drosophila embryos. Development, 130(4):729–739, February 2003. ISSN 0950-1991. doi: 10.1242/dev.00286.

62. Florence Maschat, Nuria Serrano, Neel B. Randsholt, and Gérard Géraud. Engrailed and polyhomeotic interactions are required to maintain the A/P boundary of the Drosophila developing wing. Development, 125(15):2771–2780, August 1998. ISSN 0950-1991. doi: 10.1242/dev.125.15.2771.

63. Chris Q. Doe. Molecular markers for identified neuroblasts and ganglion mother cells in the Drosophila central nervous system. Development, 116(4):855–863, 1992. doi: 10.1242/dev.116.4.855.

64. Rolf Urbach and Gerhard M. Technau. Neuroblast formation and patterning during early brain development in Drosophila. BioEssays, 26(7):739–751, July 2004. ISSN 0265-9247, 1521-1878. doi: 10.1002/bies.20062.

65. Sonia Sen, Silvia Biagini, Heinrich Reichert, and K. VijayRaghavan. Orthodenticle is required for the development of olfactory projection neurons and local interneurons in Drosophila. Biology Open, 3(8):711–717, July 2014. ISSN 2046-6390. doi: 10.1242/bio.20148524.

66. Takaki Komiyama and Liqun Luo. Intrinsic control of precise dendritic targeting by an ensemble of transcription factors. Current Biology, 17(3):278–285, 2007. doi: 10.1016/j.cub.2006.11.067.

67. Maria Lynn Spletter, Jian Liu, Justin Liu, Helen Su, Edward Giniger, Takaki Komiyama, Stephen R. Quake, and Liqun Luo. Lola regulates drosophila olfactory projection neuron identity and targeting specificity. Neural Development, 2:14, 2007. doi: 10.1186/1749-8104-2-14.

68. Robert Lichtneckert, Lionel Nobs, and Heinrich Reichert. empty spiracles is required for the development of olfactory projection neuron circuitry in drosophila. Development, 135(14): 2415–2424, 2008. doi: 10.1242/dev.022210.

69. Hongjie Li, Felix Horns, Bing Wu, Qijing Xie, Jiefu Li, Tongchao Li, David J. Luginbuhl, Stephen R. Quake, and Liqun Luo. Classifying drosophila olfactory projection neuron subtypes by single-cell rna sequencing. Cell, 171(5):1206–1220.e22, 2017. doi:10.1016/j.cell.2017.10.019.

70. Mehmet NesetÖzel, Claudia Skok Gibbs, Isabel Holguera, Mennah Soliman, Richard Bonneau, and Claude Desplan. Coordinated control of neuronal differentiation and wiring by sustained transcription factors. Science, 378(6626):eadd1884, 2022. doi: 10.1126/science.add1884.

71. Owen J. Marshall, Tony D. Southall, Seth W. Cheetham, and Andrea H. Brand. Cell-type-specific profiling of protein-dna interactions without cell isolation using targeted damid with next-generation sequencing. Nature Protocols, 11(9):1586–1598, 2016. doi: 10.1038/nprot.2016.084.

72. Thomas J.R. Frith, James Briscoe, and Giulia L.M. Boezio. Chapter Five - From signalling to form: The coordination of neural tube patterning. In Moises Mallo, editor, Vertebrate Pattern Formation, volume 159 of Current Topics in Developmental Biology, pages 168–231. Academic Press, 2024. doi: 10.1016/bs.ctdb.2023.11.004.

73. Andreas Sagner, Isabel Zhang, Thomas Watson, Jorge Lazaro, Manuela Melchionda, and James Briscoe. A shared transcriptional code orchestrates temporal patterning of the central nervous system. PLOS Biology, 19(11):e3001450, November 2021. ISSN 1545-7885. doi: 10.1371/journal.pbio.3001450.

74. Isabel Zhang, Giulia L. M. Boezio, Jake Cornwall-Scoones, Thomas Frith, Elizabeth Finnie, Junyi Luo, Ming Jiang, Michael Howell, Robin Lovell-Badge, Andreas Sagner, James Briscoe, and M. Joaquina Delás. The cis-regulatory logic integrating spatial and temporal patterning in the vertebrate neural tube. Developmental Cell, 0(0), July 2025. ISSN 1534-5807. doi: 10.1016/j.devcel.2025.06.029.

75. Markus Nevil, Tyler J. Gibson, Constantine Bartolutti, Anusha Iyengar, and Melissa M. Harrison. Establishment of chromatin accessibility by the conserved transcription factor Grainy head is developmentally regulated. Development, 147(5):dev185009, March 2020. ISSN 1477-9129, 0950-1991. doi: 10.1242/dev.185009.

76. Fillip Port, Hui-Min Chen, Tzumin Lee, and Simon L. Bullock. Optimized CRISPR/Cas tools for efficient germline and somatic genome engineering in Drosophila. Proceedings of the National Academy of Sciences, 111(29):E2967–E2976, July 2014. doi: 10.1073/pnas.1405500111.

77. Owen J. Marshall, Tony D. Southall, Seth W. Cheetham, and Andrea H. Brand. Cell-type-specific profiling of protein–DNA interactions without cell isolation using targeted DamID with next-generation sequencing. Nature Protocols, 11(9):1586–1598, September 2016. ISSN 1750-2799. doi: 10.1038/nprot.2016.084.

78. Ben Langmead and Steven L. Salzberg. Fast gapped-read alignment with Bowtie 2. Nature Methods, 9(4):357–359, April 2012. ISSN 1548-7105. doi: 10.1038/nmeth.1923.

79. Heng Li, Bob Handsaker, Alec Wysoker, Tim Fennell, Jue Ruan, Nils Homer, Gabor Marth, Goncalo Abecasis, Richard Durbin, and 1000 Genome Project Data Processing Subgroup. The Sequence Alignment/Map format and SAMtools. Bioinformatics, 25(16):2078–2079, August 2009. ISSN 1367-4803. doi: 10.1093/bioinformatics/btp352.

80. Marcus R. Breese and Yunlong Liu. NGSUtils: A software suite for analyzing and manipulating next-generation sequencing datasets. Bioinformatics, 29(4):494–496, February 2013. ISSN 1367-4803. doi: 10.1093/bioinformatics/bts731.

81. Fidel Ramírez, Devon P Ryan, Björn Grüning, Vivek Bhardwaj, Fabian Kilpert, Andreas S Richter, Steffen Heyne, Friederike Dündar, and Thomas Manke. deepTools2: A next generation web server for deep-sequencing data analysis. Nucleic Acids Research, 44(W1): W160–W165, July 2016. ISSN 0305-1048. doi: 10.1093/nar/gkw257.

82. modENCODE Consortium. Gsb ChIP-seq. GEO accession GSE256559, 2024. https://www.ncbi.nlm.nih.gov/geo/query/acc.cgi?acc=GSE256559.

83. modENCODE Consortium. Twist ChIP-seq. GEO accession GSE256656, 2024. https://www.ncbi.nlm.nih.gov/geo/query/acc.cgi?acc=GSE256656.

84. modENCODE Consortium. Bicoid ChIP-seq. GEO accession GSE257099, 2024. https://www.ncbi.nlm.nih.gov/geo/query/acc.cgi?acc=GSE257099.

85. Yong Zhang, Tao Liu, Clifford A. Meyer, Jérôme Eeckhoute, David S. Johnson, Bradley E. Bernstein, Chad Nusbaum, Richard M. Myers, Myles Brown, Wei Li, and X. Shirley Liu. Model-based Analysis of ChIP-Seq (MACS). Genome Biology, 9(9):R137, September 2008. ISSN 1474-760X. doi: 10.1186/gb-2008-9-9-r137.

86. Sven Heinz, Christopher Benner, Nathanael Spann, Eric Bertolino, Yin C. Lin, Peter Laslo, Jason X. Cheng, Cornelis Murre, Harinder Singh, and Christopher K. Glass. Simple combinations of lineage-determining transcription factors prime cis-regulatory elements required for macrophage and b cell identities. Molecular Cell, 38:576–589, 5 2010. ISSN 10972765. doi: 10.1016/j.molcel.2010.05.004.

87. Rory Stark and Gordon Brown. DiffBind: differential binding analysis of ChIP-Seq peak data, 2011. Bioconductor Vignette.

88. Caryn S. Ross-Innes, Rory Stark, Andrew E. Teschendorff, Kelly A. Holmes, H. Raza Ali, Mark J. Dunning, Gordon D. Brown, Ondrej Gojis, Ian O. Ellis, Andrew R. Green, Simak Ali, Suet-Feung Chin, Carlo Palmieri, Carlos Caldas, and Jason S. Carroll. Differential oestrogen receptor binding is associated with clinical outcome in breast cancer. Nature, 481(7381):389–393, January 2012. ISSN 1476-4687. doi: 10.1038/nature10730.

89. Lihua J. Zhu, Claude Gazin, Nathan D. Lawson, Hervé Pagès, Simon M. Lin, David S. Lapointe, and Michael R. Green. Chippeakanno: a bioconductor package to annotate chip-seq and chip-chip data. BMC bioinformatics, 11, 5 2010. ISSN 1471-2105. doi: 10.1186/1471-2105-11-237.

90. Steven M. Gallo, Long Li, Zihua Hu, and Marc S. Halfon. Redfly: a regulatory element database for drosophila. Bioinformatics, 22:381–383, 2 2006. ISSN 1367-4803. doi: 10.1093/BIOINFORMATICS/BTI794.

91. Victoria K. Jenkins, Aoife Larkin, and Jim Thurmond. Using flybase: A database of drosophila genes and genetics. Methods in Molecular Biology, 2540:1–34, 2022. ISSN 19406029. doi: 10.1007/978-1-0716-2541-5_1/SAVE-RESEARCH.

92. Charles E. Grant, Timothy L. Bailey, and William Stafford Noble. Fimo: scanning for occurrences of a given motif. Bioinformatics, 27:1017–1018, 4 2011. ISSN 1367-4803. doi: 10.1093/BIOINFORMATICS/BTR064.

93. Damla Ovek Baydar, Ieva Rauluseviciute, Dina R Aronsen, Romain Blanc-Mathieu, Ine Bon-thuis, Herman de Beukelaer, Katalin Ferenc, Alice Jegou, Vipin Kumar, Roza Berhanu Lemma, Jérémy Lucas, Mathis Pochon, Chang M Yun, Vivekanandan Ramalingam, Salil Sanjay Deshpande, Aman Patel, Georgi K Marinov, Austin T Wang, Alejandro Aguirre, Jaime A Castro-Mondragon, Damir Baranasic, Jeanne Chèneby, Sveinung Gundersen, Morten Johansen, Aziz Khan, Marieke L Kuijjer, Eivind Hovig, Boris Lenhard, Albin Sandelin, Klaas Vandepoele, Wyeth W Wasserman, François Parcy, Anshul Kundaje, and Anthony Mathelier. Jaspar 2026: expansion of transcription factor binding profiles and integration of deep learning models. Nucleic Acids Research, 54:D184–D193, 1 2026. ISSN 0305-1048. doi: 10.1093/NAR/GKAF1209.

94. James T. Robinson, Helga Thorvaldsdóttir, Wendy Winckler, Mitchell Guttman, Eric S. Lander, Gad Getz, and Jill P. Mesirov. Integrative genomics viewer. Nature Biotechnology, 29 (1):24–26, January 2011. ISSN 1546-1696. doi: 10.1038/nbt.1754.

95. Helga Thorvaldsdóttir, James T. Robinson, and Jill P. Mesirov. Integrative Genomics Viewer (IGV): High-performance genomics data visualization and exploration. Briefings in Bioinformatics, 14(2):178–192, March 2013. ISSN 1467-5463. doi: 10.1093/bib/bbs017.

96. Johannes Schindelin, Ignacio Arganda-Carreras, Erwin Frise, Verena Kaynig, Mark Longair, Tobias Pietzsch, Stephan Preibisch, Curtis Rueden, Stephan Saalfeld, Benjamin Schmid, Jean-Yves Tinevez, Daniel James White, Volker Hartenstein, Kevin Eliceiri, Pavel Tomancak, and Albert Cardona. Fiji: An open-source platform for biological-image analysis. Nature Methods, 9(7):676–682, July 2012. ISSN 1548-7105. doi: 10.1038/nmeth.2019.

